# Revisiting Stress Granule Transcriptomes Suggests Mitochondrial RNA Enrichment Despite Methodological Bias

**DOI:** 10.64898/2026.09.08.749846

**Authors:** Alexandre Kaizeler, Diogo Coutinho, Diana V. Vieira, Vanessa A. Morais, Nuno L. Barbosa-Morais

## Abstract

Stress granules (SGs) are dynamic, membraneless cytoplasmic condensates that form in response to diverse cellular stressors. Although proposed to modulate stress responses by selectively sequestering proteins and RNAs, their precise molecular composition and function remain unclear. Reported SG transcriptomes differ substantially due to methodological discrepancies, notably between differential centrifugation (DC) and proximity labeling (PL). Here, we reanalyse publicly available human SG transcriptomes across multiple stressors, cell types, and isolation strategies. DC-based profiles were strongly shaped by RNA length, consistent with a physical bias inherent to the sedimentation-based separation. Correcting for this effect reveals limited concordance between studies. Mitochondrially encoded RNAs nonetheless consistently stand out as a distinctively regulated transcript class, without a uniform direction of enrichment/depletion across datasets, and with a modest but significant enrichment in our consensus SG signature. Immunofluorescence experiments further support, without definitively demonstrating, sequestration of mitochondrial dsRNA by SGs upon stress. Since leakage of mitochondrial nucleic acids is a well-characterised damage-associated molecular pattern linked to inflammation, their sequestration in SGs suggests a potential role in modulating immune-related stress responses. These findings refine our understanding of SG composition, underscore the need to control for technical artifacts in SG isolation, and provide a framework to distinguish genuine biological signals from methodological noise in SG research.

**Graphical Abstract:** 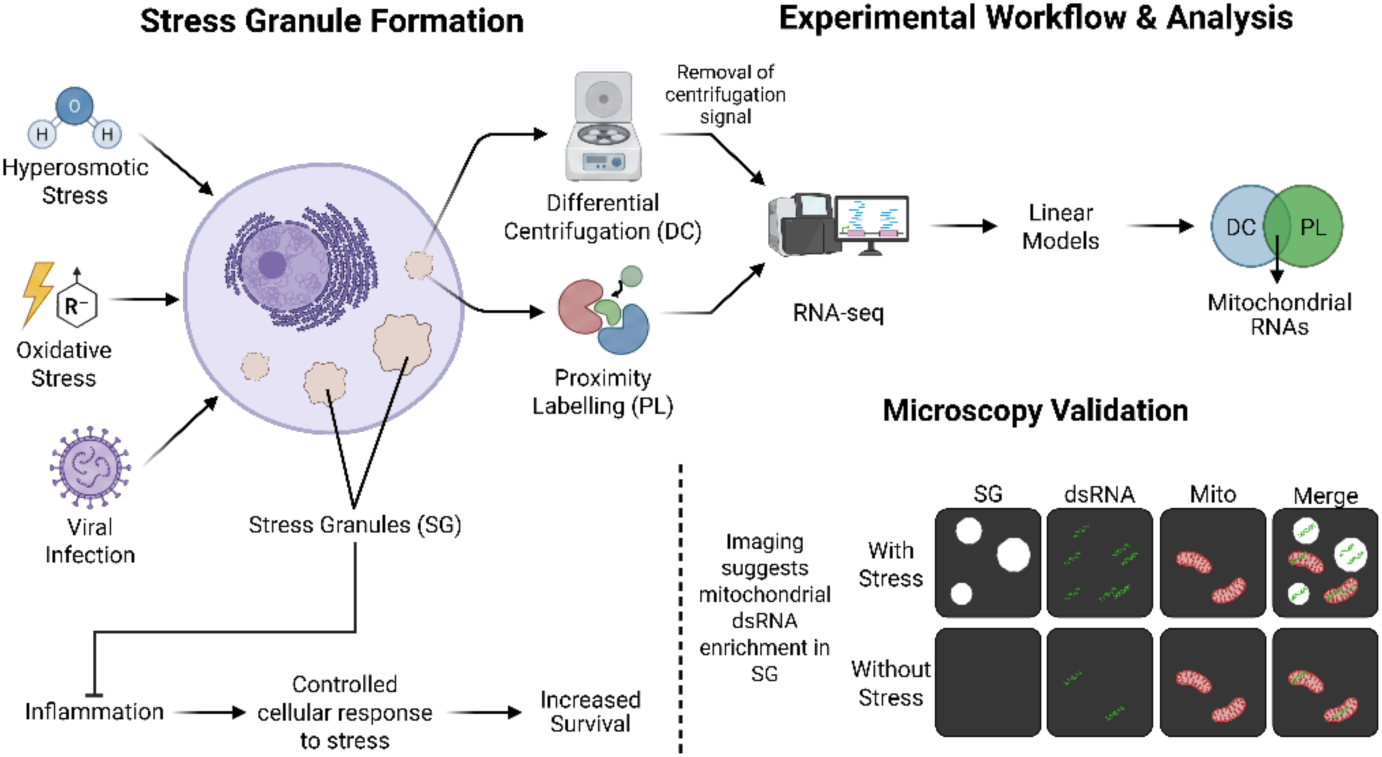

## Introduction

Stress granules (SGs) are membraneless cytoplasmic structures formed in response to a wide variety of stimuli(1), including hypoxia, nutrient deprivation, viral infection, T cell activation, heat-shock and oxidative stress(2–7). SGs comprise aggregations of RNA, stalled preinitiation complexes, and proteins(3), of which around half are RNA-binding proteins (RBPs)(8). Characterised by rapid formation and swift disassembly upon cessation of the triggering stimuli (15 and 30 minutes with oxidative and heat-shock stress, respectively)(9), SGs are highly dynamic structures, with their protein constituents continuously transitioning into and out of them(10). While SGs were first suggested to play a pivotal role in the global suppression of translation concurrent with cellular stress, they are now theorised to selectively sequester specific constituents, as stressed SG-deficient cells still exhibit repressed translation(11). As such, they may modulate and fine-tune the cellular response to environmental and biotic stress through sequestration of particular proteins and mRNAs: pro-apoptotic factors such as active caspases(12), or DDX3X to impede pyroptosis, thereby improving cell survival(13); *RB1*, protecting it from degradation in an RBFOX2-dependent manner(14); and PAI-1, hampering cell senescence(15). Notably, senescent cells actively deplete core SG components such as G3BP1 and TIA1, and G3BP1 itself is required for expression of the senescence-associated secretory phenotype (SASP) but not for the senescent state per se(16, 17), pointing to a dynamic interplay between SG assembly and senescence programmes. Nevertheless, the biological significance of these mechanisms remains ambiguous, with the function of the sequestered mRNAs uncertain, as studies have demonstrated their continued translation within the SG(18).

In addition to those physiological roles, SGs have been implicated in several pathologies. Mutations in certain SG-localised proteins, such as TIA1, TDP43 or ATXN2, are linked with amyotrophic lateral sclerosis(19–21), and certain mutations in DDX3X are suggested to originate stress granule-like assemblies that impair neurogenesis(22). Still, these results may be spurious, as both SGs and the physiopathology of these diseases rely on protein aggregation. SGs have also been implicated in antiviral responses. Viruses across essentially all major classes have evolved mechanisms to inhibit SG assembly, most commonly by targeting G3BP1 or upstream eIF2α signalling(23, 24). While this convergence suggests a genuine antiviral function, many of these strategies act on the integrated stress response (ISR) as a whole or on G3BP1 functions beyond SGs, making it difficult to attribute the antiviral effect to SGs specifically(23, 24). Cancer cells experience heightened stress due to factors such as hypoxia, nutrient deprivation, and chemotherapy/radiotherapy, all of which are reported to induce SG formation(2, 25, 26). SGs are observed in multiple tumours where they are postulated to promote metastasis while conferring resistance to chemotherapy, as their abrogation through G3BP1 and COX-1/2 inhibition impairs these phenotypes(27, 28). However, it is not known if these results solely arise from the blockade of SG formation or involve other functions of the targeted proteins, such as G3BP1 and COX-1/2, beyond their role in SGs. For example, only 20% of G3BP1 localises to SGs, reiterating how these targets may be nonspecific for SG blockage(29).

Despite the absence of direct evidence for their functions, SGs continue to garner significant attention in various physiological and pathological contexts, owing to their supposed role in selective mRNA and protein sequestration, hypothesised to contribute to the downregulation of specific pathways in numerous diseases. Research efforts have led to the curation of two databases housing information on their protein constituents(8, 30), revealing that RNA-binding proteins are their main component (>50%). Two main methodologies have emerged for elucidating their RNA content: differential centrifugation(31) (DC), and proximity RNA labelling via APEX-Seq(29) (PL), both relying on core SG proteins (usually G3BP1). Regardless, even when the same SG core protein and stress induction were used, notable disparities between transcriptomes isolated with the different techniques were observed(29, 31). DC highlights RNA length as a significant predictor of SG enrichment(31), but this effect is absent with PL(29). The enrichment in longer RNAs has been considered to have a biological nature, as SGs are formed by aggregation, longer transcripts equate to more potential binding sites for RBPs(31). Moreover, it is plausible that PL also captures G3BP1 that does not localise to SGs(29), and that therefore part of the obtained enrichment is not SG-specific. Nevertheless, concerns have been raised regarding the inherent bias of DC towards isolating longer transcripts, as longer RNAs tend to precipitate more readily given their greater number of intermolecular interaction sites, forming condensates with increased molecular weight(32, 33).

To resolve discrepancies in SG characterisation and identify authentic transcript features and their functional implications, we performed an unbiased analysis of publicly available human SG transcriptomic datasets spanning multiple cell lines, purification methods, and stressors. We found that transcriptomic profiles of SGs purified by differential centrifugation are strongly shaped by the method itself. Even after correcting for this bias, substantial differences between studies remained. Mitochondrial encoded RNAs nonetheless stood out as a distinctively regulated transcript class, with a modest but significant enrichment in our consensus SG signature despite inconsistent enrichment or depletion across individual datasets.

Our analyses further suggest that SGs may modulate the cellular stress response, potentially dampening rather than amplifying it, which could help limit downstream effects such as excessive inflammation. Although the functional significance of mitochondrial RNA enrichment is not yet resolved, immunofluorescence experiments suggest that mitochondrial dsRNAs are taken up by SGs upon stress, supporting a potential role in modulating immune signalling.

## Materials and Methods

### Reagents

| Reagent | Source | Identifier |
| --- | --- | --- |
| **Antibodies** |  |  |
| Rabbit Anti-G3BP1 | Atlas Antibodies | Cat # HPA004052,<br>RRID:AB_1849348 |
| Rabbit Anti-TIA1 | Atlas Antibodies | Cat # HPA056961,<br>RRID:AB_2683289 |
| Mouse Anti-dsRNA J2 | Jena Bioscience | Cat # RNT-SCI-10010200,<br>RRID:AB_2922431 |
| Chicken Anti-HSP60 | Bio-Techne Sales Corp. | Cat # NBP3-05536-50ul,<br>RRID:AB_3086708 |
| Donkey anti-Rabbit IgG (H+L) Highly Cross-Adsorbed Secondary Antibody, Alexa Fluor™ 647 | Thermo Fisher | Cat # A-31573,<br>RRID:AB_2536183 |
| Donkey anti-Mouse IgG (H+L) Highly Cross-Adsorbed Secondary Antibody, Alexa Fluor™ 488 | Thermo Fisher | Cat # A-21202,<br>RRID:AB_141607 |
| Donkey anti-Chicken IgY (H+L) Highly Cross Adsorbed Secondary Antibody, Alexa Fluor™ 568 | Thermo Fisher | Cat # A78950,<br>RRID:AB_2921072 |
| **Chemicals, dyes and other reagents** |  |  |
| DAPI | Sigma-Aldrich | Cat # D9452 |
| D-Sorbitol | Sigma-Aldrich | Cat # S6021 |
| DIDS | Sigma-Aldrich | Cat # 309795 |
| Triton X-100 | Sigma-Aldrich | Cat # T8787 |
| Bovine Serum Albumin | Sigma-Aldrich | Cat # A6003 |
| Magnesium chloride hexahydrate | Sigma-Aldrich | Cas # 7791-18-6 |
| Calcium chloride dihydrate | Sigma-Aldrich | Cas # 10035-04-8 |
| Goat Serum | Agilent Dako | Cat # X090710 |
| DMEM/F-12, no phenol red | Gibco | Cat # 21041025 |
| Fetal Bovine Serum, one shot | Gibco | Cat # A5209401 |
| Paraformaldehyde 16% Aqueous Solution EM Grade | Electron Microscopy Sciences | Cat # 15710 |
| Gelatine | VWR | Cat # 9764 |
| Costar® 24-well Clear TC-treated Multiple Well Plates | Corning | Cat # 3524 |
| VECTASHIELD® Antifade Mounting Medium | Vector Laboratories | Cat # H-1000 |
| VWR®, Microscope Slides | Avantor | Cat # 564378 |
| Cover Glasses Circular | Marienfeld Superior | Cat # 0111530 |

### Biological Resources

| Cell Lines | Source | Identifier |
| --- | --- | --- |
| U2-OS (Tet-On) | Clontech/Takara Bio | Cat # 630919,<br>RRID:CVCL_V335 |

### Software and Algorithms

| Resource | Source | Identifier |
| --- | --- | --- |
| FIJI (v1.54p) | NIH | ImageJ |
| ZEN (v3.12; in LSM 980) | Zeiss | ZEISS ZEN Microscopy Software |
| ZEN (v2.3 SP1 (black edition); in LSM 880) | Zeiss | ZEISS ZEN Microscopy Software |
| Imaris (v10.2.0) | Oxford Instruments | Imaris - Oxford Instruments |
| Microsoft PowerPoint (v16.0.1) | Microsoft | PowerPoint \| Microsoft 365 |
| R (v4.1.2) | R Foundation | R: The R Project for Statistical Computing |
| Perl (v5.26.1) | The Perl Foundation | The Perl Programming Language |
| Bowtie (v.1.2.2) | B. Langmead et al., 2009(34) | Bowtie: An ultrafast, memory-efficient short read aligner |
| STAR (v2.7.9a) | A. Dobin et al., 2013(35) | alexdobin/STAR: RNA-seq aligner |
| Trimmomatic (v0.3.9) | AM. Bolger et al., 2014(36) | USADELLAB.org - Trimmomatic: A flexible read trimming tool for Illumina NGS data |
| featureCounts (v2.0.3) | Y. Liao et al., 2014(37) | The Subread package |
| VAST-TOOLS (v2.5.1) | J. Tapial et al., 2017(38) | vastgroup/vast-tools: A toolset for profiling alternative splicing events in RNA-Seq data. |

### Datasets

For dataset procurement, we searched GEO Datasets using the term *“stress granule”*, filtering for expression profiling by high-throughput sequencing and restricting to human samples, excluding microarray studies. Using this approach, no single-cell datasets were retrieved. All publicly available studies up to 1 September 2024, were included. From the studies initially considered during the screening, we specifically excluded the dataset associated with Yi et al. (GSE245209 and GSE245210)(39), given that its CoCLIP method required peak-based analyses across conditions, more analogous to CLIP than standard differential expression. Part of the study from Sato et al. (GSE188397)(40) also featured samples more akin to CLIP, but, as these had considerably higher read depth, we found they could be analysed as regular RNA-Seq samples, and were thus kept.

The following datasets were used throughout this work (see Table 1; Figure 1B):

- For the SG transcriptomes: GSE99304(31); GSE138988(41); GSE121575(42); GSE138058(29); GSE119977(43, 44); GSE143413(45); GSE171792(46); GSE188397(40); GSE171655(47); GSE197001(6); GSE223295(48); GSE226161(49); GSE218180(50); GSE212380(51); GSE242766(52); PRJNA1152621(53).
- For the WC transcriptomes: GSE138988(41); GSE119977(43, 44); GSE173953(7).
- For the PB transcriptomes: GSE138988(41); GSE224858/GSE224752(54).
- The dataset for the yeast centrifugation modulation: GSE265963(33).
- The MAVS KD (senescence inhibition) dataset: GSE236521(55).

**Figure 1.**
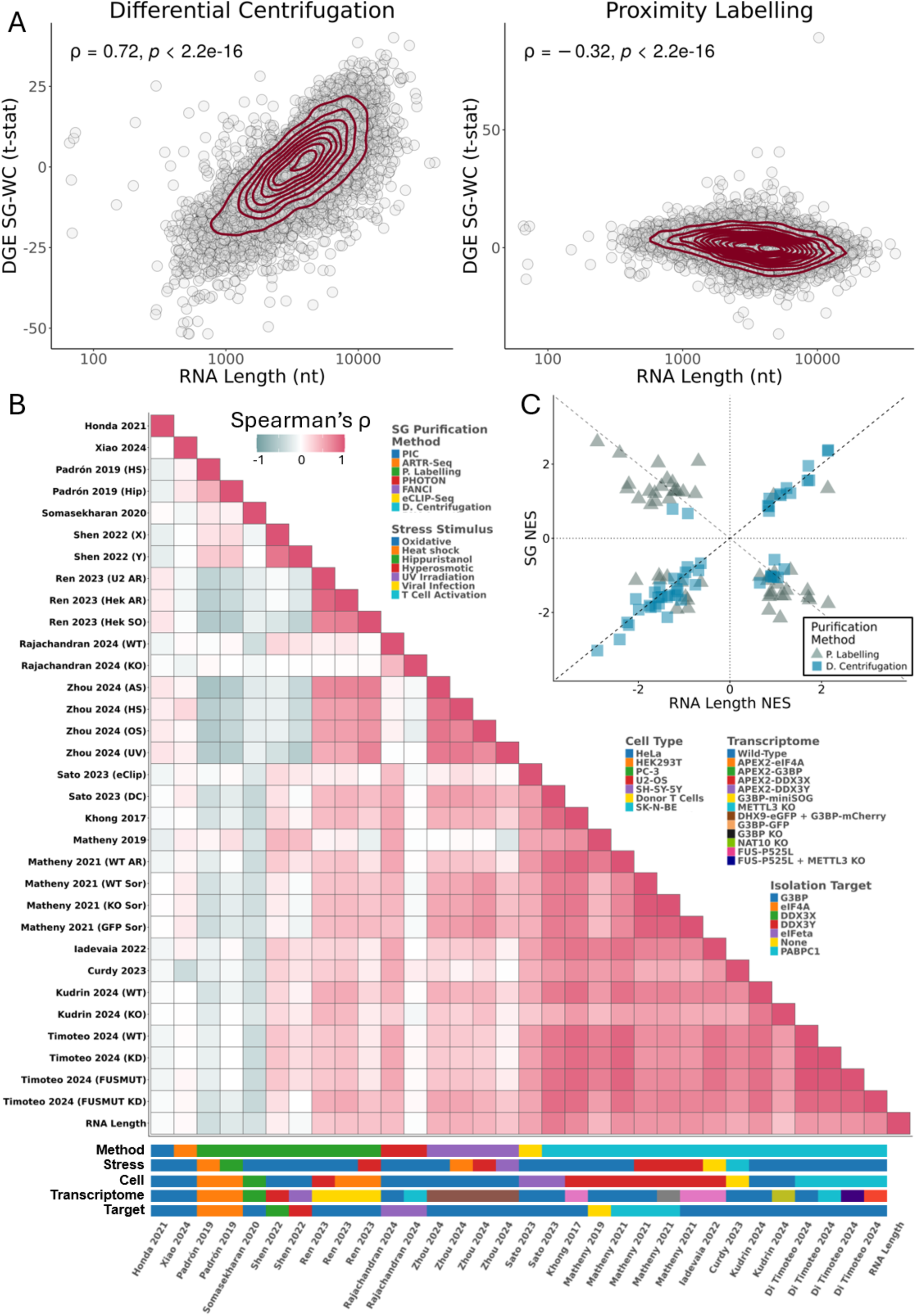
SGs purified by different methods have distinct transcriptomes. (A) Scatter plots of t-statistics of differential gene expression (DGE) between SGs and WCs (Y-axis) *versus* length of gene’s representative RNA transcript in number of nucleotides (nt) (X-axis), for SG purification by DC(31) (left) and PL(29) (right); genes as circles, density contour lines in dark red, Spearman’s correlation coefficient ρ and associated test’s p-value on top left corner. (B) Heatmap of Spearman’s correlations between t-statistics of DGE between SGs and WCs, from different studies, and RNA length. Stress stimulus, cell type, transcriptome of origin, SG purification method, and target molecule used for purification of each study are given below the heatmap. G3BP refers to G3BP1 alone or both G3BP1 and G3BP2. METTL3 knockdown and knockout are marked with the same colour. (C) Scatter plot of GSEA Normalised Enrichment Scores (NES) of Hallmarks between SGs and WCs, based on genes ranked by t-statistics of differential expression, for SG purification by DC(31) or PL(29) (SG NES - Y-axis) *versus* genes ranked by length of representative transcript (RNA Length NES - X-axis). Dashed lines depict identity (Y = X; Y = -X), and axis lines.

**Table 1.** Gene Expression Omnibus (GEO) datasets used for human SG transcriptome analyses. For each *Study*, the table lists the SG isolation *Method*, *Stress* stimulus, *Target* used for SG isolation, *Cell* types involved, and associated *Transcriptome* alterations (including transient changes), and GEO accession number (*GSEID*). G3BP refers to G3BP1 alone or both G3BP1 and G3BP2. For the PIC, PHOTON, and FANCI methods, *Target* refers to the antibody used to mark SGs via immunofluorescence, enabling photo-based or fluorescence-activated isolation. DC – Differential centrifugation; DC* – Differential centrifugation without immunoprecipitation; PL – Proximity labelling; PL** – Proximity labelling-derived technique; PIC – Photo-isolation chemistry; eCLIP-Seq – Enhanced cross-linking and immunoprecipitation sequencing; ARTR-Seq – Assay for reverse transcription-based RBP binding site sequencing; FANCI – Fluorescence-activated non-membrane condensate isolation; PHOTON – Photoselection of transcriptome over nanoscale.

| Study | Method | Stress | Target | Cell | Transcriptome | GSEID |
| --- | --- | --- | --- | --- | --- | --- |
| A. Khong 2017(31) | DC | Oxidative | G3BP | U2-OS | G3BP-GFP | GSE99304 |
| T. Matheny 2019(41) | DC* | Oxidative | None | U2-OS | Wild-Type | GSE138988 |
| A. Padrón 2019(42) | PL | Heat-shock and Hippuristanol | eIF4A | HEK293T | APEX2-eIF4A | GSE121575 |
| SP. Somasekharan 2020(29) | PL | Oxidative | G3BP | PC-3 | APEX2-G3BP | GSE138058 |
| T. Matheny 2021(43, 44) | DC | Oxidative and Hyperosmotic | PABPC1 and G3BP | U2-OS | Wild-Type, G3BP-GFP and G3BP KO | GSE119977 |
| **M. Honda**<br>**2021(45)** | PIC | Oxidative | G3BP | HeLa | Wild-Type | GSE143413 |
| **H. Shen**<br>**2022(46)** | PL | Oxidative | DDX3X<br>and<br>DDX3Y | HeLa | APEX2-DDX3X<br>and APEX2-DDX3Y | GSE171792 |
| **V. Iadevaia**<br>**2022(47)** | DC | Viral Infection | G3BP | U2-OS | G3BP-GFP | GSE171655 |
| **K. Sato**<br>**2023(40)** | DC and<br>eCLIP-Seq | Oxidative | G3BP | SH-SY-5Y | Wild-Type | GSE188397 |
| **N. Curdy**<br>**2023(6)** | DC | T-Cell<br>Activation | G3BP | Donor T-Cells | Wild-Type | GSE19700 |
| **Z. Ren**<br>**2023(48)** | PL** | Oxidative and<br>Hyperosmotic | G3BP | HEK293T<br>and U2-OS | G3BP-miniSOG | GSE223295 |
| **Y. Xiao**<br>**2024(49)** | ARTR-Seq | Oxidative | G3BP | HeLa | Wild-Type | GSE226161 |
| **Y. Zhou**<br>**2024(50)** | FANCI | Oxidative,<br>Hyperosmotic,<br>Heat-shock<br>and UV<br>Irradiation | G3BP | HeLa | DHX9-eGFP and<br>G3BP-mCherry | GSE218180 |
| **P. Kudrin**<br>**2024(51)** | DC | Oxidative | G3BP | HeLa | Wild-Type and<br>NAT10 KO | GSE212380 |
| **G. Di Timoteo**<br>**2024(52)** | DC | Oxidative | G3BP | SK-N-BE | Wild-Type,<br>METTL3 KD, and<br>FUS-P525L | GSE242766 |
| **S.**<br>**Rajachandran**<br>**2024(53)** | PHOTON | Oxidative | eIF3eta | HeLa | Wild-Type and<br>METTL3 KO | PRJNA1152621 |

For the final set of SG and WC transcriptomes corrected for centrifugation-associated RNA length biases (Figure 4A), studies were retained if they had appropriate technical controls (*i.e.*, not just SG samples, but also equivalent cellular fractions, empty vectors, in stressed and unstressed conditions) and at least three SG replicate samples (see Supplementary Methods). Even though most DC studies had no appropriate technical controls, we considered that our simulated SG samples could serve as such, and thus these studies were maintained.

Canonical transcript lengths were obtained from ENSEMBL (v. 108)(56).

ac4C RNA modification information was obtained from Arango et al. (2022)(57) Supplementary Table 2, “ac4C_gene_summary” sheet. m6A RNA modification information was obtained from Khong et al. (2022)(58) Supplementary Data 1.

### TCGA expression data

Expression data generated by the TCGA Research Network were downloaded from the GDC Data Portal (https://portal.gdc.cancer.gov/) (59, 60).

Read count data for each cancer type were pre-processed separately. We started by filtering out genes deemed uninformative due to their very low expression across samples: only genes with a normalised expression of at least 1 CPM in at least 40% of the samples were kept for analysis, using the same filtering criterion previously adopted in our analysis of other large-scale transcriptomic datasets(61).

### Dataset pre-processing

Downloaded raw reads (in FASTQ format) were trimmed using Trimmomatic (v0.39)(36) operating in either single-end (SE) or paired-end (PE) mode, depending on the original study design, with appropriate adapter sequences and parameters. Trimmed reads were then aligned to the hg38 reference genome using STAR(35) (v2.7.9a) with the following parameters:

--outFilterType BySJout \

--alignSJoverhangMin 8 \

--alignSJDBoverhangMin 1\

--alignIntronMin 20 \

--alignIntronMax 1000000 \

--alignMatesGapMax 1000000 \

--outFilterMultimapNmax 1 \

--outFilterMismatchNmax 999 \

--outFilterMismatchNoverLmax 0.04 \

Aligned reads (BAM files) were subsequently annotated using the GENCODE v42 primary assembly(62) and quantified with featureCounts(37) (v2.0.3).

### Read count data pre-processing

Raw read counts were first filtered to remove lowly expressed genes, independently for each dataset. For all datasets, filtering was applied across all samples. Filtered read counts, including those from TCGA, were then used to calculate library size scaling factors using the trimmed mean of M-values(63) (TMM; calcNormFactors function from edgeR (v3.36.0)(64)). Those read counts were subsequently normalised and log2-transformed using the voom(65) function from limma (v.3.50.3)(66).

### Differential expression analysis

To model expression changes between conditions (*e.g.*, SGs vs WCs, stressed vs unstressed cells, or stressed cells with vs without SGs), we fitted linear models to the gene expression data. The exact linear models used for each dataset are provided in the project’s GitHub repository, under each study’s corresponding GEOID. Standard errors for the models were then moderated using empirical Bayes shrinkage, and differentially expressed genes were defined separately for each dataset using empirical dataset-specific log2 fold-change (log2FC) values and B-statistics, rather than fixed thresholds. For splicing and RBP analysis, RBP-encoding genes with |log2FC| < 0.5 and adjusted p-value > 0.05 were considered non-differentially expressed.

### Correlations

Unless otherwise indicated, Spearman’s correlation coefficients (ρ) and the associated tests’ p-values were calculated for comparison between gene-level moderated t-statistics of differential expression. For correlations involving transcript length, the canonical (or the longest, in case of ambiguity) transcript obtained from ENSEMBL (v. 108)(56) was used.

### GSEA

For each gene expression contrast of interest, such as SG enrichment, GSEA(67) was performed using the fgsea (v1.20.0)(68) R package. Genes were ranked by t-statistics of differential expression, ordered from most positive to most negative, and analysed against the Hallmark gene sets(69). The normalised enrichment score (NES) was extracted to quantify the extent to which a given gene set is enriched at the extremes of the ranked gene list. Positive and negative NESs indicate gene sets enriched amongst over- and under-expressed genes, respectively. For transcript length enrichment analysis, genes were ordered from the longest to the shortest.

### Centrifugation simulation (Spin-Sim)

Transcript read counts were first normalised to Transcripts Per Million (TPM). Canonical nucleotide sequences for each transcript were retrieved from ENSEMBL(56) to determine nucleotide composition and calculate the molecular weight (MW) of each RNA using(70):

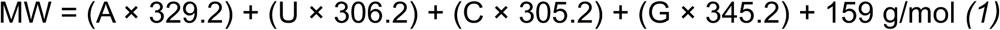

To simulate RNA sedimentation during centrifugation, we applied a simplified particle sedimentation model(71):

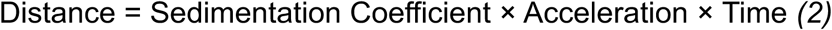

Since precise sedimentation coefficients depend on transcript-specific features (*e.g.*, shape, folding, and hydration), we used a heuristic approximation(72) assuming spherical RNA behaviour and sedimentation coefficients proportional to the cube root of MW:

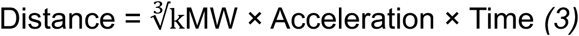

The centrifugation process was modelled after Khong et al. 2017(31), assuming initiation from the 1 mL mark in standard 2 mL microcentrifuge tubes (distance from bottom ∼2 cm). Each transcript’s sedimentation distance was scaled proportionally to estimate the fraction of each RNA that would sediment. These sedimented proportions were converted back to read counts using:

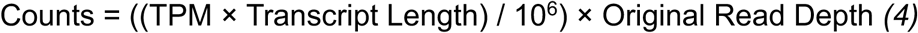

where *Original Read Depth* corresponds to the total number of reads in the associated SG sample, ensuring consistency between original and simulated datasets. This approach was used for the analyses presented in Figure 2.

**Figure 2.**
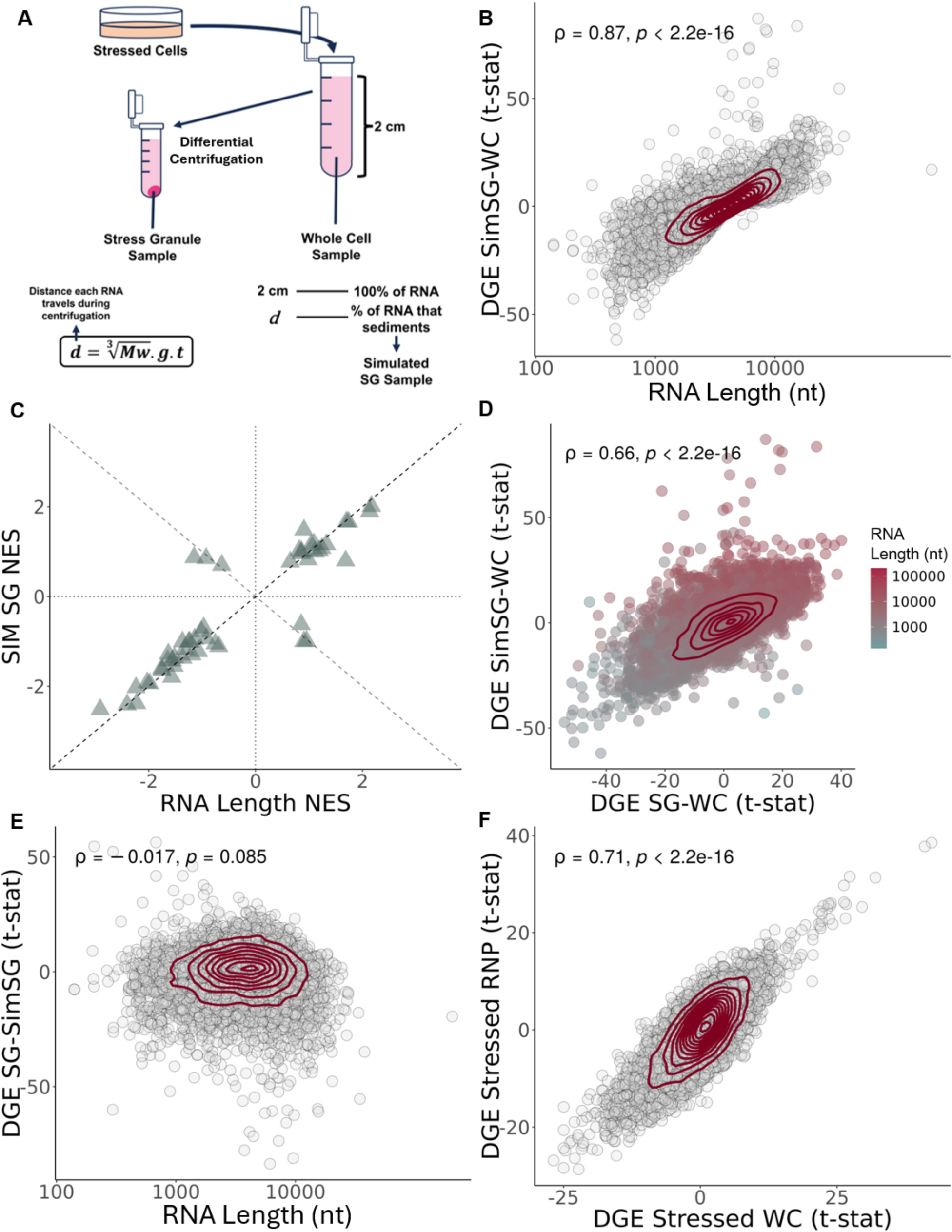
Length bias in DC-isolated SGs is compatible with inherent RNA condensation. (A) Schematic representation of standard DC protocol alongside our *in silico* pipeline for simulating the centrifugation process. *d* = distance travelled by each RNA during centrifugation; *Mw* = Molecular weight of each RNA transcript, in g/mol; *g* = centrifugation acceleration, in g; *t* = time of centrifugation, in seconds. (B) Scatter plot of t-statistics of DGE between simulated SGs, and WCs (Y-axis) *versus* length of gene’s representative RNA transcript (nt) (X-axis). (C) Scatter plot of GSEA NES of Hallmark gene sets in DGE between simulated SGs and WC samples, with genes ranked by t-statistics of differential expression (Y-axis) *versus* genes ranked by length (nt) (X-axis). Fisher’s test Odds Ratio (OR) and associated p-value on top right corner. Dashed lines depict identity (Y = X; Y = −X), and axis lines. (D) Scatter plots of t-statistics of DGE in simulated SGs (Y-axis) and DC-purified SGs (X-axis), relative to WC samples; genes as circles coloured by RNA length in nt; density contour lines in dark red, Spearman’s correlation coefficient ρ and associated test’s p-value on top left corner. (E) Scatter plots of t-statistics of DGE in DC-purified SGs *versus* simulated SGs (Y-axis) *versus* length of gene’s representative RNA transcript (nt) (X-axis, log scale); transcripts as circles; density contour lines in dark red, Spearman’s correlation coefficient ρ and associated test’s p-value on top left corner. (F) Scatter plots of t-statistics of DGE between stressed vs unstressed RNP samples (Y-axis) and between stressed vs unstressed WC samples (X-axis). Genes as circles; density contour lines in dark red, Spearman’s correlation coefficient ρ and associated test’s p-value on top left corner (B, D, E, F). The original WC expression from Khong 2017(31) was used to simulate SG samples.

For RNA modification experiments, transcript MW was adjusted by +14 g/mol for each m6A site(73) and +42 g/mol for each ac4C site(74).

Additional simulations used empirical data from Glauninger 2024(33), which measured RNA sedimentation in centrifuged yeast samples. From this dataset’s azide stressed samples, which we considered were comparable to the commonly used arsenite (Figure 1B) as both cause oxidative stress(75), we derived a linear model relating RNA length to sedimentation coefficient, which showed slightly stronger correlation than MW. After testing different models, we used the cube root of RNA length to fit the final model:

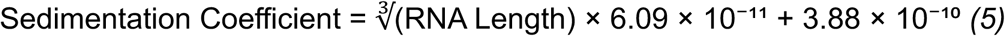

Mitochondrial RNAs exhibited disproportionately high sedimentation relative to size, so we derived a separate equation for them using a linear model relating RNA length to sedimentation coefficient, considering only mitochondrial encoded RNAs:

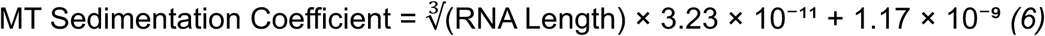

As before, sedimented RNA abundances were converted back to read counts using equation (*4*).

### Consensus SG Transcriptome

To derive a consensus SG transcriptomic signature, we computed the mean of per-gene t-statistics of differential expression between SGs and respective controls across datasets, including only genes present in all datasets, allowing the resulting consensus signal to remain directly interpretable in t-statistic units for downstream analyses.

### U-2 OS growth conditions

Human osteosarcoma U-2 OS cells were cultured in high-glucose DMEM supplemented with 10% fetal bovine serum and maintained at 37°C with 5% CO2. Cells were plated on glass coverslips in 24-well plates and used for all experimental procedures.

Reagents and material references are supplied in the Reagents and Resources Table.

### Stress conditions

To examine mRNA localisation under stress, when 80% confluency was reached, cells were exposed to 0.4 M sorbitol for 3 h. To inhibit mitochondrial dsRNA leakage into the cytosol, 50 μM 4, 4′-diisothiocyanatostilbene-2, 2′-disulfonate (DIDS) (as previously described(76)) was applied either in combination with sorbitol or alone (control) for the same duration.

Reagents and material references are supplied in the Reagents and Resources Table.

### Immunofluorescence

Following stress treatments, cells were washed three times with PBS+/+ pH 7.4. Fixation was performed with 4% paraformaldehyde in PBS+/+ for 20 min at room temperature, after which cells were washed three additional times with PBS+/+. Permeabilisation was carried out using 0.1% Triton X-100 in PBS+/+ for 10 min at room temperature, followed by three washes in unsupplemented PBS.

Glass coverslips were then removed from the 24-well plates and transferred to a humid chamber prepared with 20 µL droplets in blocking buffer with 5% goat serum. Coverslips were inverted onto the droplets (cell side facing down) and incubated for 60 min at room temperature. Primary antibodies against HSP60 (mitochondria), J2 (dsRNA), and G3BP1 or TIA1 (stress granules) were diluted 1:200 in blocking buffer with 5% goat serum and applied to coverslips using the drop technique (10 µL per coverslip).

Incubation proceeded for 2 h at room temperature in a humid chamber. Afterwards, coverslips were washed three times in unsupplemented PBS.

Secondary antibodies were diluted 1:500 in blocking buffer with 5% goat serum together with DAPI (1:1000) and applied to coverslips (10 µL per coverslip) for 1 h in the dark at room temperature. Coverslips were then washed three times in unsupplemented PBS with agitation (5 min each wash). For mounting, 2 µL of Vectashield was placed on each microscope slide, and two coverslips were carefully placed cell-side down onto the drops. Coverslips were sealed with nail polish before imaging.

PBS+/+ (supplemented with Ca2+ and Mg2+) was prepared as a 1 L solution by combining 600 mL of MilliQ H2O with 100 mL of 10x PBS-/- (unsupplemented). MgCl2·6H2O (33 mL of a 0.01 M stock solution; final concentration 0.33 mM) and CaCl2·2H2O (90 mL of a 0.01 M stock solution; final concentration 0.9 mM) were then added dropwise, with the solution brought to a final volume of 1 L with MilliQ H2O. pH was then adjusted to 7.4, followed by filtering.

The blocking buffer used throughout the immunofluorescence protocol was prepared by warming 1x PBS-/- (70 mL) and dissolving 0.2% tissue-culture-grade gelatin (0.2 g) with the aid of a magnetic stirrer. The solution was then cooled to room temperature before addition of 2% FBS (2 mL), 2% BSA (20 mL of a pre-made 10% BSA stock solution), and 0.3% Triton X-100 (1.5 mL of a 20% stock solution). The buffer was brought to its final volume with 1× PBS, followed by filtering.

Reagents and material references are supplied in the Key Resources Table.

### Microscopy and image analysis

For Supplementary Figures 11 and 14, after fixation, cells were imaged on a ZEISS LSM 880 confocal microscope, using a 63x/1.4 Plan-Apochromat DIC oil immersion objective. For each condition, a z-stack of a single field of view (1024 x 1024 pixels; pixel size 0.13 µm) was acquired, comprising 10–20 Z-planes, with a 0.48 µm step size, according to Nyquist, with a scan zoom of 1. G3BP1 and TIA1 were excited using a 633 nm laser, HSP60 using a 561 nm laser, dsRNA J2 using a 488 nm laser, and DAPI using a 405 nm laser. Images were observed in FIJI, with maximum intensity thresholds defined for each channel using sorbitol-stressed samples. These same parameters were then applied to all other conditions to ensure consistency across images.

For Figure 6, the same samples from Supplementary Figure 11 were re-acquired using a ZEISS LSM 980 with Airyscan 2 confocal microscope, with a 63x/1.4 Plan-Apochromat DIC oil immersion objective, using the Airyscan 2 in Super Resolution mode (Airyscan SR). Three fields of view were acquired per condition (2245×2245px, with a px size of 0.035 µm), with each z-stack comprising 35–65 Z-planes with a 0.130 µm step size, with a scan zoom of 1.7. G3BP1 was excited using a 639 nm laser, HSP60 using a 561 nm laser, dsRNA J2 using a 488 nm laser, and DAPI using a 405 nm laser. Raw Airyscan data were processed using Airyscan Joint Deconvolution to improve resolution (Sample structure “Dense” and 5 Maximum Iterations). Basal autofluorescence levels were determined from a control sample incubated with secondary antibodies only and used as a minimal threshold for normalising fluorescence signals in experimental images.

Colocalisation analysis was performed in Imaris 10.2.0 CF, using the Coloc module, quantifying the percentage of overlapping voxels across different channels. Initially, colocalisation between HSP60 (mitochondria) and J2 (dsRNA) was measured. Negative controls were used to define the minimum intensity threshold for colocalisation of each channel. A new channel was subsequently generated, containing only the J2 signal that did not overlap with HSP60. This non-mitochondrial J2 signal was then analysed for colocalisation with G3BP1 (stress granules) under both unstressed and sorbitol-stressed conditions. All image processing and quantifications were performed on an image analysis and processing workstation with support from the Bioimaging Platform of the Gulbenkian Institute for Molecular Medicine Foundation.

Image analysis software references are supplied in the Reagents and Resources Table.

Colocalisation proportions were compared between stressed and unstressed samples using Student’s *t*-tests. Given the limited sample size, beta distributions were also fitted to the three replicates per condition. From these fitted distributions, 1000 random samples were drawn and compared across several statistical metrics, including Cohen’s *d* and Area Under the Curve (AUC). Student’s test two sided p-value was calculated using the original values (n=3 per condition).

### Alternative hypothesis testing for SGs implication in cancer

To explore mechanisms beyond SG assembly that could account for the phenotypes observed in *G3BP1* and *PTGS2* knockout cells, specifically the effects on survival and drug sensitivity, we leveraged multiple computational resources. *PTGS2* perturbagens were identified using the Connectivity Map(77). Drug sensitivity profiles were obtained from the Genomics of Drug Sensitivity in Cancer (GDSC(78)) database.

### SG signatures

To define an expression signature indicative of SG presence in a given sample, we first created a “discovery” dataset by integrating GSE138988(41) and GSE119977(43, 44). This integration allowed comparison of stressed cells with and without SGs while controlling for confounding variables, including cell type, genotype, stressor, and study of origin. “Batch effects” arising from these variables were corrected before to analysis, following the methodology we had previously applied to the analysis of other large-scale transcriptomic datasets(61). We then employed an elastic net regression with 1000 bootstrap iterations with a mixing parameter (alpha) of 0.75, yielding a relatively stringent signature. Genes consistently selected in at least 900 iterations were retained. This signature included genes both up-(positive) and down-regulated (negative) in SG-positive cells.

SG score for individual samples was calculated using a non-parametric gene ranking method. Within each sample, all genes were ranked by expression. The SG score for a given sample was computed as the sum of the ranks of the genes in the signature, weighted by (*i.e.* multiplied by) +1 for positive or −1 for negative genes, and normalised by the number of genes in the set. This signature was validated on an independent dataset, GSE173953(7), with the same batch effect correction as mentioned above(61).

This rank-sum approach was also used to estimate mitochondrial/nuclear-encoded mitochondrial transcripts enrichment score.

For the tumour *v*ersus normal tissue comparison, resulting Welch’s t-test p-values were corrected for the number of tissues tested through the Benjamini-Hochberg (BH) procedure.

For survival analysis, GDC-TCGA samples(59, 60) were stratified into top and bottom quartiles based on SG scores (or expression of a specific gene, in Supplementary Figure 15). Kaplan-Meier survival curves were generated, and statistical significance was assessed using the log-rank test. To evaluate the significance of observed survival associations, 100 random signatures matching the original SG score signature in size and gene directionality (positive/negative weight) were generated by sampling gene names from the TCGA LAML dataset(59, 60). This dataset was chosen for its large number of expressed genes after filtering. Log-rank p-values were corrected for the number of tissues tested through the BH procedure. Survival analyses were repeated for each random signature, and false discovery rates were calculated as the proportion of random signatures with log-rank p-values lower than those obtained from the original SG signatures.

### Differential splicing analysis

We used GEO dataset GSE173953(7), which profiles U2-OS and A549 cells stressed with dsRNA, both with and without SG formation, because its sequencing depth was substantially higher than that of our alternative option (the “Discovery Dataset”(41, 43, 44)). High read depth is particularly critical for splicing analyses, as exon-exon junction reads and low-abundance isoforms are often underrepresented, making shallow sequencing insufficient to reliably capture alternative splicing events. Raw reads (FASTQ format) were first trimmed using Trimmomatic (v0.3.9)(36) in PE mode. Alternative splicing was quantified with VAST-TOOLS (v. 2.5.1), following the align (against human reference genome hg38) and combine steps(38).

Differential splicing analysis was then conducted using the betAS (v1.2.1)(79) R package, with modifications to the default filtering procedure. By default, the alternativeEvents function excludes events in which at least one sample does not meet the percent spliced-in (PSI) thresholds (1<=PSI<=99)(79). However, applying these thresholds resulted in the inclusion of a large number of events that appeared constitutive, showing minimal variation rather than true alternative splicing.

To improve specificity for alternative splicing, we incorporated prior information from the MAIN PSI TABLE from VASTDB (https://vastdb.crg.eu/wiki/Downloads)(38). For exon-skipping events, we computed the average PSI and variance across tissues and cell types. Events were classified as alternative if they satisfied at least one of the following empirical criteria: variance >= 750 or 12.5 <= PSI average <= 87.5. Only events meeting these criteria were retained for downstream filtering of the VAST-TOOLS output.

For shRNA experiments, data from the ENCODE project(80, 81) (HepG2 and K562 treated with scrambled shRNA controls) were processed jointly using our modified betAS splicing pipeline(79). The same process was applied to the samples from GSE173953(7), with U2-OS and A549 jointly processed. Although this approach yielded fewer differentially spliced events overall, the events identified are likely to represent more robust and reproducible changes.

#### RBP binding analyses pipeline (eCLIPSE)

eCLIPSE (eCLIP for Splicing Evaluation) is an as-yet unpublished computational pipeline developed to integrate eCLIP data from the ENCODE project with splicing annotations from VastDB(38), generating RNA splicing maps. By aligning eCLIP-derived RBP binding profiles to the genomic coordinates of annotated splicing events, eCLIPSE enables the systematic assessment of how individual RBPs regulate alternative splicing, either directly (via their own binding) or indirectly (through downstream effects on other RBPs). All components required to reproduce this pipeline are available on Zenodo (at https://doi.org/10.5281/zenodo.21724194) and at GitHub (at https://github.com/DiseaseTranscriptomicsLab/CHARM).

A brief description of each relevant step is provided in the sections below.

#### Splicing-genome creation

A reference “splicing-genome” was first created from VastDB annotations. Alternative exons shorter than 10 bp were excluded. For each exon-skipping event, specific regions of interest (as exemplified in Supplementary Figure 12) were defined as follows:

- Upstream exon end: −50 bp to the exon boundary
- Upstream intron start: intron boundary to +200 bp
- Upstream intron end: −200 bp to the intron boundary
- Alternative exon start: exon boundary to +50 bp
- Alternative exon end: −50 bp to the exon boundary
- Downstream intron start: intron boundary to +200 bp
- Downstream intron end: −200 bp to the intron boundary
- Downstream exon start: exon boundary to +50 bp

Additional rules for handling short exons or introns (<100 bp or <400 bp, respectively) were defined. When upstream or downstream introns were shorter than 400 bp, they were divided between the start and end regions, to ensure we maintained comparable regions of interest (*i.e.*, near the splice sites) for these events. For even-length introns, the split was equal; for odd-length introns, the extra base was assigned to the end. For alternative exons shorter than 100 bp, the same rule was applied. For alternative exons with alternative start sites, the start site that maximised exon length was chosen.

#### eCLIP samples

One eCLIP file per cell line for each available RBP from the ENCODE project(80, 81) (252 RBP files) was considered. Processed GRCh38 bed narrowPeak files from each RBP, labelled as “default”, were downloaded, and binding events with “Signal” greater than 3 were retained.

#### Binding Metagenome Assembly

RBP binding peaks were intersected with the predefined splicing regions to extract the binding signal in hg38 coordinates. For each RBP, each base in the original genome was coded as 0 (no binding) or 1 (binding present). To enable comparison across events of different lengths, each event was mapped to a standardised “splicing genome” coordinate system corresponding to the regions of interest (*e.g.*, upstream exon, upstream intron, alternative exon, downstream intron, downstream exon). Each splicing event was subsetted and rescaled to fixed splicing genome coordinates (*e.g.*, 1–50 corresponding to −50 bp to upstream exon end in hg38, 51–250 corresponding to upstream intron start to +200 bp, etc.), preserving the relative position of binding within the region. As stated above, because individual exons or introns can be shorter than the full reference ROI (*i.e.*, exons <100 bp or introns <400 bp), some positions in the splicing genome may not have corresponding hg38 coordinates with binding information. These positions are assigned NA to indicate missing binding data. Binding information in splicing genome coordinates was strand-specific. All conversions were implemented in Python3 (v3.10.11). Since Python indexing starts at 0, a +1 was added to Python-defined coordinates to ensure consistency with R-defined coordinates, whose indexing starts at 1.

#### RNA-binding maps

betAS-derived(79) differential splicing events were divided into categories as maintained, increased, or decreased, based on a chosen difference in PSI (dPSI) threshold (|10| in this analysis). For each of these categories, and per RBP, normalised binding profiles were computed across the splicing genome coordinates for the corresponding events. Specifically, for each nucleotide position in the splicing genome, the binding signal was calculated to ensure a per-position signal that accounts for missing values due to shorter regions in individual events, which would otherwise be overrepresented:

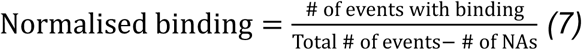

For assessment of differential binding, a chi-squared test was applied to each position using the following contingency matrix:

|  |  |
| --- | --- |
| Number of Increased/Decreased Events | Number of Maintained Events |
| Number of Increased/Decreased Events after Normalised Binding | Number of Maintained Events after Normalised Binding |

The resulting chi-squared statistic values from the test are then multiplied by −1 for positions with decreased binding.

Finally, position-wise normalised binding profiles were plotted along the splicing genome coordinates for each category, with the corresponding Chi-Squared statistic shown in the bottom half of the plot. An example of this analysis and respective plots is given as Supplementary Figure 12, using RBFOX2 as a case-study.

Taken together, these steps generate statistically assessed RBP binding maps for maintained, increased, and decreased exon inclusion events. This process was performed in R (v. 4.1.2).

## Results

### Different SG purification techniques yield different SG transcriptomes

We first compiled publicly available transcriptomic datasets of human SGs isolated through various methods (Table 1).

For each dataset, we compared gene (mRNA) expression between SGs and the corresponding whole-cell or cytoplasmic fraction (henceforth collectively referred to as WC). SGs isolated by DC consistently contained longer transcripts on average (Figure 1A, left; Khong 2017 study(31) shown as example), irrespective of stress stimuli and cell type (Figure 1B). Gene expression profiles of DC-purified SGs exhibit a consistently high correlation with each other (Figure 1B). In contrast, PL-based SG transcriptomes showed a weak enrichment for shorter transcripts (Figure 1A, right; Somasekharan 2020 study(29) given as example) and exhibited minimal correlation between SG gene expression profiles, even under the same stress stimulus (Figure 1B).

We performed Gene Set Enrichment Analysis (GSEA)(67) for the Hallmarks collection(82) (designed to minimise redundancy between gene sets) on differentially expressed genes (DEGs) in SGs induced by oxidative stress in the DC and PL example datasets, compared with their WC fraction (Figure 1A, B). In DC-purified transcriptomes, Hallmark enrichment in SGs was strongly confounded with transcript length (Fisher’s p < 1.5e-5, Odds Ratio = 23.77), with several enriched and depleted Hallmarks associated with long (*e.g.*, mitotic spindle) and short (*e.g.*, oxidative phosphorylation) transcripts, respectively (Figure 1C). In PL-purified transcriptomes, the observed enrichment in short transcripts in SGs (Figure 1A, right) was also reflected on the Hallmark enrichment analysis (Figure 1C; p < 0.0001, Odds Ratio = 0.06).

Notably, these datasets differ in cell lines and arsenite concentrations, factors previously suggested to influence SG transcript composition(29, 83). To evaluate whether these biological variables could explain the observed differences, we examined the transcriptomic oxidative stress response across the corresponding cell lines (Supplementary Figure 1A). Arsenite-treated U2-OS and PC-3 cells share highly upregulated genes, including *MT1X* and *MT2A* (metallothioneins, a protein class protective against oxidative stress(84)), *DUSP1* (upregulated during oxidative stress(85)), and *DNAJB1*/*HSP40* (which mitigates oxidative stress–induced cytotoxicity(86)). Furthermore, GSEA on the DEGs between stressed and unstressed states in both cell lines revealed similar Hallmark enrichment (Supplementary Figure 1B; Fisher’s p < 3.5e-4, Odds Ratio = 21.66). Consistent with this, WC transcriptomes from these cell lines under oxidative stress also showed a degree of correlation in the stress response (Supplementary Figure 1A), together indicating that cell line alone is unlikely to account for the differences seen in SG transcriptomes. Most likely, the variation in SG transcriptomes reflects the different isolation methods used in these example datasets.

### Simulation of the centrifugation process yields RNA length bias compatible with that observed in DC-isolated SGs

SG assembly is driven by molecular aggregation, suggesting that longer RNAs, with greater potential for multivalent interactions, may be preferentially enriched. Nevertheless, the strong correlation observed between SG and WC transcriptomes (Supplementary Figure 1B) is compatible with the null hypothesis that the SG transcriptome is just a sample of the cell’s cytosolic transcriptome. This raises the possibility of the RNA length bias in DC-purified SGs not being biological but rather a technical artefact.

We first considered whether this bias could originate from sampling artefacts inherent to RNA-seq library preparation; however, *in silico* simulations ruled out this possibility (Supplementary Data, Supplementary Figures 2 and 3).

RNAs tend to naturally form condensates in the cell, independently of their localisation to SGs(32, 33). Thus, DC purification, which isolates condensate-rich fractions prior to their immunoprecipitation, may also capture RNA condensates other than SGs. To test this, we simulated centrifugation *in silico* using WC expression data(31), modelling RNA precipitation as dependent primarily on transcript molecular weight under the assumption of uniform condensate shape (Figure 2A). Although RNAs sediment as part of condensates, we modelled each transcript as sedimenting independently, using its molecular weight as a proxy for the condensate (see Methods). This approach yields a gene expression profile attributable solely to the technical effect of centrifugation, which we interpret as putative SGs. The simulated SGs displayed a positive RNA length bias relative to WC samples, matching that seen in the original DC data(31) (Figure 2B). GSEA for Hallmarks on DEGs between simulated SGs and WC samples was also confounded by transcript size (Fisher’s p < 1e-9, Odds Ratio = 44.18) (Figure 2C), consistent with previous results (Figure 1C).

We found a strong correlation between simulated and DC-purified SGs in DEGs relative to WC samples (Figure 2D). Since the simulated SGs are derived only from WC expression and lack SG-specific biological signals, we interpret them as representing the centrifugation component of the DC method. Comparing transcriptomes between the simulated SGs and the DC-purified ones allows us to identify transcripts sequestered beyond what would be expected from their sedimentation alone. These “*corrected*” SG transcriptomes show no length bias (Figure 2E), suggesting that the reported length bias in the original studies likely stems from centrifugation-related artefacts in the purification process. Although this simulation relies on coarse approximations, most notably the assumption that each RNA transcript behaves as an independent sphere, the fact that even this simplified model closely reproduces the DC length bias strengthens the interpretation that centrifugation is a dominant contributor to this effect.

If the SG signal is confounded by natural RNA condensation, a more appropriate control for DC-purified SGs requires identically processed samples lacking SGs. In the Matheny 2019 dataset(41), WC and ribonucleoprotein (RNP) granule fractions were collected under stressed and unstressed conditions, using DC but without immunopurification(41). Stressed RNPs fractions contain SGs together with other granules, such as processing bodies (PBs), a type of cytoplasmic granules also present in unstressed cells and linked with RNA degradation, whereas unstressed RNPs lack SGs(41). If SG-specific RNA sequestration occurs beyond effects of stress and centrifugation, stressed versus unstressed RNP transcriptome differences would noticeably deviate from those observed in homologous WC comparisons. The two comparisons are strongly correlated (ρ = 0.71; Figure 2F), indicating that the SG transcriptome predominantly mirrors the cellular transcriptome, be it stressed or unstressed, with minimal transcript-specific sequestration. Variance analysis further shows that SG-specific effects are smaller than those attributable to stress and centrifugation (Supplementary Figure 4). Notably, this study did not perform immunopurification for a core SG protein(41), thus potentially diluting the SG-specific signal among other RNA condensates and explaining their lack of clear detection. Also, one fewer centrifugation step was performed compared to other DC protocols, which may explain the weaker RNA length bias observed in this dataset, even relative to other DC studies using the same cell types and stress conditions (Figure 1B).

Beyond RNA length and translation efficiency(32), RNA modifications such as m^6^A and ac^4^C have been associated with SG enrichment obtained through DC(51, 52, 87, 88). To evaluate whether this association is genuine, we compared differential SG versus WC expression in modification-knockout cells relative to wild-type counterparts, using interaction model residuals to isolate modification-dependent effects on SG sequestration (Supplementary Figure 5A). While m^6^A knockout produced no discernible shift in SG enrichment, ac^4^C knockout was associated with a modest but detectable effect (Supplementary Figure 5B). However, the magnitude of this shift correlated only weakly with the per-transcript number of ac⁴C modifications (Supplementary Figure 5C), arguing against a direct causal role for the modification itself. *In silico* simulations taking into account the additional weight from these modifications showed only limited correlation with enrichment. Instead, RNA length substantially accounted for the observed differences (Supplementary Figure 5D), suggesting that the ac⁴C-associated signal reflects a length-dependent bias rather than a direct effect of the modification on SG transcript sequestration.

### Differential centrifugation yields similar transcriptomes for SGs and other RNP granules

Based on the observations above, we hypothesise that the SG transcriptome isolated by DC is substantially confounded by the centrifugation process itself rather than representing exclusive biological sequestration. Consequently, other samples isolated by similar DC protocols, regardless of their identity, should yield comparable profiles, with enrichment for longer RNAs.

In the Matheny 2019 dataset(41), stressed PBs and bulk RNP granules were purified by DC, with PBs distinguished by an additional immunoprecipitation step using EDC3(41). The original Khong 2017 dataset(31) used a similar three-round DC protocol to isolate SGs (using G3BP for immunoprecipitation).

Despite representing distinct cytoplasmic granules, DC-purified SGs and PBs displayed similar transcriptomes (Supplementary Figure 6A), as noted by the authors(41). Moreover, both SGs and PBs were more similar to each other than to bulk RNP granules purified by two rounds of centrifugation (Supplementary Figure 6B-C). It is unclear if this pattern reflects the contribution of multiple other condensate types in the bulk RNP granule preparations or, more simply, the additional centrifugation step applied to both SGs and PBs.

Additionally, in Matheny 2019(41), these DC-isolated stressed PBs showed lower similarity to unstressed PBs isolated by fluorescence-activated particle sorting (FAPS) from Hubstenberger 2017(89) than to SGs. We confirmed this observation using another independent FAPS dataset, Kodali 2024(54), where, albeit correlated, stressed DC-isolated PBs still exhibited higher similarity to either DC-isolated SGs or RNPs (Supplementary Figure 6A-D).

In Zhou 2024(50), stress granules were isolated using fluorescence-activated non-membrane condensates isolation (FANCI), a flow cytometry–based approach similar to FAPS. Notably, even after batch correction for cell type and stress (see Methods), FANCI/FAPS-isolated SGs remained less similar to DC-isolated SGs than the latter are to PB-isolated SGs, despite SGs and PBs being distinct structures (Supplementary Figure 6E).

While Matheny 2019(41) posits that stress causes a shift in the PB transcriptome by increasing the pool of available nontranslating mRNAs in the cell(41), hence increasing their similarities to SGs, our reanalysis does not support this explanation.

We integrated the Zhou 2024(50) and Kodali 2024(54) datasets, which used these flow cytometry-based methods in HeLa cells. Both studies included unstressed WC controls, allowing batch correction for stress (see Methods). This adjustment confirmed that similarities between SGs and PBs isolated by FAPS/FANCI persist even after controlling for stress, but they remain far weaker than those observed with DC isolation (Supplementary Figure 6F).

It is important to note that this batch correction approach relies on the assumption of equivalent stress effects in SGs and PBs, which may not fully capture more complex biological interactions. Nevertheless, in the absence of stressed PB data purified by FAPS, it remains uncertain whether the divergence between DC and FAPS datasets reflects genuine stress-induced remodelling of PBs or, instead, an artefact introduced by centrifugation, such as length-dependent RNA bias.

### Method-specific corrections for distinct confounders reveal SG-specific RNA sequestration

To assess the generalisability of our centrifugation hypothesis, we analysed publicly available Sed-seq data(33), which measured RNA sedimentation by sequencing pelleted and unpelleted fractions from stressed and unstressed yeast cells. This experimental dataset enabled us to derive transcript-specific sedimentation coefficients (*S*) (*i.e.*, how much a particle sediments, see Methods), offering a more precise estimate of RNA sedimentation than our first weight-based-only approximations.

Applying these coefficients to simulate SG transcript-sequestration against WC in an independent yeast dataset(31) yielded a strong correlation with observed SG profiles (Supplementary Figure 7A). These results support our hypothesis in an experimental system, albeit in a different organism.

We next extended this framework to human cells by deriving formulas linking *S* to RNA length (see Methods). These enabled us to estimate theoretical *S* values for human RNAs and simulate their expected sedimentation in human samples, providing an *in silico* model of putative SGs. Simulations based on *S* values more closely matched empirical data than our earlier weight-based approach (Figure 3B *versus* Figure 2D). Comparing observed SG sequestration(31) with these simulations allowed us to identify transcripts enriched or depleted beyond their predicted *S* values, thereby isolating a corrected SG signal that reflects sequestration independent of centrifugation effects. As a proof of concept, this approach recovered known SG markers, such as *NORAD* as enriched and *GAPDH* as depleted, consistent with previous RNA FISH validation(31) (Supplementary Figure 7B). The corrected SG signal exhibited no meaningful correlation with RNA length (Figure 3C), differing from the original study(31). GSEA reveals that corrected SG transcriptomes were enriched for mitotic spindle-related transcripts (Figure 3D), consistent with the possibility that SGs selectively sequester cell-cycle-associated RNAs to reduce cell division during stress. However, this interpretation is challenged by concurrent signatures of TNF-α signalling via NF-κB pathway downregulation (a core stress response pathway) and upregulation of metabolic pathways, leaving the functional implications of SG sequestration in cellular regulation unresolved.

**Figure 3.**
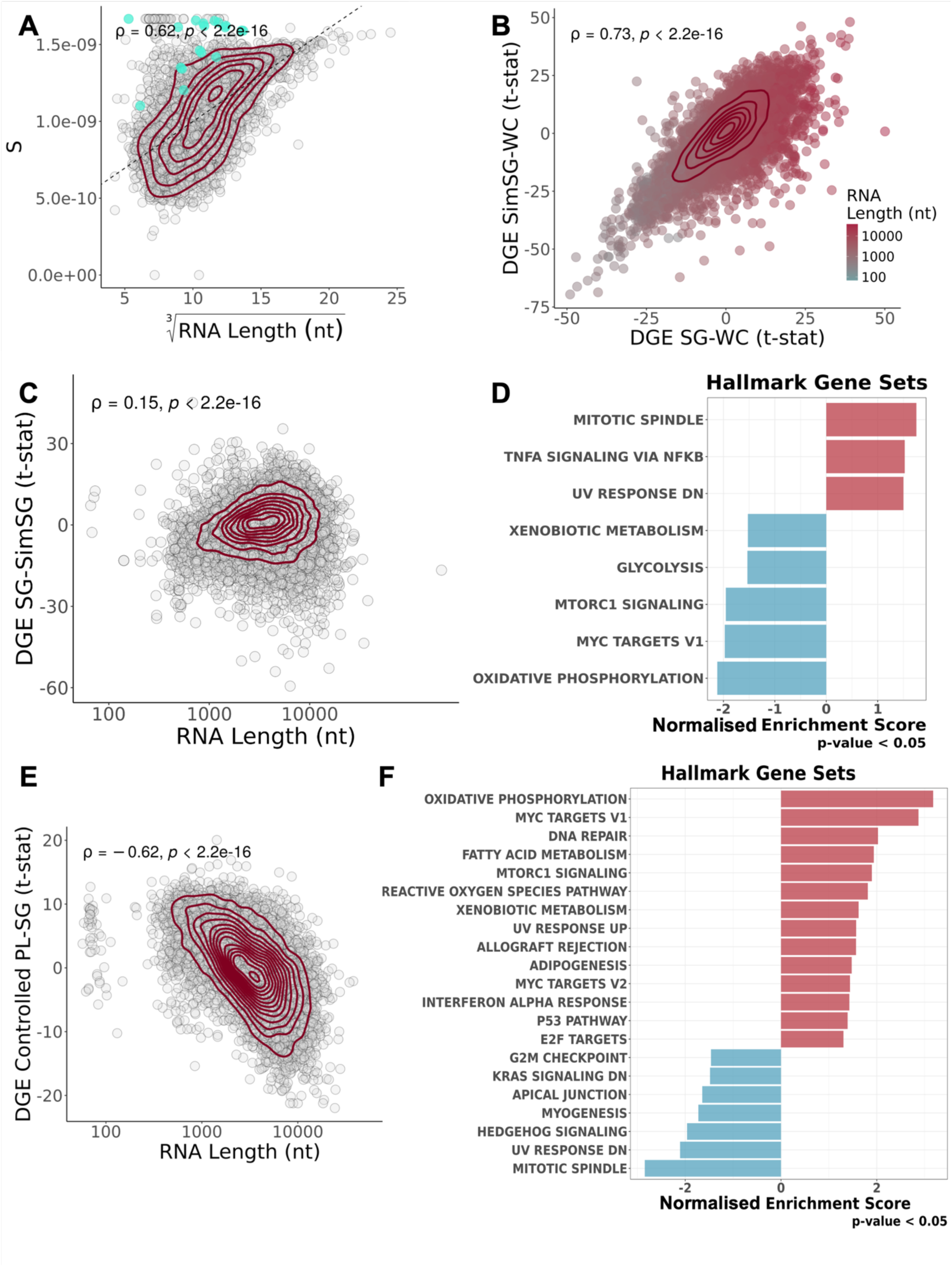
Method-specific corrections for SGs transcriptomes unveil their specificities. (A) Scatter plot of yeast *S* values obtained from Sed-seq data(33) (Y-Axis) and the cubic root of yeast RNA length (nt) (X-Axis); Each circle denotes a transcript, with mitochondrially encoded transcripts coloured in cyan; density contour lines in dark red, Spearman’s correlation coefficient ρ and associated test’s p-value on top left corner. The dashed line corresponds to the least-squares linear regression fit between the two variables. (B) Scatter plot of t-statistics of DGE of simulated SGs *versus* WCs, with S estimations based on Sed-seq data(33) (Y-Axis) and DGE of DC-purified SGs *versus* WCs(31) (X-Axis); transcripts as circles coloured by RNA length; density contour lines in dark red, Spearman’s correlation coefficient ρ and associated test’s p-value on top left corner. (C) Scatter plot of RNA length (X-Axis, log scale) and t-statistics of DGE of DC-Purified SGs *versus* Simulated SGs(31) (thus what we consider to be centrifugation-corrected SGs) (Y-Axis); transcripts as circles; density contour lines in dark red, Spearman’s correlation coefficient ρ and associated test’s p-value on top left corner. (D) GSEA NES of Hallmark gene sets, with genes ranked by t-statistics of differential expression, in Simulated SGs *versus* DC-Purified SGs(31). (E) Scatter plot of t-statistics of DGE in linear modelled PL-isolated SGs(29) (Y-Axis) and RNA length (X-Axis, log scale); transcripts as circles; density contour lines in dark red, Spearman’s correlation coefficient ρ and associated test’s p-value on top left corner. (F) GSEA NES of Hallmark gene sets, with genes ranked by t-statistics of differential expression, in linear modelled PL-isolated SGs(29).

To further refine our understanding of SG composition in human cells, we applied a correction strategy to account for both centrifugation and stress effects using the Matheny 2019 dataset(41). This dataset includes both unstressed and stressed RNP and WC samples, which can control for centrifugation effects from the DC method. However, directly comparing RNPs from stressed and unstressed cells (which, as abovementioned, contain SGs in the former) is likely to primarily reflect stress-induced changes rather than SG-specific composition. To disentangle SG-specific signal from general stress responses, we applied linear models that accounted for both centrifugation and stress effects (see Methods and Resource Availability). This yielded a refined SG profile showing enrichment for *NORAD* and depletion of *GAPDH* and *ACTB* (Supplementary Figure 4C), as originally validated with RNA FISH(31). This controlled RNP/SG-specific signal exhibited a positive, but weaker than previously reported, correlation with RNA length (Supplementary Figure 7D). Mirroring the DC-corrected data (Figure 3D), GSEA shows downregulation of pathways related to metabolism (Supplementary Figure 7E). However, it also suggests downregulation of apoptosis pathways (Supplementary Figure 7E), challenging the hypothesis that SGs inhibit apoptosis by sequestering pro-apoptotic transcripts(12, 13).

PL methods, although not relying on centrifugation, are also subjected to confounding factors. Notably, the labelling protein G3BP1 does not exclusively localise to SGs(29), and thus PL may capture RNAs beyond bona fide SGs. Although we cannot directly control for this, we used linear models with interaction effects, similar to our approach with the Matheny 2019(41) dataset, and controlled for other confounding variables (see Methods) to isolate the SG-specific gene expression signal. In PL-purified SGs, this showed a negative correlation with RNA length (Figure 3E), and GSEA revealed upregulation of metabolism and stress response pathways, consistent with selective transcript sequestration (Figure 3F).

While the previous corrections provided a putative SG-specific gene expression signal, WC transcriptomes can offer complementary insights into the effects of SGs across the cell. Importantly, as the majority of publicly available datasets do not purify SGs, inferring their presence and effects in the cell from WC transcriptomes would prove invaluable in studying their effect/function in other biological contexts. However, a major challenge in analysing the WC transcriptome is separating the general stress response from SG-specific contributions. G3BP1/G3BP2 KO cells, which do not form visibly detectable SGs under oxidative stress, allowed us to distinguish these effects(41, 43, 44). By comparing KO and WT cells under stressed and unstressed conditions, we isolated the contributions of SGs, stress, and the KO itself to the WC transcriptome (see Methods)(41, 43, 44). Contrary to their theorised function in increasing the efficacy of the stress response, SGs appear to hamper it, as evidenced by the downregulation of inflammation-related pathways (Supplementary Figure 7F).

### Meta-analysis of SG transcriptomes suggests SG-related dampening of inflammatory signalling

Given the differing results across SG purification methods, we applied our correction strategies to the remaining datasets we deemed suitable (see Methods), and compared them (Figure 4A). When DC-purified PBs were available, they were compared with DC-purified SGs, to control for the technique. We used this approach for the Matheny 2019 dataset, as we consider that it better recaptures the SG transcriptome than simply comparing RNP granules alone.

**Figure 4.**
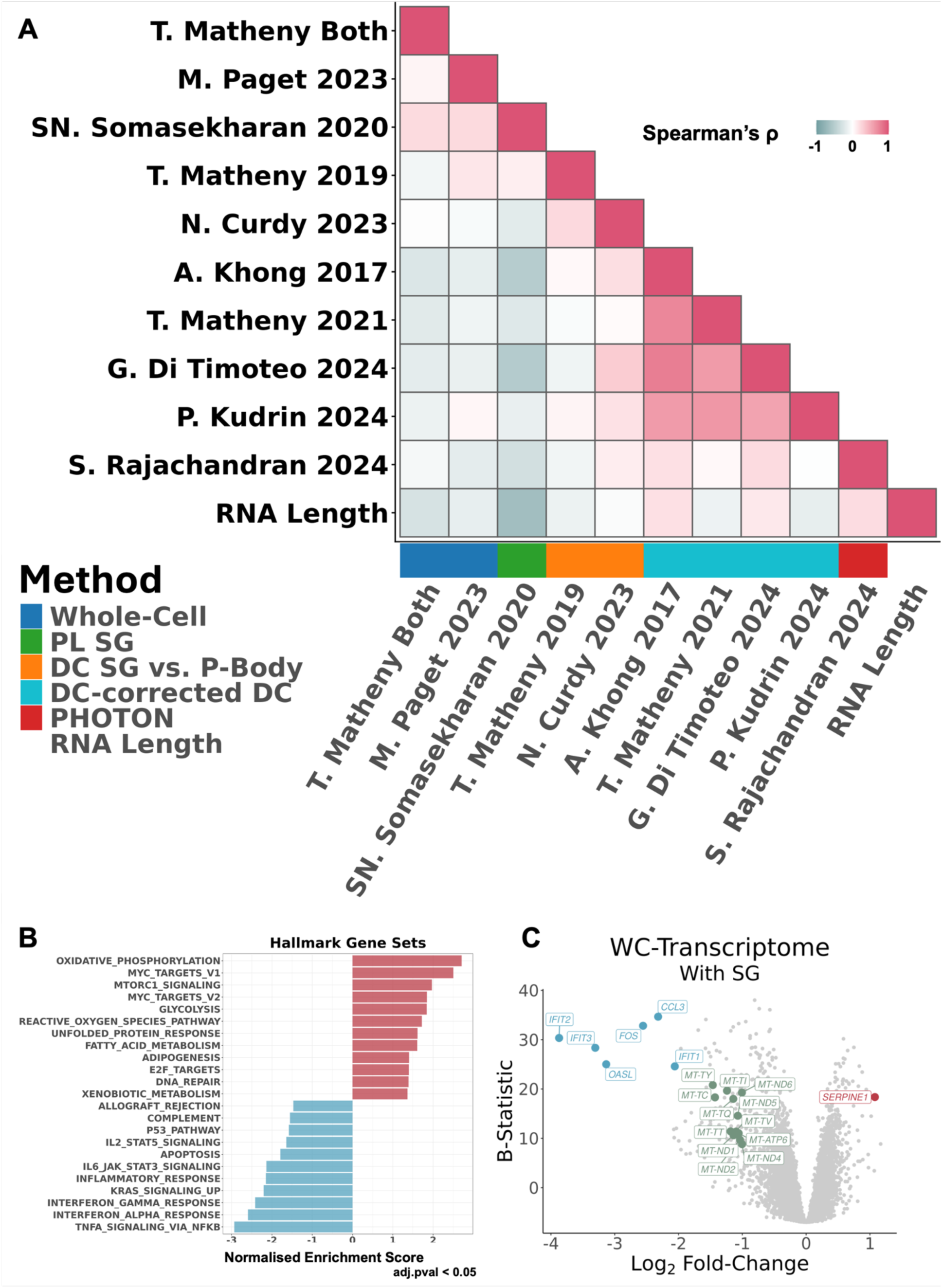
Meta-analysis of SG transcriptomes reveals SG assembly-specific gene expression alterations in whole cells. (A) Heatmap of Spearman’s correlations between SG and WC transcriptomes, corrected for centrifugation-associated RNA length biases, from different studies and RNA length. The SG purification method of each study is given by the colour below the heatmap. (B) GSEA NES of Hallmark gene sets, with genes ranked by t-statistics of differential expression, of the SG-specific signal (*i.e.*, what changes in the cells’ gene expression upon SG assembly)(7, 41, 43). (C) Volcano plot of DGE (X-Axis – Log2 Fold-Change; Y-Axis – B-Statistic) of the SG-specific signal; transcripts as circles; most differentially expressed transcripts are coloured red (if upregulated, log2FC>1 with B-Statistic>15), blue (if downregulated, log2FC< −2 with B-Statistic>15), or green (if mitochondrially encoded, log2FC< −1).

Despite our corrections, substantial discrepancies remain across datasets (Figure 4A), independent of purification method. The exception is the transcriptomes of DC-purified SGs corrected for centrifugation, consistent with one another. SGs may largely reflect a random sample of cellular transcripts, with a difficult to detect, small minority of specifically sequestred RNAs, and each purification method may introduce biases that shape observed profiles. WC transcriptomes generated with standard RNA-seq are also relatively similar. Notably, the PL dataset from Somasekharan 2020(29) more closely resembles WC transcriptomes than other SG datasets (Figure 4A), perhaps reflecting that only ∼20% of G3BP1 localises to SGs.

Our corrections largely remove length bias in the transcriptomes of DC-purified SGs (Figure 4A). By contrast, the PL study Somasekharan 2020(29) finds enrichment for short RNAs, while PHOTON (Rajachandran 2024(53)) preferentially captures longer transcripts (Figure 4A). These inconsistencies underscore the persistent challenge of reliably purifying and characterising SG transcriptomes. GSEA also revealed no pathways consistently enriched or depleted in SGs across all methods (Supplementary Figure 8A-Left).

Because WC-transcriptomes are less affected by technical biases from purification methods, we first focused on these datasets, integrating and analysing them together (Matheny 2019(41), 2021(43, 44) and Paget 2023(7)). GSEA identified consistent downregulation of stress- and inflammation-related pathways in SG-containing WC transcriptomes, including interferon responses, TNF signalling via NF-κB, and IL6–JAK–STAT signalling (Figure 4B). Apoptosis pathways were also downregulated, supporting the proposed role of SGs as inhibitors of apoptosis. Functionally, SGs may act as “shock absorbers”, preventing excessive immune responses, a phenomenon previously observed with dsRNA(7). Notably, similar patterns were observed regardless of the stressor, as the T. Matheny(41, 43, 44) datasets used oxidative and hyperosmotic stresses.

We observed downregulation of *FOS*, an immediate-early stress-responsive gene(90), and upregulation of *SERPINE1*, a senescence-inducing gene and component of the senescence-associated secretory phenotype (SASP)(15) (Figure 4C).

Several datasets showed an enrichment of mitochondrially encoded genes in the putative SG. To rule out a technical explanation for this enrichment, we revisited our yeast sedimentation coefficients. We observed that mitochondrial transcripts consistently exhibited higher S values than other RNAs of comparable length, perhaps reflecting their polycistronic nature(76) (Figure 3A, teal circles). As such, we recalculated sedimentation coefficients using separate equations for nuclear and mitochondrial transcripts. After this correction, the enrichment persisted in a subset of datasets (Supplementary Figure 9, DC-studies without mitochondrial correction in Supplementary Figure 10). To investigate this further, we performed GSEA across all datasets using a gene set containing the thirty-six mitochondrially encoded transcripts obtained from Ensembl(91) (Supplementary Figure 9C). As a control, we analysed nuclear-encoded transcripts localised to mitochondria, selecting top-scoring genes from the RNALocate database(92) (Supplementary Figure 9C). Mitochondrially encoded transcripts were significantly depleted in both WC studies and in two SG transcriptome studies, (Matheny 2019(41), Di Timoteo 2024(52)) and significantly enriched in three (Somasekharan 2020(29), Kudrin 2024(51), Rajachandran 2024(53)), with the remainder showing no significant enrichment or depletion (Supplementary Figure 9). We found no evidence that these results can be generally attributable to expression-level or length-based biases in the underlying data (Supplementary Figure 9B).

Intriguingly, those results are not easily reconciled with known transcript properties. Mitochondrial transcripts are generally short, yet the Rajachandran et al. 2024 dataset(53) showed a bias toward longer RNAs in SGs but also displayed enrichment of those short mitochondrial transcripts (NES = 2.26, adj.p = 0.0035). Moreover, the Kudrin et al. 2024 study(51), even after mitochondrial sedimentation correction, also showed mitochondrial gene enrichment in SGs (NES = 2.91, adj.p < 0.001), independently corroborated by elevated mitochondrial signature scores in empirical versus simulated SGs (Cohen’s f = 1.76, adj.p = 1.20e-05).

To assess the specificity of these signals, we repeated the analysis using 100 random gene sets matched in size to each mitochondrial group, thereby deriving an empirical False Positive Rate (FPR). This yielded FPR < 0.05 in eight mito-encoded datasets, four enriched and four depleted (including both WC studies), as well as in three nuclear-encoded datasets, one depleted, and two enriched (one WC study). The opposing directions of enrichment and depletion across datasets may reflect genuine variation in experimental conditions, such as cell line, stress type, and duration, rather than noise, as the specificity of the signal relative to size-matched random gene sets argues against a chance finding. Notably, the relative absence (when compared to mitochondrial) of a signal for nuclear-encoded mitochondrial genes suggests that this effect is specific to mitochondrially encoded transcripts rather than a general property of mitochondria-associated biology. Collectively, these results suggest that mitochondrially encoded transcripts may be dynamically regulated, albeit with only a modest effect, in the context of SG formation in a condition-dependent manner, warranting further investigation. It is important to note that the direction of this modulation, whether they are enriched or depleted in the SG, is unclear from our analyses. Given the specificity, but limited magnitude of this signal, we next sought to test experimentally whether SGs sequester mitochondrial dsRNA (results shown below).

To derive a consensus SG transcriptomic signal, we averaged t-statistics of DGE across datasets (see Methods). Although mito-encoded RNAs are typically enriched in SGs, they are not among the most strongly enriched transcripts overall (Supplementary Figure 8B). Still, their enrichment is strongly significant (Supplementary Figure 8C), suggesting their high sensitivity in marking biological effects and/or technical biases. Among the most consistently enriched transcripts, of particular interest is *PPP1R15A*, encoding for a protein that promotes apoptosis and inflammation through the NF-κB pathway(93), potentially contributing to the anti-inflammation observed upon SG assembly(7).

To further explore the disagreement between studies for individual transcripts, we examined *NORAD*, the RNA with the highest presence-in-SG score (Supplementary Figure 8B) and a commonly used SG marker(31). *NORAD* displayed very strong enrichment in SGs in some datasets but also substantial disagreement between the underlying studies (Supplementary Figure 8B). We found that the strong enrichment was driven almost entirely by DC studies (Supplementary Figure 8D). In contrast, other purification methods showed comparatively little or no enrichment of *NORAD*. This discrepancy was not mitigated by our correction procedures, suggesting that it at least partially reflects a genuine methodological difference in SG capture.

### Mechanistic model of SG-mediated inflammation dampening via mitochondrial RNA sequestration

To explore mechanisms underlying the dampening of stress and inflammatory signalling suggested by our meta-analysis, we compared the transcriptomic signal of WCs bearing SGs with that of SGs relative to their host WCs and observed no meaningful correlation (Figure 5A).

**Figure 5.**
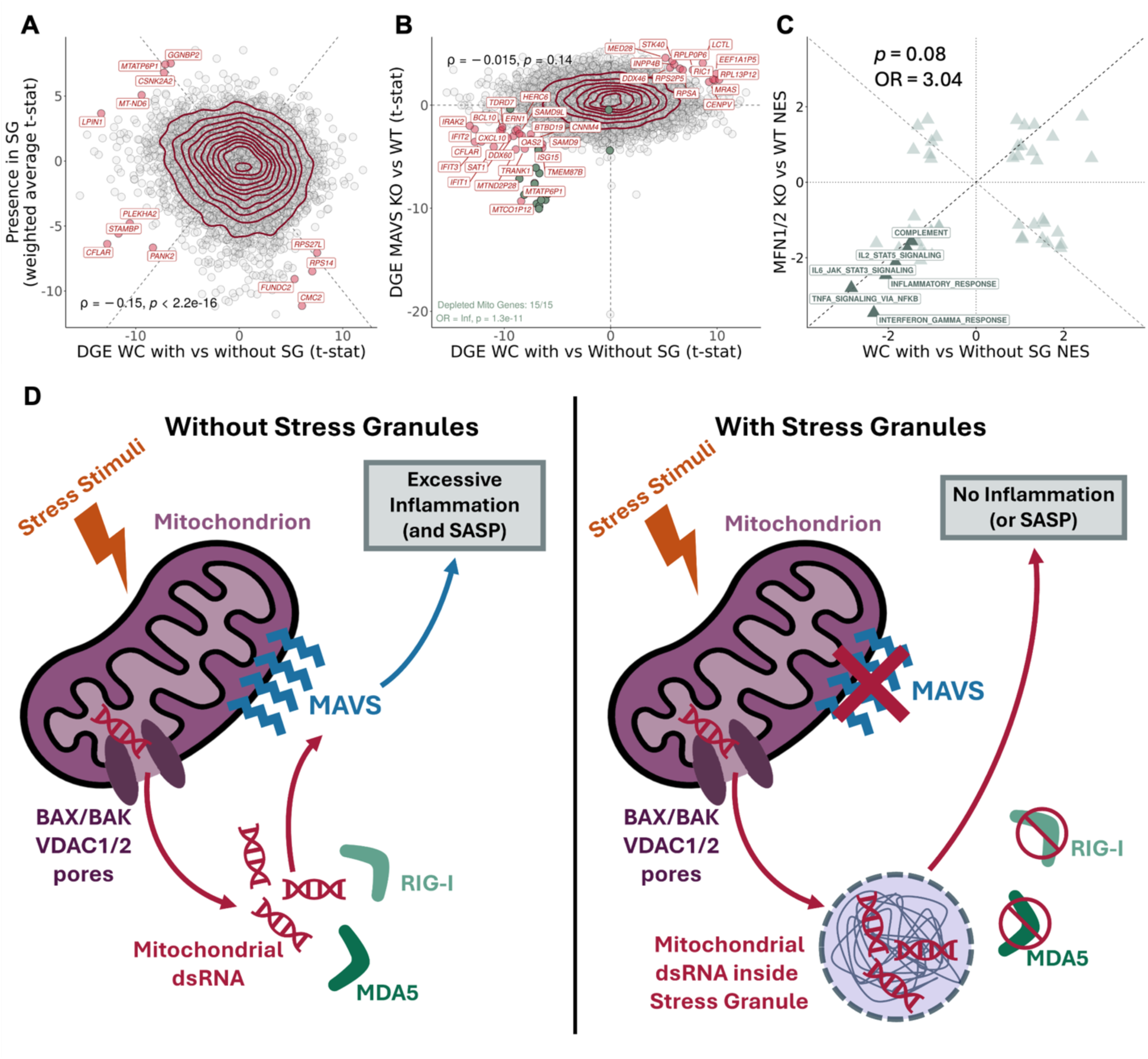
Mechanistic model of mitochondrial dsRNA–SG axis in inflammation control. (A) Scatter plot of t-statistics of DGE between SG-bearing WC samples and their non-bearing counterparts(7, 41, 43) (X-Axis), and weighted average of t-statistics (see Methods) of DGE between SGs and WCs across studies (Y-Axis); transcripts as circles; transcripts with most differential/concurrent enrichment between axes labelled and coloured dark red (|WC Enrichment * SG Enrichment| > 40 t-statistic); density contour lines in dark red, Spearman’s correlation coefficient ρ and associated test’s p-value on bottom left corner. (B) Scatter plot of t-statistics of DGE between SASP-inhibited WC samples(55) through siRNA treatment against MFN1/2(55) and their non-inhibited counterparts (Y-Axis – SASP Inhibition), and t-statistics of DGE between SG-bearing WC and their non-bearing counterparts(7, 41, 43); transcripts as circles; trancripts with most conserved enrichment between axes labelled and coloured dark red (SG Enrichment * SASP Inhibition > 20), and green if mitochondrially-encoded; density contour lines in dark red, Spearman’s correlation coefficient ρ and associated test’s p-value on top left corner; Fisher’s test for mitochondrially-encoded transcripts on bottom left corner. (C) Scatter plot of GSEA NES of Hallmark gene sets, with genes ranked by t-statistics of DGE, in SG-bearing cells compared to their non-bearing counterparts(29, 50, 51, 53) (X-Axis) and SASP-inhibited samples against their non-inhibited counterparts(55) (Y-Axis). Fisher’s test OR and associated p-value are shown in the plot. Hallmarks linked to inflammation are highlighted and labelled. (D) Schematic of the proposed mitochondrial dsRNA–SG axis in inflammation control. In the absence of SGs (left), cellular stress promotes mitochondrial nucleic acid leakage, likely via BAX/BAK and VDAC1/2 pores(55). Leaked dsRNA is sensed by RIG-I and MDA5, which signal through MAVS to trigger an inflammatory cascade(55). When SGs are present (right), leaked dsRNAs are sequestered within them, shielding these RNAs from RIG-I and MDA5 recognition and thereby preventing or attenuating inflammatory signalling. In A and C, dashed lines depict identity (Y = X; Y = -X). In C, dotted lines depict X = 0 and Y = 0.

Consistent with SGs’ reported role as senescence inhibitors(15), we found that a recent study targeting the SASP showed downregulation of pathways similar to those we observed in SG assembly(55). However, a direct comparison reveals notable differences in gene expression between SASP inhibition and SG formation (Figure 5B). Nevertheless, both conditions shared downregulation of mitochondrially encoded RNAs and, although overall correlations were weak, GSEA revealed broad suppression of inflammation-related pathways (Figure 5C).

Since senescent cells exhibit cytoplasmic accumulation of mitochondrial dsRNA that drives inflammation(55), and SGs have been reported to counteract senescence(15), we propose a working model (Figure 5D). In the absence of SGs, cellular stress promotes cytoplasmic leakage of mitochondrial dsRNA, which activates the RIG-I/MDA5/MAVS innate immune pathways(55). These innate immune pathways normally detect viral nucleic acids (like dsRNA) and trigger interferon and inflammatory responses, ultimately promoting SASP(94). Despite its cellular intrinsic origin, it has been previously shown that mitochondrial dsRNA can activate these pathways(94). When SGs are present, they may sequester these leaked nucleic acids, possibly explaining their frequent proximity to mitochondria(95), thereby dampening inflammatory signalling and perhaps even delaying SASP onset.

Indeed, several recent studies have shown that mitochondrial dsRNA can localise to SGs, either through increased mitochondrial permeability(76) or stress-induced release through fibrosis(96) or ageing(97). However, this raises an important issue: both fibrosis and ageing are strongly linked to oxidative stress(98, 99), which in turn promotes mitochondrial dysfunction and nucleic acid leakage(100). It therefore remains unclear whether the proposed mitochondria–SG–inflammation axis represents a general stress response, or whether it is specific to stresses that directly disrupt mitochondrial integrity.

### Immunofluorescence suggests selective sequestration of mitochondrial dsRNA into SGs during hyperosmotic stress

To avoid confounding effects of oxidative stress, which is tightly linked to mitochondrial dysfunction and nucleic acid leakage(100), we instead used hyperosmotic stress (through sorbitol exposure) in U2-OS cells, a widely employed model in SG research (reviewed in Figure 1B). Although hyperosmotic stress can impact mitochondrial function(101), we reasoned that it would perturb mitochondria less directly than oxidative stress, allowing us to better assess stress granule-associated RNA dynamics.

Previous work has shown that inhibition of either BAX/BAK or the mitochondrial porins VDAC1/2 prevents leakage of mitochondrial dsRNA into the cytoplasm(76). We hypothesised that this mechanism would also apply during hyperosmotic stress and therefore included the VDAC1/2 inhibitor 4, 4’-diisothiocyanatostilbene-2, 2’-disulfonate (DIDS)(76) as an additional control. Detecting dsRNA within SGs under these conditions would strengthen the case that its origin is mitochondrial rather than from alternative sources.

To test our model and directly visualise the putative SG–mitochondria–inflammation axis, we performed immunofluorescence using antibodies against G3BP1 (SG marker, reviewed in Figure 1B) and HSP60 (mitochondrial marker(102)), J2 (dsRNA marker(76)), and DAPI (nuclei).

Using the conditions described in Matheny 2021(43, 44), we confirmed by using imaging that U2-OS cells exposed to 0.4 M sorbitol for 3 h robustly formed SGs, visualised as G3BP1-circular foci, which were absent in unstressed controls (Supplementary Figure 11). Strikingly, co-treatment with 50 μM DIDS(76), a VDAC inhibitor, completely abrogated SG assembly (Supplementary Figure 11). This finding suggests that mitochondrial permeability via the VDAC channel activity, is required for SG formation under hyperosmotic stress (or that it impedes G3BP1 localisation into SGs).

Notably, SGs have previously been reported to modulate mitochondrial permeability by regulating VDAC activity(95). The observation that VDAC inhibition blocks SG assembly could point toward a reciprocal relationship, in which SGs both influence and depend on mitochondrial signalling.

At the same time, closer inspection showed that in both stressed and unstressed cells treated with DIDS, G3BP1 displayed a broadly similar diffuse distribution (Supplementary Figure 11). This suggests that DIDS may also exert off-target or global cellular effects, making it difficult to attribute the phenotype exclusively to VDAC inhibition.

Nevertheless, the requirement for VDAC activity is consistent with a model in which mitochondrial-derived signals contribute to SG assembly. While leakage of mitochondrial nucleic acids remains a plausible mechanism, especially given VDAC’s reported role in their transport(76), other damage-associated products (*e.g*., cytochrome, cardiolipin, metabolites) could also serve as triggers for inflammatory signalling and subsequent SG assembly(103). Alternatively, impaired import of mitochondrial factors might explain the effect.

Because SGs preferentially formed near mitochondria(95), this imaging setup did not provide sufficient spatial resolution to directly quantify dsRNA colocalisation within SGs.

To overcome the resolution limitations of conventional confocal microscopy, we re-imaged the same samples using Airyscan with joint deconvolution, which markedly improved spatial resolution. Because DIDS treatment altered the baseline distribution of G3BP1, we restricted our comparisons to stressed versus unstressed conditions only. At this resolution, G3BP1 signal appeared as small, spherical foci in both stressed and unstressed cells, rather than the diffuse distribution typically observed in unstressed cells with conventional confocal imaging (Figure 6A). This made it more challenging to distinguish bona fide SGs from background G3BP1 signal. This observation highlights a broader issue in the field: image-processing choices for SG detection are rarely reported in detail, yet they can substantially influence what is interpreted as an SG.

**Figure 6.**
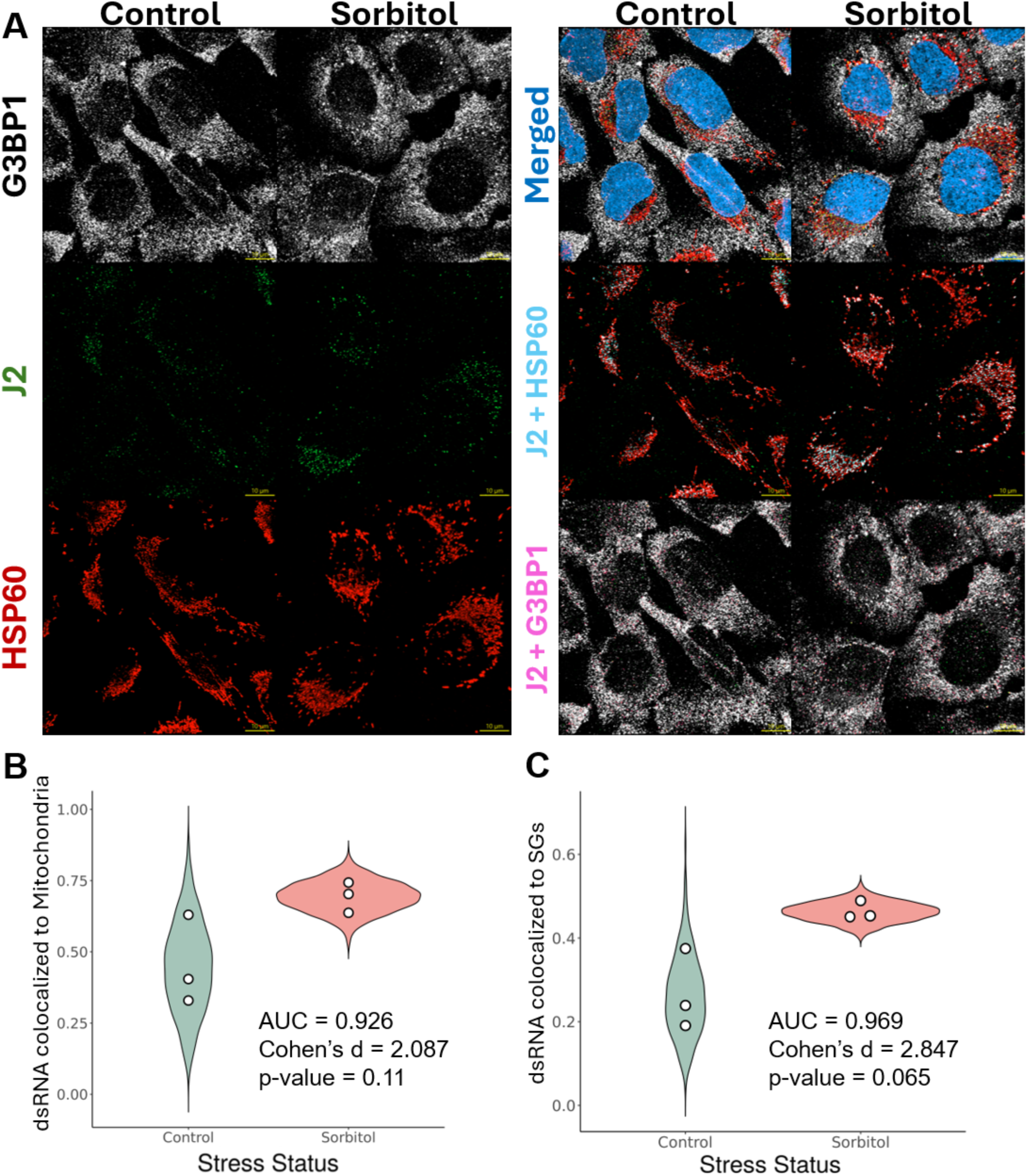
SGs potentially uptake leaked mitochondrial dsRNA. (A) Representative immunofluorescence images of U2-OS cells untreated or treated with sorbitol (0.4M). G3BP1 (SG marker, see Figure 1B) is shown in white, J2 in green (dsRNA marker(76)), HSP60 in red (mitochondrial marker(102)), and nuclei (DAPI) in blue. Merged images of these previous channels are also provided. In the “Cyan” row, a merge of J2 is shown with HSP60. Colocalisation of these volumes (in voxels) is coloured in cyan. In the “Magenta” row, a merge of J2, that did not colocalise with HSP60, is shown with G3BP1. Colocalisation of these volumes (in voxels) is coloured in magenta. The scale bar is shown on bottom right corner of each image, in yellow (10 μm). (B-C) Colocalisation proportions between J2 and HSP60 (B) or G3BP1 (C) were first quantified across all images acquired in the experiment (n = 3). For each condition, beta distributions were then fitted to both the empirical measurements and the corresponding null distributions generated from randomised colocalisation controls. Shown are violin plots representing 1000 values sampled from these fitted beta distributions for each condition. White dots denote the observed empirical colocalisation values. Statistical metrics (AUC, Cohen’s d, p-value) are displayed directly on the plot. AUC: probability that a randomly selected *Sorbitol* sample has a higher value than a randomly chosen *Control* sample; p-value obtained from a Student’s t-test. In B and C, AUC and Cohen’s d values were obtained by comparing the two sets of 1000 values sampled from the fitted beta distributions, whereas the p-value was obtained by comparing the two sets of empirical triplicates.

First, we calculated the fraction of mitochondrial volume that colocalised with dsRNA volume (in voxels) (Figure 6A, in Cyan). Unexpectedly, mitochondrial–dsRNA overlap was greater in stressed than in unstressed cells. This may reflect stress-induced remodelling of mitochondrial morphology that increases measured colocalisation, or enhanced dsRNA production under acute stress. Alternatively, even under nominally unstressed culture conditions, cells may experience a baseline level of stress insufficient to trigger SG assembly yet sufficient to promote mitochondrial dsRNA leakage, consistent with our observation that TIA1 displayed a cytoplasmic rather than nuclear distribution even in unstressed cells. Because SGs are thought to have protective functions(7), their assembly during acute stress might reduce such leakage, independently of any direct uptake. Moreover, evidence from *C. elegans* suggests that inhibition of protein synthesis enhances mitochondrial function(104). Since protein synthesis is typically suppressed under strong stress(11), this could paradoxically increase mitochondrial activity and dsRNA production.

To accurately assess SG-associated uptake, we excluded dsRNAs overlapping both mitochondria and SGs before quantifying their colocalisation with G3BP1. Using this filtered signal, we then measured the proportion of dsRNA volume colocalising with G3BP1 signal (Figure 6A, in Magenta). Colocalisation was higher under stress, both within mitochondria (Figure 6B) and in SGs, the latter being consistent with dsRNA sequestration into SGs (Figure 6C). To account for the limited sample size, data were modelled using beta distributions (see Methods). The small sample size (n=3, per condition) makes statistical significance meaningless, but the large observed effect size (AUC = 0.969; Cohen’s d = 2.847) (Figure 6C) suggests the hypothesis that SGs sequester cytoplasmic dsRNA under hyperosmotic stress is worthy of further exploration. Given the technical limitations and modest sample size, these data should be regarded as supportive rather than definitive, and further experimental validation will be required to consolidate this model.

Taken together, these observations suggest the hypothesis that SGs sequester cytoplasmic dsRNA under hyperosmotic stress. However, given the technical limitations and modest sample size, these data should be interpreted as supportive rather than definitive, and additional experiments will be required to fully validate this model.

### Sequestration of TIA1 in SGs recapitulates splicing alterations from TIA1 depletion

In addition to transcriptomes, we investigated SG-associated proteins, with a focus on RBPs, which account for roughly 50% of the SG proteome(8). Because only a fraction of the total amount of these proteins in the cell is sequestered in SGs (*e.g.*, only ∼20% of G3BP1(29)), the functional impact of their partial sequestration remains unclear. RBPs are multifunctional, with roles spanning splicing regulation, translational control, and RNA stability, among others. When sequestered inside SGs, they are displaced from the nucleus, potentially impairing their functions therein, namely splicing regulation, and leading to downstream biological effects.

To test this, we compiled a set of RBPs known to localise to SGs(8, 30) and mapped their RNA targets using eCLIP data (see Methods). We then performed a differential alternative splicing analysis (DSA) between dsRNA-stressed cells and unstressed cells(7), focusing on exon skipping events classified as increased, unchanged, or decreased, assessing binding enrichment for each of these categories (see Methods, Supplementary Figure 12 as example). This analysis identified six RBPs (TIA1, QKI, DDX3X, RBM15, PCBP1, and YBX3) with significant binding enrichment or depletion in these splicing events. To minimise confounding between expression-driven splicing alterations, we excluded RBPs whose expression differed between stressed and unstressed cells, in both SG-assembly competent and incompetent cells(7) (Supplementary Figure 13A), resulting in a final list of 4 potential targets, QKI, TIA1, PCBP1, and YBX3.

To assess whether these splicing changes were consistent with RBP sequestration, we compared splicing patterns in cells with shRNA-mediated knockdown of each candidate RBP(80, 81) (Supplementary Figure 13B-F – Left). RBPs whose RNA binding profiles resembled the splicing changes observed upon knockdown were considered likely to affect cellular function through SG sequestration (Supplementary Figure 13B-F – Middle). Of the four RBPs, only TIA1 and YBX3 showed visually similar patterns, with Pearson correlations of 0.44 and 0.77, respectively, between their binding profiles and KD-induced splicing changes (using the Chi-Squared statistic). However, these effects may stem from general stress-induced changes in RBP localisation or function, rather than SG sequestration specifically.

To determine whether these effects were SG-specific, we repeated the splicing analysis in G3BP1/G3BP2 knockout cells exposed to dsRNA, a condition under which SGs do not form(7) (Supplementary Figure 13B-F – Right). Without SG assembly, the correlation for TIA1 dropped markedly (r = 0.07), suggesting that its impact on splicing is specifically contingent on SG formation and thus consistent with functional impairment through SG sequestration. In contrast, the correlation for YBX3 remained substantial (r = 0.64), indicating that the pattern of its impact on splicing reflects a general stress response independent of SG assembly.

We next performed immunofluorescence for TIA1 under control and hyperosmotic stress conditions. Although we initially considered including DIDS to block SG formation under stress, providing a closer parallel to our *in silico* comparisons, the compound’s broad off-target and cell-wide effects (Supplementary Figure 14A) led us to focus instead on a direct comparison between stressed and unstressed cells. Under homeostatic conditions, TIA1 is typically nuclear(105, 106), but in our experiments the signal was predominantly cytoplasmic, likely reflecting the baseline stress inherent to cell culture. Despite this, we did not detect obvious differences in TIA1 localisation between stressed and unstressed samples. Nevertheless, TIA1 staining reliably marked SGs, consistent with its canonical role (Supplementary Figure 14A).

Beyond its roles in SG assembly(107), TIA1 is a known splicing regulator(108), implicated in various neurological disorders(109). Alongside TIAR, it participates in viral infection and replication(110), with viruses such as West Nile and dengue shown to interact with both proteins, inhibiting SG formation(110). Whether this reflects a viral strategy to disrupt SG function or an indirect consequence of altering TIA1 activity remains unresolved. Importantly, the gene expression changes (Supplementary Figure 14B) and the splicing alterations (Supplementary Figure 14C) following TIA1 knockdown(80, 81) did not reproduce the differences observed upon SG-assembly in U2-OS and A549 treated with dsRNA(7). TIA1 sequestration is therefore unlikely to fully explain the anti-inflammatory phenotype associated with SG assembly.

### Proposed links between SGs and cancer may reflect indirect mechanisms

Given the established link between inflammation and cancer(111), SGs’ involvement in inflammatory regulation may help explain this association. Namely, SGs have been reported to enhance the metastatic potential of sarcoma cells(27), based on comparisons between SG-competent and SG-deficient cells. However, SG ablation in these studies was achieved via G3BP1 knockout, a protein with functions beyond SG assembly, which may have influenced the phenotype. Recent studies describing SGs as tumour-promoting in pancreatic cancer specifically disrupted G3BP1’s domain responsible for SG assembly(112), but functions of this domain beyond those could similarly affect the phenotype. Moreover, G3BP1 knockout cells can still form SGs under certain stressors, such as hyperosmotic stress(43), which may occur in the tumour microenvironment. Thus, it remains unclear whether SGs were truly ablated, complicating interpretation of their direct role in metastasis.

Multiple genes, including *G3BP1*(113), *AGO2*(114), *BRF1*(115), *CAPRIN1*(116), *CPEB1*(117), *DDX6*(117), *DISC1*(118), *EIF2A*(119), *FASTK*(120), *FMR1*(121), *FXR1*(121), *G3BP2*(113), *L1TD1*(122), *SAMD4A*(123), *SMN1*(124), *TIA1*(125), *TIAL1*(125) and *ZFP36*(115), are known to induce spontaneous SG assembly when overexpressed (compiled elsewhere(126)). We hypothesised that, if SGs directly drive tumour aggressiveness, high expression of these SG-nucleating genes would correlate with poorer patient survival. Analysis of Genomic Data Commons’s The Cancer Genome Atlas (GDC-TCGA)(59, 60) sarcoma dataset revealed that only *AGO2* expression was significantly associated with patient survival (Supplementary Figure 15; HR = 2.43, Benjamini-Hochberg-adjusted p-value = 0.0178). Notably, *G3BP1*, the primary SG nucleator implicated in the sarcoma metastasis study, showed no significant association (Supplementary Figure 15). While the *AGO2* association is nominally consistent with a pro-tumourigenic role of SG assembly, it is more parsimoniously explained by its well-established SG-independent function as a core component of the RNA-induced silencing complex (RISC) whose dysregulation has been independently linked to cancer progression(127).

SGs have also been proposed to promote chemotherapy resistance(28). KRAS mutant cells reportedly enhance SG assembly paracrinally via PTGS2 upregulation(28). However, PTGS2 has SG-independent roles, and observed resistance may not be solely due to SG assembly. KRAS mutant cells exhibit constitutive activation of the EGFR pathway(128). As KRAS functions downstream of EGFR, its mutation predicts poor response to EGFR inhibitors but improved response to MEK inhibitors, which act further downstream(129) (Supplementary Figure 16A, B). PTGS2 KD or inhibition leads to expression changes that correlate with EGFR KD (and ERBB3 KD, another protein from the same family), suggesting that PTGS2 may indeed be part of the same pathway (Supplementary Figure 16C). Hypothesising that PTGS2 acts downstream of KRAS, its inhibition may mimic MEK inhibition, targeting a key oncogenic driver and explaining increased treatment sensitivity. While PTGS2 is known to be regulated by the EGFR pathway (reviewed in(130)), this proposed mechanism remains to be experimentally validated.

### A SG gene signature and its clinical value in cancer

The results above highlight the limitations of single-gene analyses in isolating SG-specific effects due to overlapping roles in other stress-related pathways. To address this, we developed a custom SG transcriptomic signature. First, we integrated datasets from Matheny 2019 and 2021(41, 43), as this combination had stressed samples with and without SGs, applying batch effect correction for cell genotype, stressor type, and study of origin. We then used an elastic net regression model with bootstrapping to derive a SG signature (see Methods). Genes retained in nearly all (>90%) bootstrap iterations were used to construct the final signature, comprising 41 genes, which we dubbed the SG Score. The SG Score was validated on an independent dataset, Paget 2024(7), which had also undergone batch correction for the abovementioned variables, and successfully distinguished SG-positive from SG-negative samples (Figure 7A, B). Nevertheless, it is important to note that the SG Score provides a relative, not binary, measure of SG presence. This means that higher scores indicate a greater likelihood but do not confirm SGs are exclusive to samples with elevated values, and that basal SG scores in distinct cell types and states may be different.

**Figure 7.**
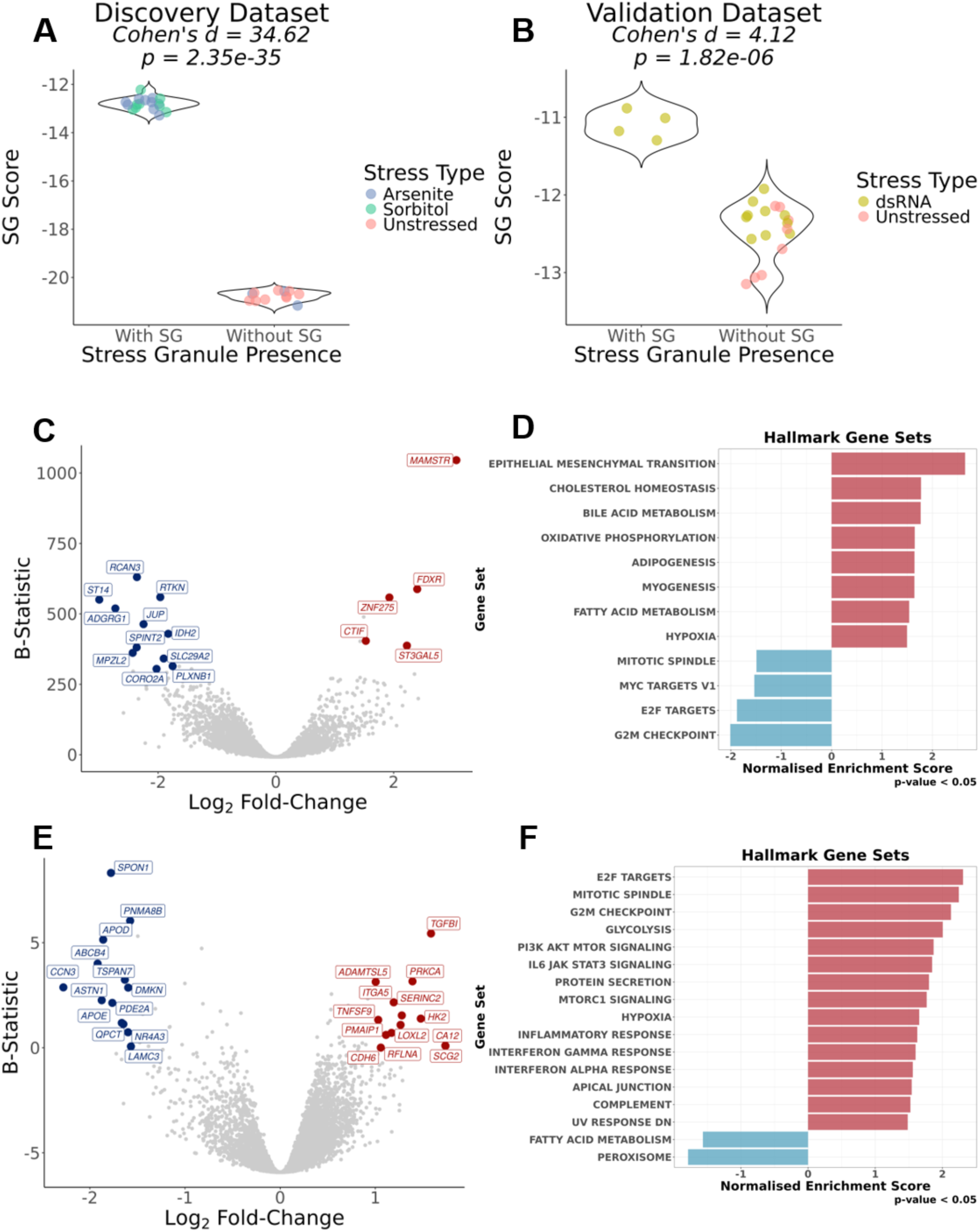
Development and validation of SG Score and its prognostic potential in GDC-TCGA datasets. (A/B) Violin plots of SG signature scores, calculated through a ranking metric (see Methods), in cells without and with SGs. WC samples are coloured in accordance with stress-type stimuli. The discovery dataset (A) refers to Matheny 2019 and Matheny 2021(41, 43), whereas the validation dataset (B) refers to Paget 2024(7). Effect Size (Cohen’s d) and corresponding statistical significance, given by Welch’s t-test (p-value) are shown above the plots. (C) Volcano plot of DGE (X-Axis – Log_2_ Fold-Change; Y-Axis – B-Statistic) between ACC and MESO tumour samples, and remaining GDC-TCGA tumour cohorts(59, 60), corrected for dataset of origin; genes as circles; most differentially expressed genes are coloured red (if upregulated, log2FC>1.5 with B-Statistic>300) or blue (if downregulated, log2FC< −1.75 with B-Statistic>300); (D) GSEA NES of Hallmark gene sets, with genes ranked by t-statistics of DGE, between ACC and MESO tumour samples, and remaining GDC-TCGA cohorts(59, 60), corrected for dataset of origin; (E) Volcano plot of DGE (X-Axis – Log_2_ Fold-Change; Y-Axis – B-Statistic) between top 25% and bottom 25% of SG-Score tumour samples in ACC and MESO, corrected for dataset of origin; genes as circles; most differentially expressed genes are coloured red (if upregulated, log2FC>1 with B-Statistic>0) or blue (if downregulated, log2FC< −1.5 with B-Statistic>0); (F) GSEA NES of Hallmark gene sets, with genes ranked by t-statistics of differential expression, between top 25% and bottom 25% of SG-Score tumour samples in ACC, and MESO, corrected for dataset of origin.

We next evaluated the performance of our SG signature in GDC-TCGA datasets(59, 60). These include: Acute Myeloid Leukemia (LAML); Adrenocortical Cancer (ACC); Bile Duct Cancer (CHOL); Bladder Cancer (BLCA); Breast Cancer (BRCA); Cervical Cancer (CESC); Colon Cancer (COAD); Endometrioid Cancer (UCEC); Esophageal Cancer (ESCA); Glioblastoma (GBM); Head and Neck Cancer (HNSC); Kidney Chromophobe (KICH); Kidney Clear Cell Carcinoma (KIRC); Kidney Papillary Cell Carcinoma (KIRP); Large B-cell Lymphoma (DLBC); Liver Cancer (LIHC); Lower Grade Glioma (LGG); Lung Adenocarcinoma (LUAD); Lung Squamous Cell Carcinoma (LUSC); Melanoma (SKCM); Mesothelioma (MESO); Ocular melanomas (UVM); Ovarian Cancer (OV); Pancreatic Cancer (PAAD); Pheochromocytoma & Paraganglioma (PCPG); Prostate Cancer (PRAD); Rectal Cancer (READ); Sarcoma (SARC); Stomach Cancer (STAD); Testicular Cancer (TGCT); Thymoma (THYM); Thyroid Cancer (THCA); Uterine Carcinosarcoma (UCS).

Given that cancer cells often encounter conditions known to promote SG formation(2, 25, 26), we hypothesised that our SG signature would yield higher scores in tumour samples compared to their normal counterparts. Among the 17 TCGA cancer types with at least 5 normal and tumour samples, only three: BLCA, LIHC, and PRAD, showed an increased SG Score in tumour samples (t-test, p < 0.05). In contrast, seven cancer types: ESCA, HNSC, KIRC, KIRP, LUAD, LUSC, and STAD, displayed lower SG Scores in tumours compared to normal tissue (t-test, p < 0.05), in particular ESCA and STAD (Cohen’s d < − 1) (Supplementary Figure 17).

To assess prognostic relevance, we performed a survival analysis comparing patients with tumour samples in the top and bottom quartiles of SG scores, based on the rationale that more aggressive, therapy-resistant cancers would show higher scores, and thus lower overall survival(27, 28) (Supplementary Figure 18).

Since random signatures tend to exhibit prognostic value in cancer(131, 132), we further tested the SG Score against randomly generated signatures to assess their specificity (see Methods; Supplementary Figure 18). Compared to 100 random signatures, our signature showed a false positive rate (FPR) < 0.05 in two cancer types (ACC and MESO). In both, higher SG Scores were consistently associated with improved patient survival, contrary to the hypothesis that SG assembly promotes tumour aggressiveness. One possible interpretation is that SG formation in these tumour types reflects an intact stress response capacity, which may correlate with a less aggressive phenotype or greater sensitivity to therapy.

To better understand why SGs showed prognostic potential only in these specific cancer types, we compared tumour samples from the two relevant GDC-TCGA cohorts with the remaining GDC-TCGA datasets(59, 60), while controlling for tissue of origin. This revealed strong upregulation of *MAMSTR* (Figure 7C), a MEF2 coactivator previously linked to EMT and improved survival(133, 134), alongside downregulation of proliferation-associated Hallmarks and upregulation of EMT and inflammation-related programmes (Figure 7D). The association between elevated inflammatory signalling and improved survival in high SG Score samples was unexpected, given the proposed role of SGs in attenuating inflammation(7). This raises the possibility that, at least in these cancers, this increased inflammation may be boosting anti-tumour immunity, thus impairing tumour growth (Figure 7D) (potential mechanisms reviewed elsewhere(135)).

To further investigate the basis of this association, we compared the top and bottom quartiles of SG Score within each of the two cancer types, while controlling for tissue of origin. DGE analysis identified upregulation of *hexokinase 2* (*HK2)* (Figure 7E), which binds VDAC to promote pore opening(136). While this typically facilitates ATP release(136), previous results(76) (Figure 5A) suggest it could also permit mitochondrial nucleic acid leakage, which we previously linked to SG assembly and inflammation (Figure 4 and 6). Consistent with this, GSEA revealed an apparent “inversion” relative to the basal state (Figure 7F): tumours with higher SG scores showed reduced EMT (previously upregulated) and enhanced proliferation-associated programmes (previously downregulated). However, increased interferon-α and -γ responses persisted, seemingly at odds with the idea that SGs suppress inflammation. One possibility is that we are not detecting a net anti-inflammatory effect but rather comparing tumours under stress with tumours under lesser amounts of stress. In this comparison, SGs may mitigate, but not eliminate, stress-induced inflammation.

## Discussion

SGs have long been purported to fine-tune the stress response by selectively sequestering specific proteins and transcripts, thereby protecting cells from stress-induced damage(12–15). Because cancer cells experience elevated stress, SGs have been proposed as contributors to tumour aggressiveness and therapeutic resistance(27, 28, 112). However, purified SG transcriptomes reported to date vary substantially across studies (Figure 1B), motivating our meta-analysis of published human SG transcriptomes.

Transcriptomes isolated by DC consistently contained longer RNAs than those obtained by other methods, even using the same cells and stressors (Figure 1B). *In silico* simulations of SG isolation, using empirical sedimentation values derived from yeast(33), reproduced this length bias (Figures 2D, 3B), supporting a technical origin. Centrifugation, being weight-dependent, preferentially enriches for longer/heavier RNA-containing condensates. Correcting for this technical signal reduced or eliminated the length bias in DC-derived transcriptomes (Figure 4A), although method-specific differences between studies remained (Figure 4A, Supplementary Figure 8D).

This correction also revealed that mitochondrial encoded transcripts sediment more than their length alone would predict (Figure 3A), a property we incorporated into our model. Across our corrected meta-analysis, mito-encoded transcripts showed no consistent enrichment or depletion in SGs across datasets, but they were more sensitive to differences between SGs and respective controls than nuclear-encoded mitochondrial genes. Mito-encoded transcripts were modestly but significantly enriched in our consensus SG signature, albeit not ranking among the most strongly enriched transcripts overall. Although this mtRNA signal was too modest and inconsistent across datasets to support a strong claim of mtRNA enrichment in SGs on its own, the specific sensitivity of those transcripts motivated us to test directly whether SGs sequester mitochondrial dsRNA.

We next compared stressed cells with and without SGs, observing an apparent depletion of inflammation-related pathways in cells containing SGs (Supplementary Figure 8). These results are consistent with a model in which SGs dampen the stress response, in line with previous work linking dsRNA sensing to inflammatory signalling(7), and extend this observation to oxidative and hyperosmotic stress.

Mitochondrial nucleic acids act as damage-associated molecular patterns that trigger inflammation(94). We hypothesised that SGs may sequester mitochondrial dsRNA, limiting its release and downstream activation of MDA5/RIG-I signalling. Consistent with this, inhibiting VDAC pores with DIDS, previously shown to block mitochondrial dsRNA efflux(76), abolished SG assembly (Supplementary Figure 11), indicating that permeability of mitochondria is required for SG formation, although we cannot rule out nonspecific effects of DIDS or contributions from other mitochondrial factors.

High-resolution Airyscan imaging revealed that SG-like foci, readily distinguishable by conventional confocal microscopy, became harder to resolve from the broader G3BP1 signal (Supplementary Figure 11 vs Figure 6A), highlighting a broader definitional challenge. SGs are identified almost entirely by size-based imaging criteria that likely underestimate their presence under moderate stress. Within this imaging framework, dsRNA showed greater colocalisation with G3BP1-positive foci under hyperosmotic stress than at baseline (Figure 6B–D). We interpret this as supportive of, rather than definitive evidence for, our model, as the J2 antibody detects dsRNA broadly and we cannot exclude non-mitochondrial sources. The convergent evidence from the VDAC perturbation and imaging experiments nonetheless argues for a mitochondria-derived origin.

We also found that TIA1-regulated splicing events in SG-containing cells resembled those in TIA1 knockdown cells (Supplementary Figure 12), consistent with substantial TIA1 sequestration into SGs during stress. However, we cannot distinguish a direct functional consequence of SG assembly from a parallel effect, as SG assembly itself depends on G3BP1/G3BP2. Immunofluorescence showed predominantly cytoplasmic TIA1 even in unstressed cells (Supplementary Figure 6), which is not fully consistent with the canonical model in which stress drives TIA1 to the cytoplasm for SG sequestration(105, 106). We do not currently have an explanation for this observation and flag it as an open question.

Because SG-associated genetic models typically involve proteins with functions beyond SG formation, disentangling SGs’ contribution to tumour aggressiveness from these pleiotropic effects is difficult(27, 28, 112). Using the null hypothesis that these additional functions explain the reported SG–aggressiveness link, our analysis could not reject this alternative explanation (Supplementary Figures 14, 15), indicating that single-gene knockout approaches are insufficient to isolate SGs’ specific contribution.

As an orthogonal approach, we developed an SG gene-expression signature (SG-Score) via elastic net regression, validated in an independent dataset (Figure 7A, B), and applied it to GDC-TCGA tumour transcriptomes(59, 60). Elevated SG-Score in tumour samples was restricted to a subset of cancer types, arguing against a universal increase in SG formation across cancers (Supplementary Figure 16) and consistent with SG formation requiring stress conditions more severe than typical physiological ones(32). Survival analysis, benchmarked against 100 randomly generated signatures, identified two cancer types (ACC, MESO) where lower SG-Score was associated with poorer outcomes, opposite to the expectation that SG assembly promotes aggressiveness. In these cancers, higher SG-Score correlated with higher proliferation and sustained interferon responses (Supplementary Figure 14B/C), suggesting SGs may mitigate but not fully suppress stress-induced inflammation, with implications for the balance between tumour progression and immune activation.

A further notable finding was upregulation of *HK2* in SG-Score-high ACC and MESO tumour samples (Figure 7E), a modulator of VDAC activity and mitochondria-to-cytosol ATP transport(136). Given VDAC’s role in mitochondrial dsRNA efflux (76), we hypothesise that this axis could link SG formation to inflammatory signalling in these cancers. We present this explicitly as a hypothesis for future functional validation.

This study has several limitations:

- Analysis of published SG transcriptomic datasets was complicated by dataset-specific technical issues, including sample mislabelling, limited specificity of SG-purification techniques, and inadequate gene filtration (see Supplementary Methods).
- Our correction for centrifugation-associated length bias relied on sedimentation parameters derived from yeast, which may not fully capture human mitochondrial RNA dynamics.
- As discussed above, SGs are identified almost entirely by size-based imaging criteria; image-processing choices for SG detection are, moreover, not consistently reported in the field, which complicates cross-study comparison.
- Our dsRNA–SG colocalisation and TIA1-splicing analyses were correlational and based on limited sample sizes.
- Multiple SG subtypes have been described based on their dependence on eIF2á phosphorylation, and differ in protein composition(137). Our analyses did not distinguish between subtypes, which may contribute to variability across datasets. Our differential expression analyses used linear models assuming additive, constant effects, which may not capture non-linear, threshold-dependent, or epistatic regulatory relationships.
- Finally, the prognostic value of the SG-Score was identified in a small subset of cancer types and requires independent, prospective validation before translational application.

We nonetheless identify recurring technical sources of variability across published SG transcriptomes and propose a corrected, consolidated SG transcriptome alongside the transcriptomic changes accompanying SG assembly. SG formation was associated with downregulation of stress- and inflammation-related pathways, suggesting a stressor-independent protective response, and our data support a model in which SGs modulate inflammatory signalling via sequestration of mitochondrial dsRNA, with translational relevance suggested by the SG-Score’s behaviour in a subset of cancers.

## Supplementary Data

Supplementary data are available.

## Data Availability

All analysis code is available in a public Zenodo repository (https://doi.org/10.5281/zenodo.21724124) and in a public GitHub repository (https://github.com/DiseaseTranscriptomicsLab/StressGranules), organised into: *Datasets* (analysis of each dataset referenced under “Datasets used”), *Files* (including the SG Score signature and the WC and SG meta-analysed profiles), *Scripts* (bash scripts for preprocessing and analysis), and *Simulations* (files and scripts for reproducing Mimi-Seq and Spin-Sim simulations).

The eCLIPSE pipeline is still under active development and as-yet unpublished. However, components necessary to re-create the analysis, and related work are available in separate repositories in Zenodo (https://doi.org/10.5281/zenodo.21724194), and in GitHub (https://github.com/DiseaseTranscriptomicsLab/CHARM: Comprehensive Hub for Alternative Regulatory Mapping), with the full pipeline provided upon request.

## Acknowledgments

We thank all members of the Disease Transcriptomics Lab for providing valuable feedback on the manuscript, in particular Rita Martins-Silva and Francisca Xara-Brasil. We thank all members of the Mitochondria Biology & Neurodegeneration Lab for support during laboratory experiments. We thank the technical support of the Bioimaging Platform of the Gulbenkian Institute for Molecular Medicine, funded by PPBI-POCI-01-0145-FEDER-022122. We are also grateful to the Maria Carmo-Fonseca Lab for providing U2-OS cells and to the Leonor Saúde Lab for providing Sorbitol.

Graphical abstract was created in BioRender (https://BioRender.com/oblsxww). This work includes the use of generative artificial intelligence tools, specifically ChatGPT (OpenAI, GPT-4 and GPT 5.5) and Claude (Anthropic, Sonnet 4.6), to assist with language editing, text refinement, and code troubleshooting. These tools were used under the lead author’s supervision, and all generated content was critically reviewed and verified for accuracy and appropriateness. We confirm that the intellectual content, interpretation, and conclusions presented in this work are our own, with AI used solely as a supporting tool, and we take full responsibility for the content of the published article.

Further information and requests for resources, data, and/or code should be directed to, and will be fulfilled by the lead corresponding author, Alexandre Kaizeler.

## Author contributions

A.K. conceived the project; A.K., V.A.M., and N.L.B.-M. designed the experiments; A.K. wrote the manuscript, with contributions from all the other authors; A.K. performed all experimental work; A.K, D.C and D.V.V performed image acquisition, with support of the Bioimaging Platform of the Gulbenkian Institute for Molecular Medicine; D.C performed image analysis; A.K. performed all other data analyses.

