## Supplemental File for "Revisiting Stress Granule Transcriptomes Suggests Mitochondrial RNA Enrichment Despite Methodological Bias"

Alexandre Kaizeler[1,2,3]*

Diogo Coutinho[1]

Diana V. Vieira[1]

Vanessa A. Morais[1,2,]**

Nuno L. Barbosa-Morais[1,2,3]***

[1] GIMM - Gulbenkian Institute for Molecular Medicine, 1649-035 Lisbon, Portugal

[2] Faculdade de Medicina, Universidade de Lisboa, 1649-028 Lisbon, Portugal

[3] Comprehensive Health Research Centre (CHRC), NOVA Medical School|Faculdade de Ciências Médicas, NMS|FCM, Universidade NOVA de Lisboa, 1169-056 Lisbon, Portugal

### Supplementary Data

### Materials and Methods

#### Sampling bias simulations (Mimi-Seq)

To assess potential sampling bias, we simulated RNA-seq fragmentation using expression and fragment length distributions derived from GSE99304(1) WC and SG samples. For each sample, paired-end alignments were used to extract two metrics from BAM files:

1. the observed template length (TLEN) field, which reports the observed fragment size;
2. the median fragment position, calculated for each read pair as:

$$\frac{start\ (read_1)+start\ (read_2)+read\ length}{2} \quad (1)$$

For each fragmentation event, fragment lengths were randomly sampled from the corresponding empirical size distribution, and transcripts were iteratively fragmented until no additional fragment could be accommodated within the remaining sequence. This approach preserved both transcript identity and fragment length characteristics in the simulated reads.

Following simulation, transcript abundances were converted back to discrete read counts to generate sequencing-like count matrices, allowing downstream differential expression analyses, while preserving the intended total library depth of empirical SG datasets. Three simulated WC and three simulated SG samples were generated to assess whether subsampling to approximately 10% of total reads, mimicking the SG fraction, was sufficient to induce length-dependent transcript representation biases.

#### Dataset-specific caveats and exclusion criteria

For the full list of datasets used throughout this work (see Table 1; Figure 1B), please refer to the main publication's Materials and Methods sub-section "Datasets".

For the final set of SG and WC transcriptomes corrected for centrifugation-associated RNA length biases (Figure 4A), studies were maintained if they had appropriate technical controls (*i.e.*, not just SG samples, but also equivalent cellular fractions, empty vectors, in stressed and unstressed conditions) and at least 3 SG replicate samples. Even though most DC studies had no appropriate technical controls, we considered that our simulated SG samples could serve as such, and thus these studies were maintained.

A more detailed assessment of some of the datasets, as well as their reason for exclusion/inclusion, is provided below:

- **Matheny 2021 (GSE119977**(2, 3)**)**: In the course of our reanalysis, a subset of samples labelled as G3BP KO appeared, by expression profile, more consistent with G3BP WT. We raised this with the original authors, who confirmed and corrected the labelling(3); the corrected dataset was used in our analyses.
- **Padron 2019 (GSE121575**(4)**)**: This study was performed in HEK293 cells, which have a female karyotype. Several Y-chromosome-specific proteins (*e.g.*, DDX3Y) were nonetheless reported as being especially present within SG (4). Additionally, we found limited replicate concordance under stress conditions (Supplementary Figure 19A) and, given that only two replicates were available, this dataset was excluded from our analyses.
- **Honda 2021 (GSE143413**(5)**)**: As only one SG sample was available, this dataset was excluded.
- **Sato 2023 (GSE188397**(6)**)**: In several conditions, stressed replicates correlated more closely with unstressed samples than with each other (Supplementary Figure 19B, G3BP1 interactome as example). As only two replicates were available per condition, this dataset was excluded.
- **Xiao 2023 (GSE226161**(7)**)**: Several replicates exhibit limited correlation with one another (Supplementary Figure 19C). As only two replicates were available per condition, this dataset was excluded.
- **Ren 2023 (GSE223295**(8)**)**: Although four putative SG samples were available, these corresponded to two distinct stressors (arsenite and sorbitol, two replicates each) rather than three or more replicates of a single condition. A triplicate sample group, collected one hour after stress removal, was reported as representing a recovery-phase SG state; however, as no other dataset in our meta-analysis included a comparable recovery, we opted to exclude the dataset from our analyses.
- **Shen 2022 (GSE171792**(9)**)**: After filtering lowly expressed genes, the two "Stressed APEX2-GFP" replicates showed lower concordance with each other than other replicate pairs in the same dataset, as well as accounting for most of the variability within the dataset (Supplementary 19 D, E). With only two replicates available, this dataset was excluded.
- **Zhou 2024 (GSE218180**(10)**)**: The three SG replicates in this dataset were concordant. However, the study design did not include a technical control (*e.g.*, the same purification protocol applied to a non-SG structure), which would be needed to separate a biological SG signal from effects of the purification technique itself. Without this control, we were unable to confidently attribute the observed signal to SG biology, and this dataset was excluded.

- **Rajachandran 2024 (PRJNA1152621**(11)**):** This study included three SG replicates alongside two nucleolus samples, profiled using the same purification technique. As the nucleolus samples were generated under an identical technical protocol applied to a different cellular structure, we considered them an adequate technical control, allowing us to distinguish SG-specific biological signal from effects of the purification method itself. This dataset was therefore retained for our analyses.
- **Iadevaia 2022 (GSE171655**(12)**)**: In this dataset, SG-designated samples correlated more closely with unstressed whole-cell (Total) samples than with stressed whole-cell samples (Supplementary Figure 19F–G), differing from the pattern observed in other DC-purified datasets. To further investigate this, we generated matched simulated samples using our SpinSim approach; these exhibited a markedly stronger RNA length bias than simulated samples derived from other DC studies (Supplementary Figure 19H), suggesting the isolated structures may differ from canonical SGs in composition. This is consistent with the original study's own characterisation of these structures as a distinct entity, termed paracrine granules, rather than canonical SGs(12). Given this, we considered the dataset outside the scope of our SG meta-analysis and excluded it from our analyses.

Analyses of all datasets (both retained and excluded) are available in a public Zenodo repository (https://doi.org/10.5281/zenodo.21724124), as well as in a public GitHub repository (https://github.com/DiseaseTranscriptomicsLab/StressGranules). Analyses are available within the Datasets folder, under the corresponding GSE ID.

### Results

#### Sampling bias does not explain the RNA length bias in DC-derived SGs

Since SGs contain only about 10% of the total cellular mRNA(1), the RNAs detected in SGs may not fully represent the diversity of the whole-cell transcriptome. In small mRNA samples, longer transcripts may be overrepresented. This occurs because the RNA-seq library preparation process involves fragmentation, which yields more short sequencing reads per full molecule for longer transcripts. Standard normalisation methods that account for transcript length do not fully correct for this overrepresentation bias(13, 14).

To formally assess whether technical or sampling effects could contribute to a length bias, we performed *in silico* RNA sequencing (RNA-seq) simulations using empirical RNA expression levels and fragment length distributions from the original SG and WC samples(1) (Supplementary Figures 2, 3A). Additional simulations, in which expression data from one population were paired with the alternate fragmentation profile, demonstrated that fragmentation patterns depended only on the input expression, confirming equivalence of SG and WC fragmentation profiles (Supplementary Figure 3B). Finally, we simulated WC and SG samples by drawing reads from the full WC transcriptome, with SG samples representing a 10% subset of that expression. These simulated SG transcriptomes exhibited a negative length bias, in contrast to the positive bias in DC-purified SG transcriptomes (Supplementary Figure 3C). This reversal is most likely due to the high abundance of short RN7SL1–3 RNAs in the original WC transcriptome, which constitute approximately 50% of total reads(1), and outweigh contributions from longer transcripts in the simulation.

Even in this simplified scenario, designed to maximise technical enrichment of longer RNAs, the positive length bias in SGs was not reproduced. In fact, library complexity and sequencing depth of SGs in the original study(1) appear sufficient to prevent the hypothesised sampling bias (Supplementary Figure 3D).

Collectively, these results suggest that the positive length bias in DC-purified SG transcriptomes is unlikely to arise from either sampling bias or other RNA-seq technical effects owing to the lower number of RNA molecules in SGs, when compared to the remainder of the cell.

### Supplementary Figures

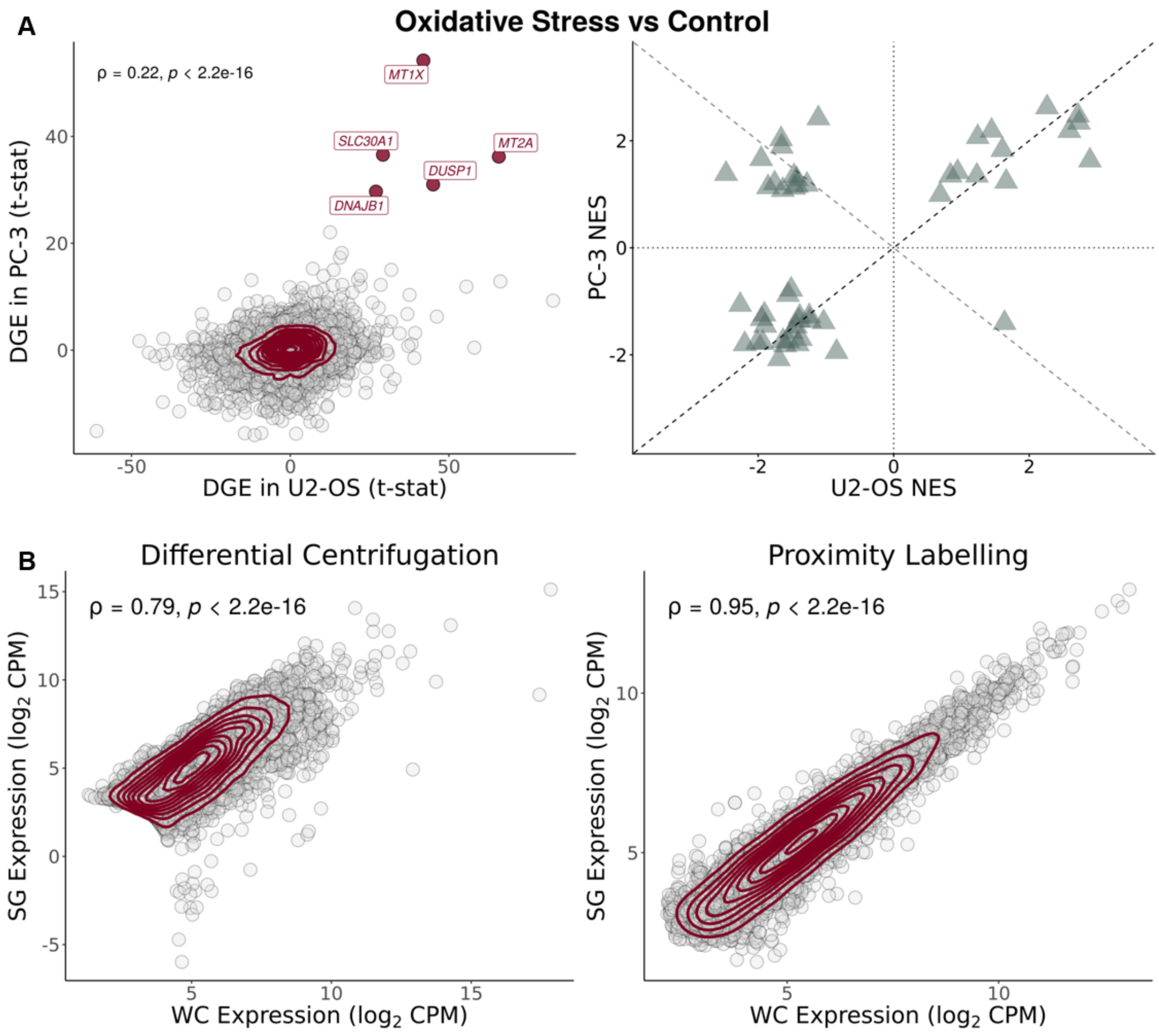


**Supplementary Figure 1. U2-OS and PC-3 cells have a similar stress response.** (A) (left) Scatter plot of t-statistics of DGE between SGs and WCs in PC-3(15) (Y-Axis) and U2-OS(16) (X-Axis) cell lines exposed to oxidative stress through arsenite; genes as circles, density contour lines in dark red, Spearman's correlation coefficient ρ and associated test's p-value on top left corner; strongly upregulated genes in both cell lines (t-statistic > 25) are labelled and highlighted in dark red. (right) Scatter plot of GSEA NES of Hallmarks, ranked by t-statistics of DGE, in PC-3(15) (Y-axis) *versus* U2-OS(16) (X-axis) cell lines, exposed to oxidative stress through arsenite. Dashed lines depict identity (Y = X; Y = -X). (B) Scatter plots of gene expression (in $\log_2$ counts-per-million (CPM)) in SGs (Y-Axis) *versus* WCs (X-Axis), for SG purification by DC(1) (left) and PL(15) (right); genes as circles, density contour lines in dark red, Spearman's correlation coefficient ρ and associated test's p-value on top left corner.

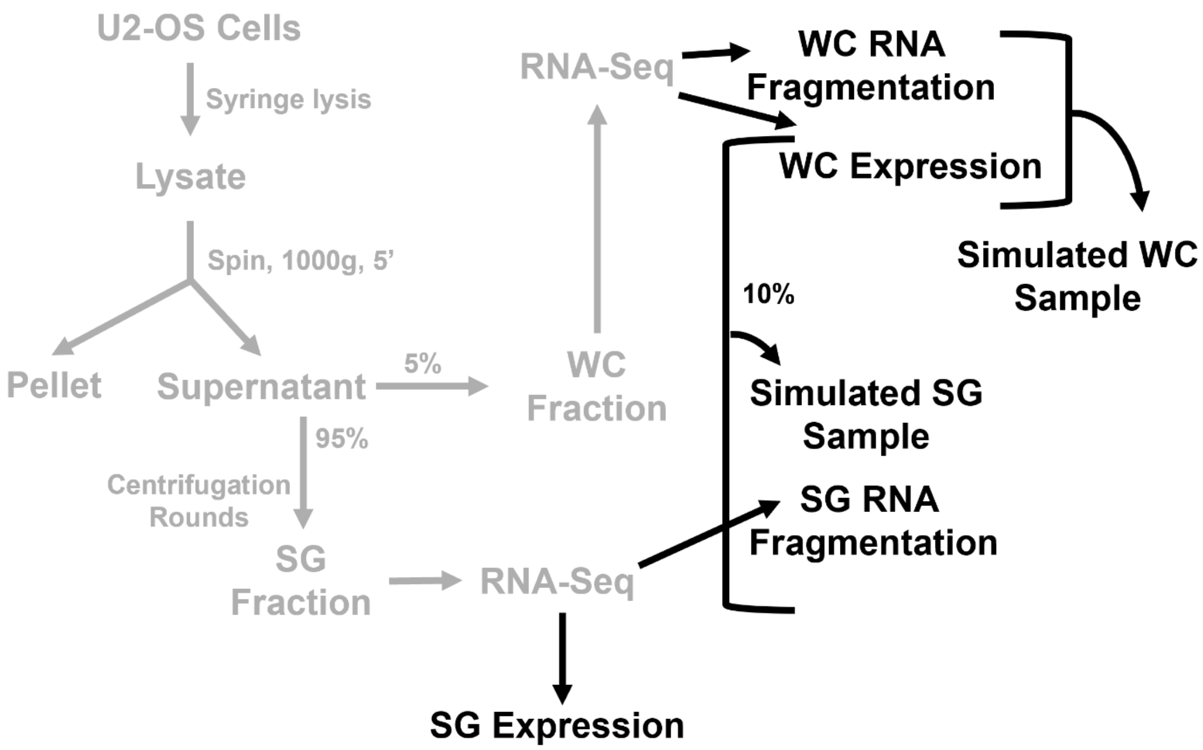


**Supplementary Figure 2. Sampling bias simulation pipeline.** The diagram depicts the experimental workflow from the original study for obtaining WC and SG samples(1) (steps in grey) and the additional in silico steps performed in our analyses (steps in black).

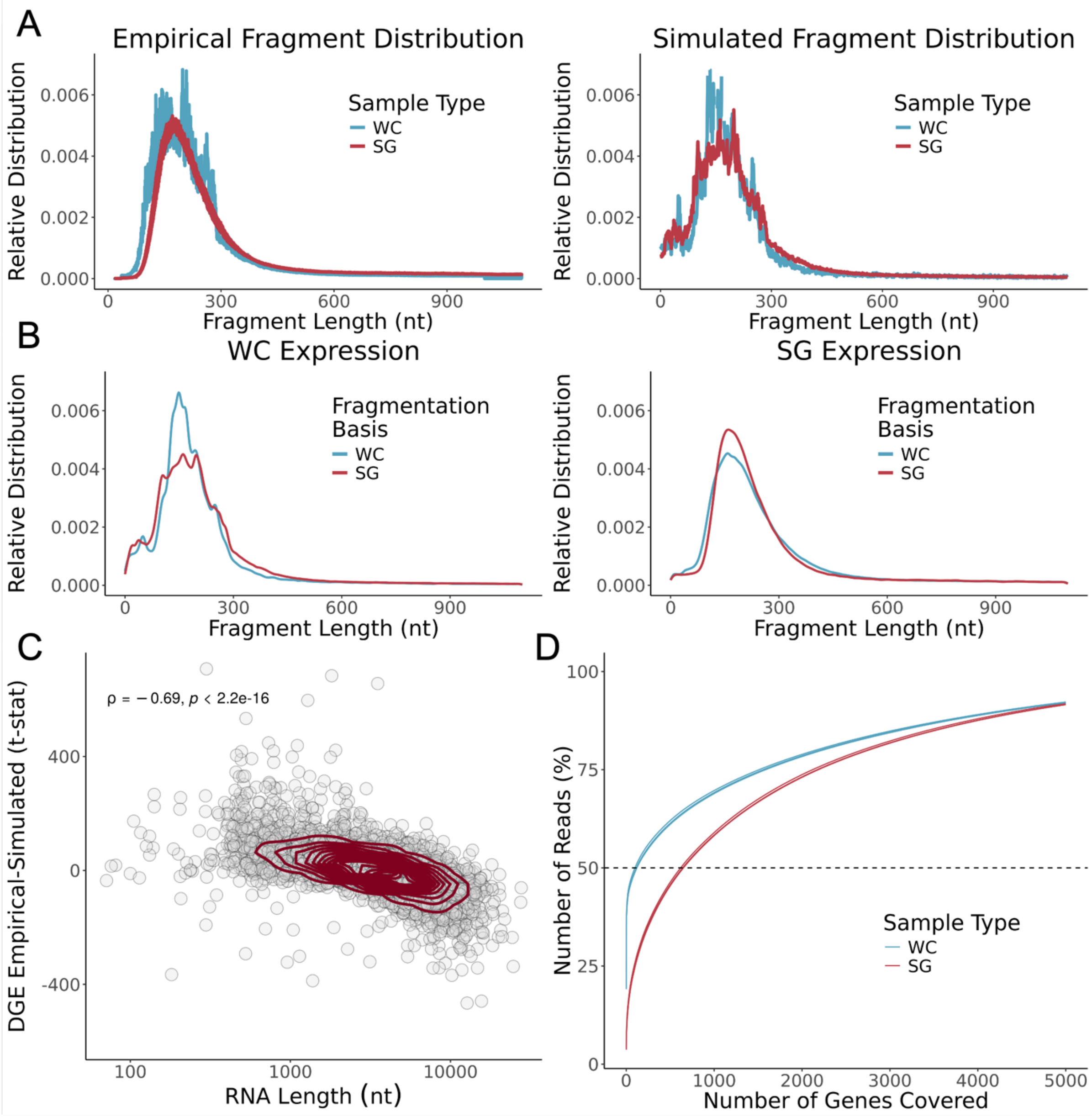


**Supplementary Figure 3. Transcriptomes of DC-isolated SGs show no evidence for RNA fragmentation-associated sampling bias.** (A) Relative distribution (density) of fragment length (nt) in original samples [28] RNA-seq (Left) and simulated samples (Right). (B) Relative distribution (density) of representative RNA fragment length (nt) in simulated samples, using WC (Left) or SG (Right) expression as a basis, and WC or SG fragmentation distributions as a basis. (C) Scatter plot of t-statistics of DGEA between SG and WC simulated samples (Y-Axis) and representative RNA transcript length (nt) (X-Axis, log scale); genes as circles, density contour lines in dark red, Spearman's correlation coefficient ρ and associated test's p-value on the top left corner. The original WC expression(1) was used as the basis for both simulated WC and SG samples, with their respective fragment size distribution as the basis. (D) Cumulative distribution of the percentage of RNA-seq reads mapping to the increasing number of genes, ordered by decreasing read coverage, in the original dataset(1). More genes account for 50% of the reads in SG samples than their WC counterparts, suggesting greater RNA library complexity.

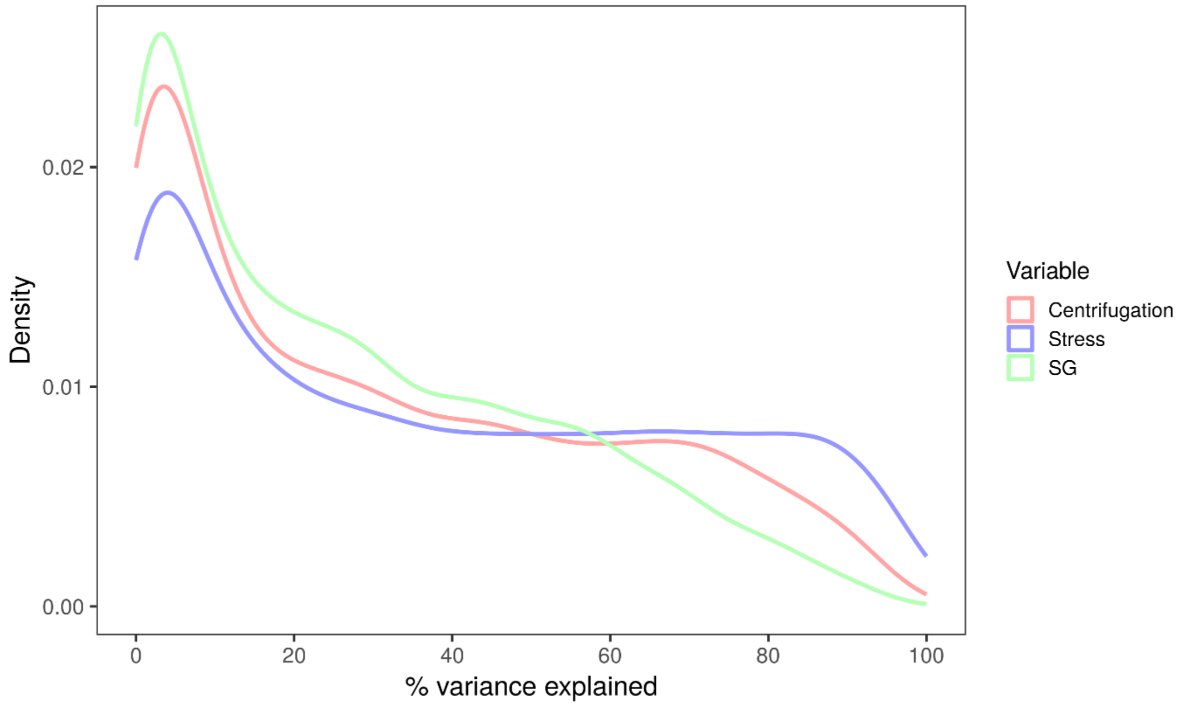


**Supplementary Figure 4**. **Relative contributions of different sources to gene expression variance in the Matheny 2019**(16) **dataset**. SG contributes less to gene expression variance than the stress itself or the centrifugation effect.

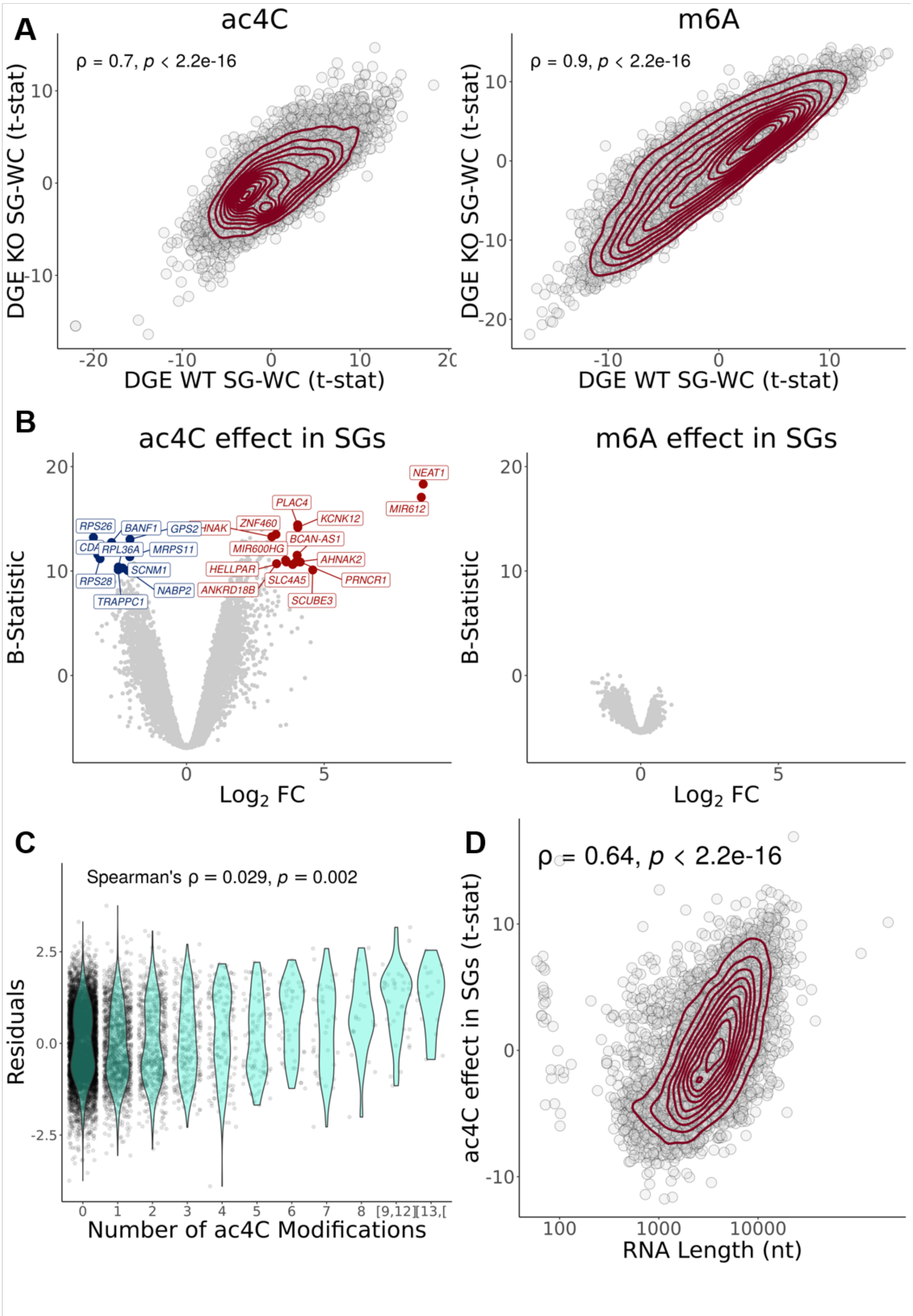


**Supplementary Figure 5. ac4C, but not m6A, is associated with altered SG transcript enrichment, driven by RNA length.** (A) Scatter plots of t-statistics of DGE in WT SGs *versus* WCs (Y-axis) and in SGs *versus* WCs in cells without RNA modifications (X-axis) - ac4C KO(17) (Left) and m6A KO(18) cells

(Right); transcripts as circles, density contour lines in dark red, Spearman's correlation coefficient ρ and associated test's p-value on the top left corner. (B) Volcano plots of the transcriptomic effect (given by linear modelling with interaction) of ac4C (Left) and m6A (Right) modifications in SG samples. (C) Violin plots of residuals (the difference between the actual value and the value predicted by the linear model applied for any given transcript) as a function of the number of ac4C modifications per transcript; Spearman's correlation coefficient ρ and associated test's p-value shown on the top left corner. (D) Scatter plot of residuals (Y-axis) and transcript length (nt) (X-axis); transcripts as circles, density contour lines in dark red, Spearman's correlation coefficient ρ and associated test's p-value shown on the top left corner. For all panels concerning ac4C, the GSE212380 dataset was used(17). For those concerning m6A, the GSE242766 dataset was used(18).

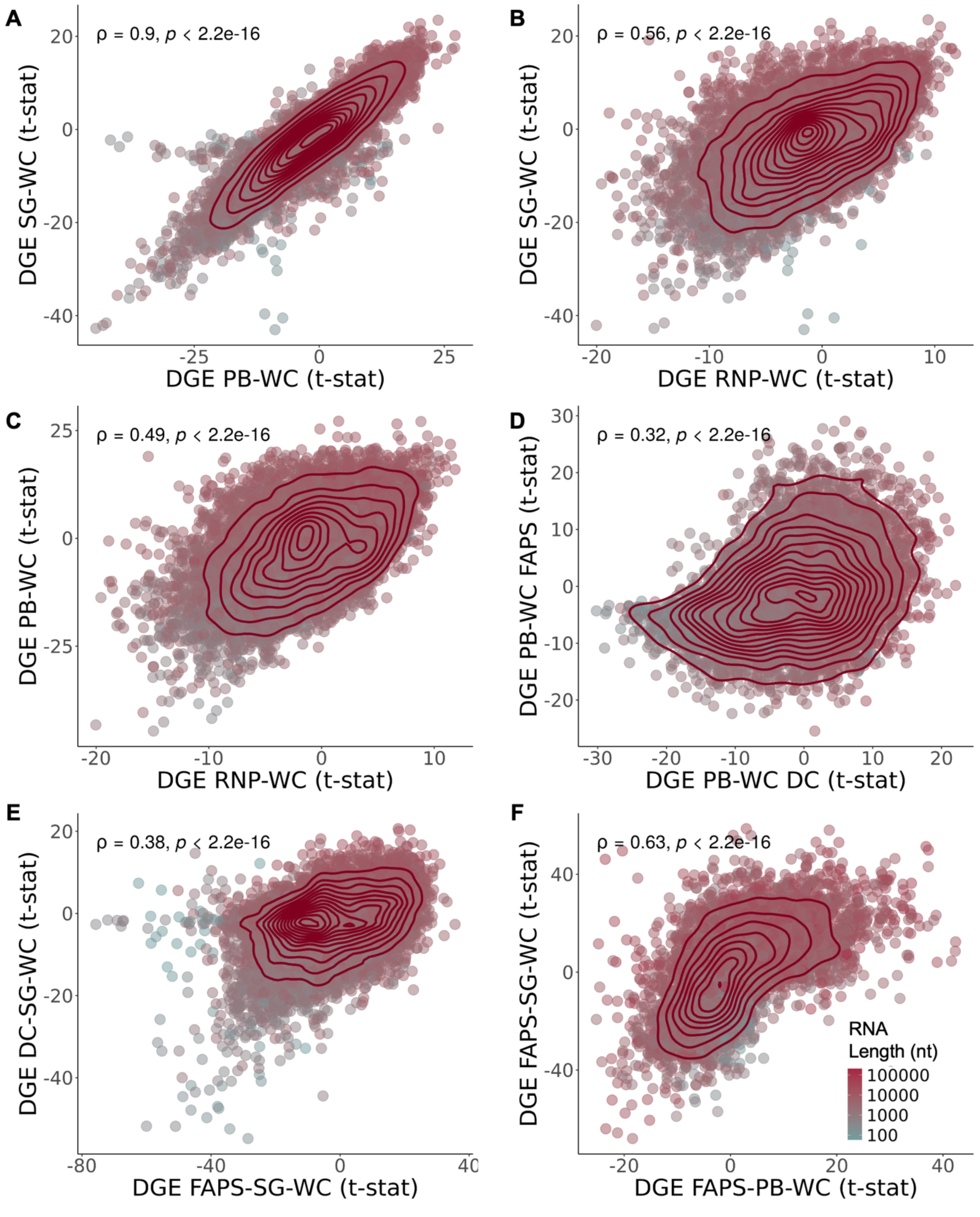


**Supplementary Figure 6. Samples isolated through DC are highly correlated.** (A) Scatter plot of t-statistics of DGE in SGs *versus* WCs(1) (Y-Axis) and in PBs *versus* WCs(16) (X-Axis). (B) Scatter plot of t-statistics of DGE in SGs *versus* WCs(1) (Y-Axis) and in RNP granules *versus* WCs(16) (X-Axis). (C) Scatter plot of t-statistics of DGE in PBs *versus* WCs(16) (Y-Axis) and in RNP granules *versus* WCs(16) (X-Axis). (D) Scatter plot of t-statistics of DGE in PBs purified by FAPS(19) (Y-Axis) and by DC(16), compared to WCs (X-Axis). (E) Scatter plot of t-statistics of DGE in SGs purified by DC(1) (Y-Axis) and by FAPS/FANCI(10) (X-Axis), compared to WCs, after batch correction for cell type and stress. (F) Scatter plot of t-statistics of DGEA in SG (Y-Axis) and PB (X-Axis) purified by FAPS/FANCI(10, 19) compared to WCs, after batch correction for stress. In all panels, genes as circles coloured by RNA length (nt); density contour lines in dark red, Spearman's correlation coefficient ρ and associated test's p-value on top left corner.

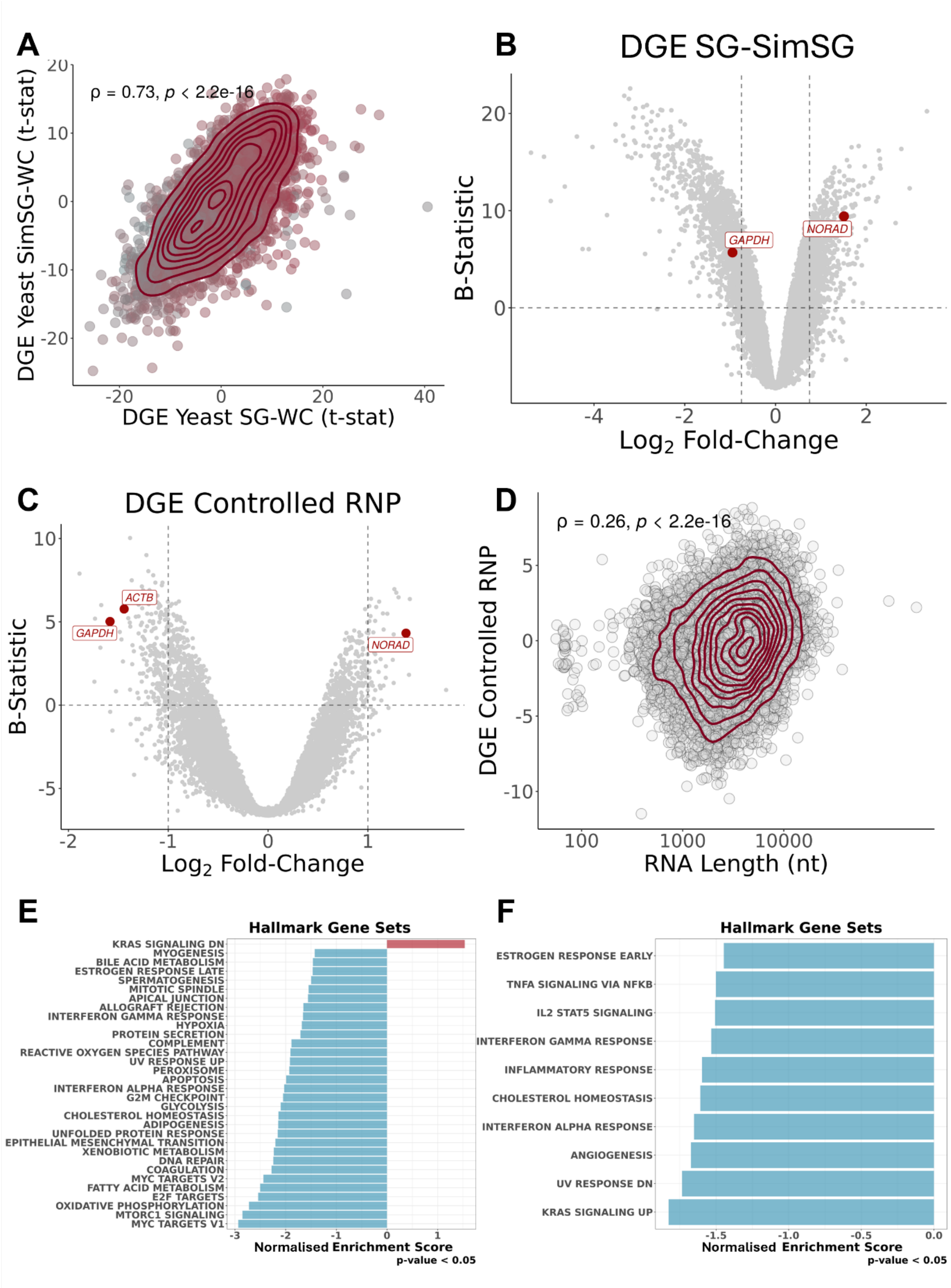


**Supplementary Figure 7.** (A) Scatter plot of t-statistics of DGE of simulated yeast SG *versus* WC (Y-Axis), with *S* estimations based on Sed-seq data[31], and t-statistics of DGE in yeast SGs purified by DC[29]

(X-Axis); transcripts as circles coloured by RNA length; density contour lines in dark red, Spearman's correlation coefficient ρ and associated test's p-value on top left corner. (B) Volcano plot of DGEA (Y-Axis – B-Statistic; X-Axis - log2 Fold-Change) between original SGs(1) and simulated SGs, based on Sed-Seq(20) data; transcripts as circles, with commonly used RNA FISH controls(1) in red. Vertical dashed lines correspond to -0.75 and 0.75 log 2FC. Horizontal dashed line corresponds to B-statistic = 0. (C) Volcano plot of DGEA (Y-Axis – B-Statistic; X-Axis - Log2 Fold-Change) of interception of stress with RNP sample types, given by linear modelling (*i.e.*, corresponding to DC-isolated RNP granules stress-specific sequestration)(16); transcripts as circles, with commonly used RNA FISH controls(1) in red. Vertical dashed lines correspond to -1 and 1 log2FC. Horizontal dashed line corresponds to B-statistic = 0. (D) Scatter plot of t-statistics of DGE in linear modelled DC-isolated RNP granules stress-specific sequestration(16) (Y-Axis) and RNA length (X-Axis); transcripts as circles; density contour lines in dark red, Spearman's correlation coefficient ρ and associated test's p-value on top left corner. (E) GSEA NES of Hallmark gene sets, with genes ranked by t-statistics of differential expression, in linear modelled DC-isolated RNP granules stress-specific sequestration(16). (F) GSEA NES of Hallmark gene sets, with genes ranked by t-statistics of differential expression, in SG-bearing WC samples *versus* their Non-SG-bearing counterparts(2, 16).

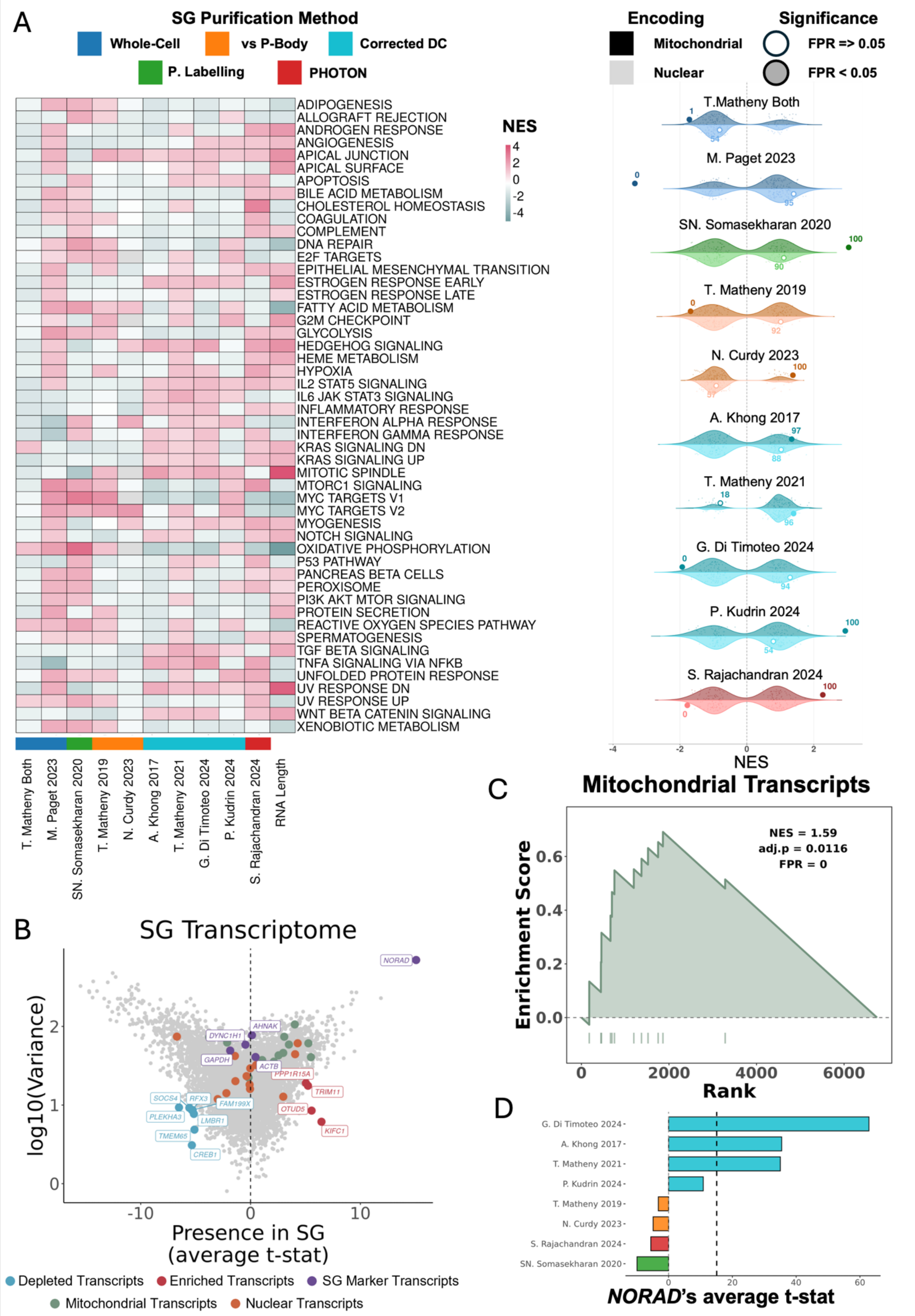


**Supplementary Figure 8. Mitochondrial RNA enrichment across SG transcriptomes.** (A) (Left) Heatmap of NES from GSEA using Hallmark gene sets, based on genes ranked by t-statistics of DGE between centrifugation-corrected SGs and controls or between WCs with and without SGs across multiple studies. (Right) Paired half-violin plots showing the distribution of NES values obtained from GSEA using 1000 size-matched random gene sets (violins), for mitochondrially encoded (darker shades) or nuclear-encoded mitochondrially localised genes (lighter shades), across the transcriptomic profiles of corrected/controlled SGs from different studies. The observed NES for each gene set is shown as a large dot; filled dots indicate empirical FPR < 0.05. Numbers adjacent to each dot indicate the percentile of the observed NES within the corresponding null distribution. Colour of each violin indicates SG purification method, as indicated above the image. (B) Cross-dataset variance of t-statistic of DGE between centrifugation-corrected SGs and controls plotted against its average across datasets (see Methods); transcripts as circles, coloured green if mitochondrial encoded, orange if nuclear encoded and mitochondrial located, purple if used as SG RNA markers(1), blue if "consistently depleted" (Variance t < 10 and Average t < -5), and red if "consistently enriched" (Variance t < 20 and Average t > 4.5). (C) Enrichment plot of GSEA of mitochondrially encoded transcripts ranked by the cross-dataset average t-statistic of DGE between centrifugation-corrected SGs and controls. (D) Bar plot of *NORAD's* average t-statistic of DGE between centrifugation-corrected SGs and controls across different studies. Average value is represented by the dashed black line.

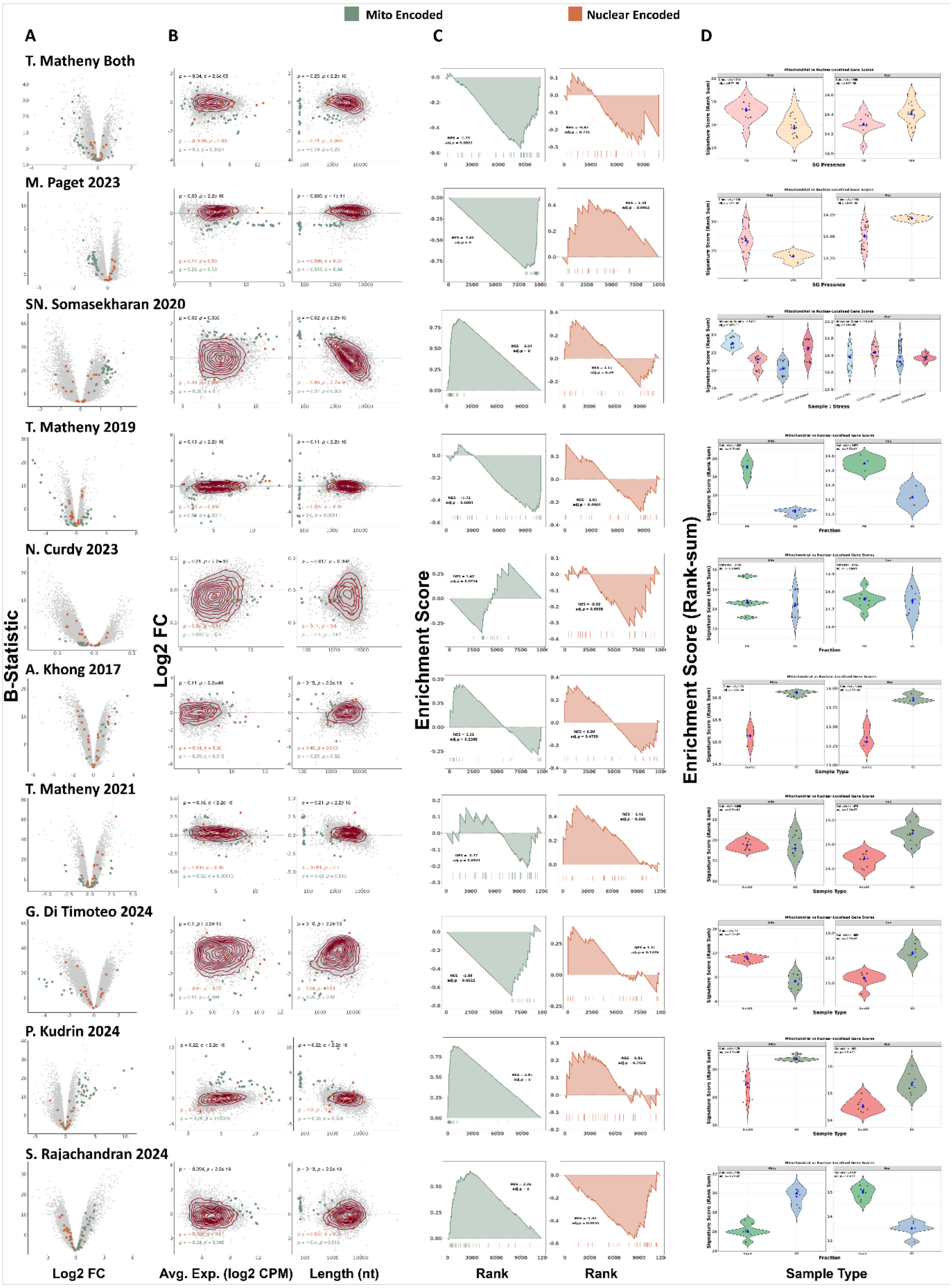


**Supplementary Figure 9.** Multi-panel differential expression and gene set enrichment summary for mitochondrially-encoded and nuclear-encoded mitochondria-localised transcripts in SG transcriptome studies. (A) Volcano plots of DGE between centrifugation-corrected SGs and controls or between WCs with and without SGs (Y-Axis – B-Statistic; X-Axis - log2 Fold-Change) per study; transcripts shown as grey circles, with mitochondrially-encoded transcripts highlighted in green and nuclear-encoded mitochondria-localised transcripts highlighted in orange. (B-Left) MA plots (Y-Axis – B-

Statistic; X-Axis – Average transcript expression) per study; transcripts shown as grey circles, with mitochondrially-encoded transcripts highlighted in green and nuclear-encoded mitochondria-localised transcripts highlighted in orange; density contour lines in dark red; Spearman's correlation coefficient ρ and associated test's p-value shown for all transcripts (black), the nuclear-encoded subset (orange), and the mitochondrially-encoded subset (green). (B-Right) Scatter plots of log2 Fold-Change of DGE between centrifugation-corrected SGs and controls or between WCs with and without SGs (Y-Axis) and RNA Length (X-Axis) per study; transcripts shown as grey circles, with mitochondrially-encoded transcripts highlighted in green and nuclear-encoded mitochondria-localised transcripts highlighted in orange; density contour lines in dark red; Spearman's correlation coefficient ρ and associated test's p-value shown for all transcripts (black), the nuclear-encoded subset (orange), and the mitochondrially-encoded subset (green). (C) GSEA enrichment plots for mitochondrially-encoded (green) and nuclear-encoded mitochondria-localised (orange) gene sets, respectively, with genes ranked by t-statistics of DGE between centrifugation-corrected SGs and controls or between WCs with and without SGs across multiple studies; NES and Benjamini-Hochberg (BH)-adjusted p-values are shown for each. (D) Violin plots of mitochondrial and nuclear-encoded gene signature scores (Rank-sum, see Methods) by sample type (defined on a study-study basis, see Methods); Cohen's f and BH-adjusted p-value from one-way ANOVA are shown for each panel. Blue diamonds and triangles indicate group means and medians, respectively. Violin colours are related to specific sample types within each SG-purification method (see Methods). Across the image, every row corresponds to an individual study, as labelled.

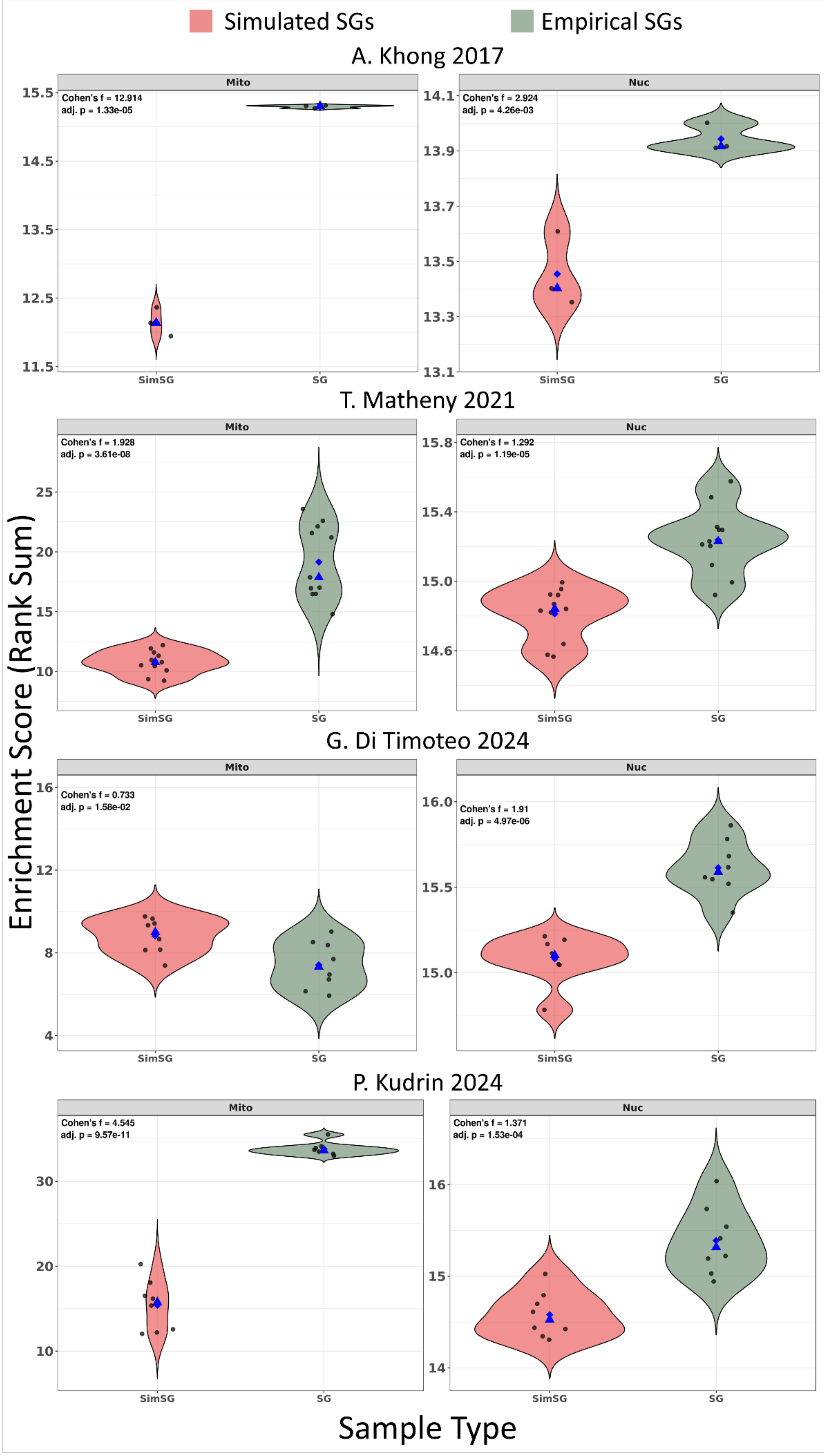

Simulated SGs
Empirical SGs
A. Khong 2017
Mito
Nuc
Cohen's f = 12.914
adj. p = 1.33e-05
Cohen's f = 2.924
adj. p = 4.26e-03
T. Matheny 2021
Cohen's f = 1.928
adj. p = 3.61e-08
Cohen's f = 1.292
adj. p = 1.19e-05
G. Di Timoteo 2024
Cohen's f = 0.733
adj. p = 1.58e-02
Cohen's f = 1.91
adj. p = 4.97e-06
P. Kudrin 2024
Cohen's f = 4.545
adj. p = 9.57e-11
Cohen's f = 1.371
adj. p = 1.53e-04
SimSG
SG
Enrichment Score (Rank Sum)
Sample Type

**Supplementary Figure 10.** Violin plots of mitochondrial-encoded (Mito) and nuclear-encoded mitochondria-localised (Nuc) gene signature scores in empirical (SG) versus simulated (SimSG) stress granule samples, across DC studies. For each sample, all genes were rank-ordered by expression, and a signature score was computed as the summed rank of genes in each set, normalised to the total number of expressed genes (see Methods). Effect size (Cohen's f, derived from one-way ANOVA) and BH-adjusted p-value are shown in the top-left corner of each panel. Blue diamonds and triangles indicate group means and medians, respectively.

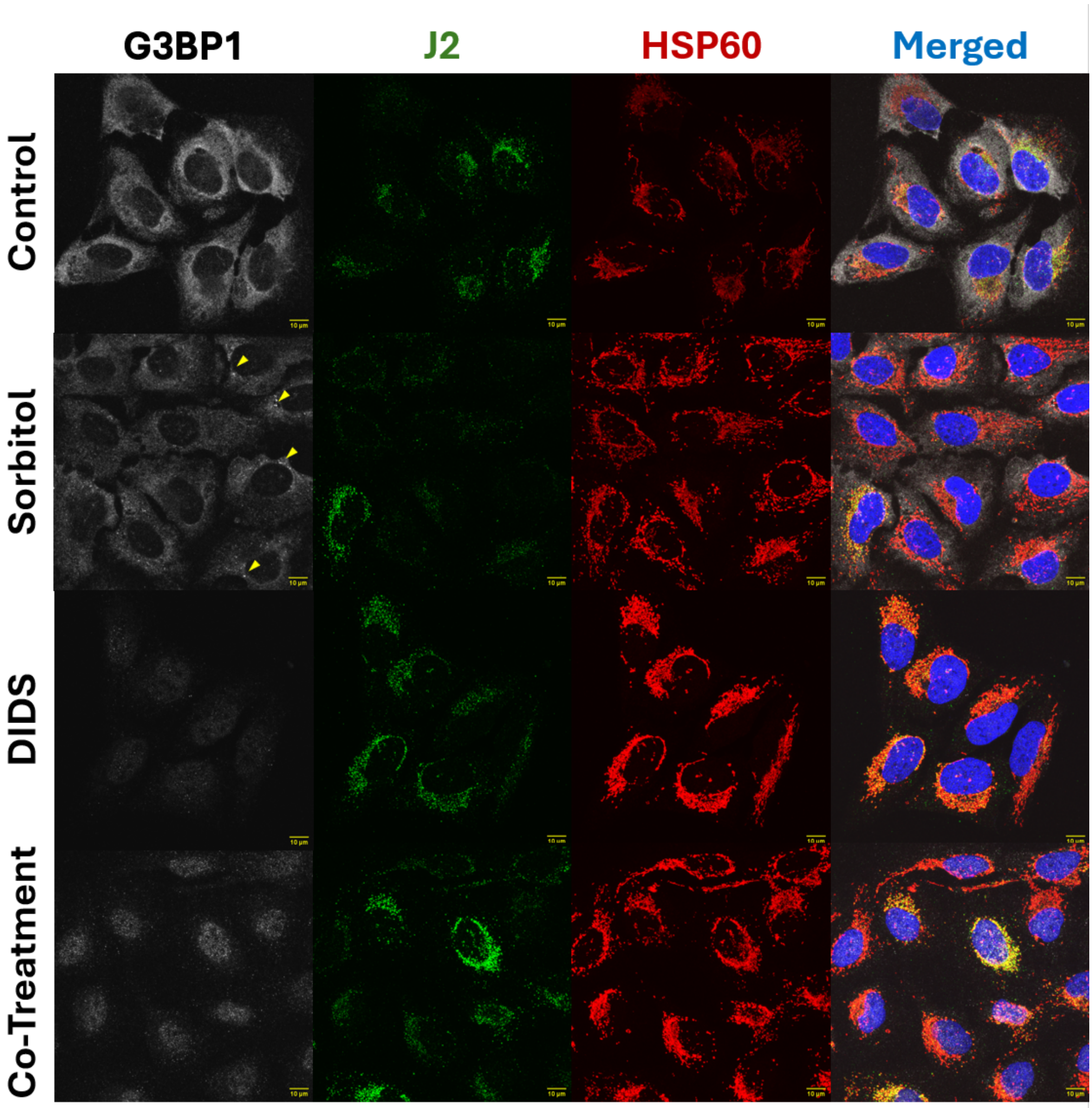


**Supplementary Figure 11. Blockage of mitochondrial permeability is compatible with SG assembly inhibition.** Representative Immunofluorescence images of U2-OS cells untreated or treated with sorbitol (0.4 M), DIDS (50 μM), or both. G3BP1 (SG marker, v. Figure 1B) is shown in white, J2 in green (dsRNA marker(21)), HPS60 in red (mitochondrial marker(22)), and nuclei (DAPI) in blue. Merged images are also provided. Yellow arrows in sorbitol-treated samples on the G3BP1 column highlight representative SGs. The Scale bar is shown on bottom right corner of each image, in yellow (10 μm).

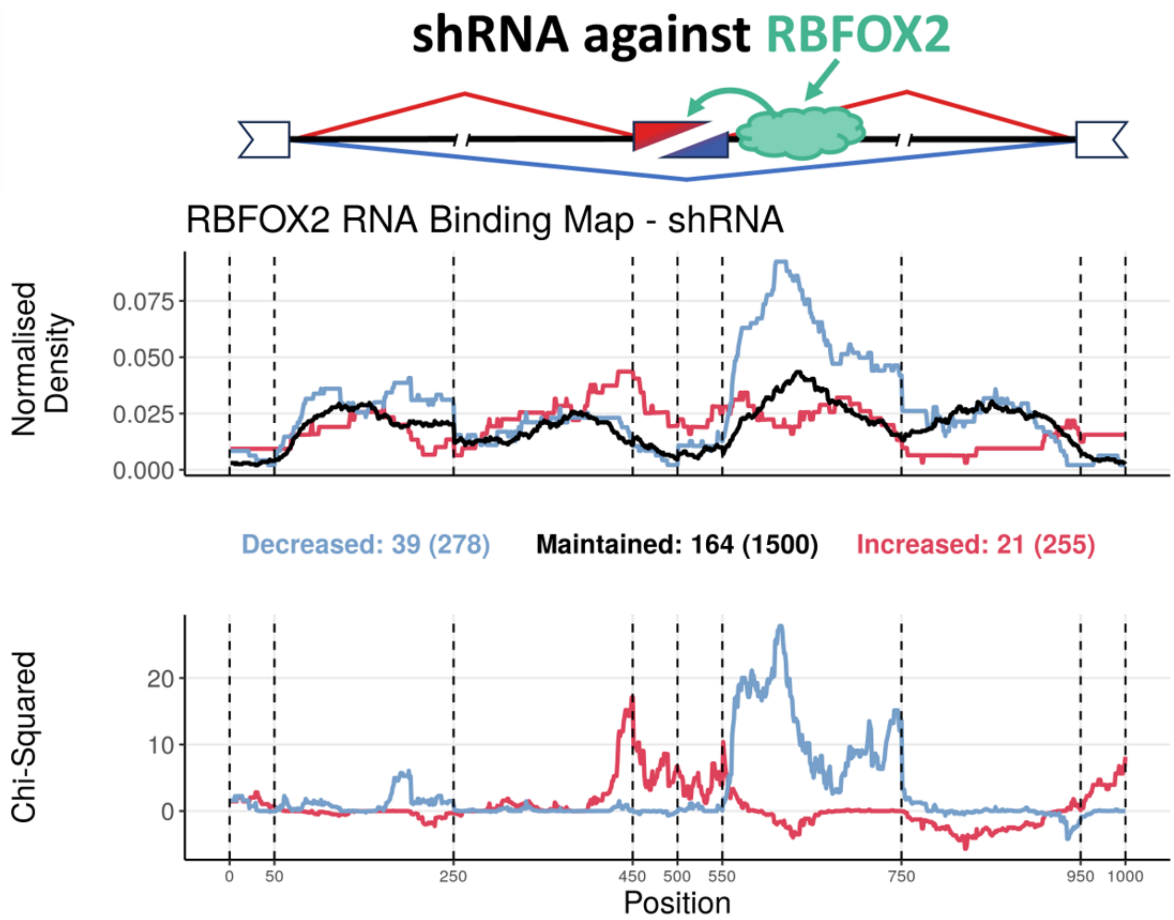


**Supplementary Figure 12. RNA binding map of RBFOX2 in HepG2 and K562 cells exposed to shRNA against RBFOX2.** (Top) Schematic of an alternative exon skipping event, regulated by an RBP, RBFOX2. (Middle) Normalised binding of RBFOX2 to metagenomic regulatory sequences of alternative exons. The plot discriminates excluded (blue), included (red) or maintained (black) alternative exons upon RBFOX2 knockdown, obtained from ENCODE(23, 24). The total number of events in each category can be found in brackets beneath the plot, preceded by the number of events for which there is at least one binding site for that RBP. (Bottom) Chi-Squared Statistic of excluded (blue) or included (red) normalised binding, in relation to maintained events. Vertical lines delimit the following metagenomic splicing regulatory sequences, in order: last 50 nt of upstream constitutive exon; first 200 nt of upstream intron; last 200 nt of upstream intron; first 50 nt of alternative exon; last 50 nt of alternative exon; first 200 nt of downstream intron; last 200 nt of downstream intron; first 50 nt of downstream constitutive exon. This example shows normalised RBFOX2 binding enriched downstream of the alternative exon. Upon RBFOX2 knockdown, this binding is therefore lost, consistent with increased exon skipping. These results align with the known role of RBFOX2 in promoting exon inclusion(25), providing a proof of concept for our approach.

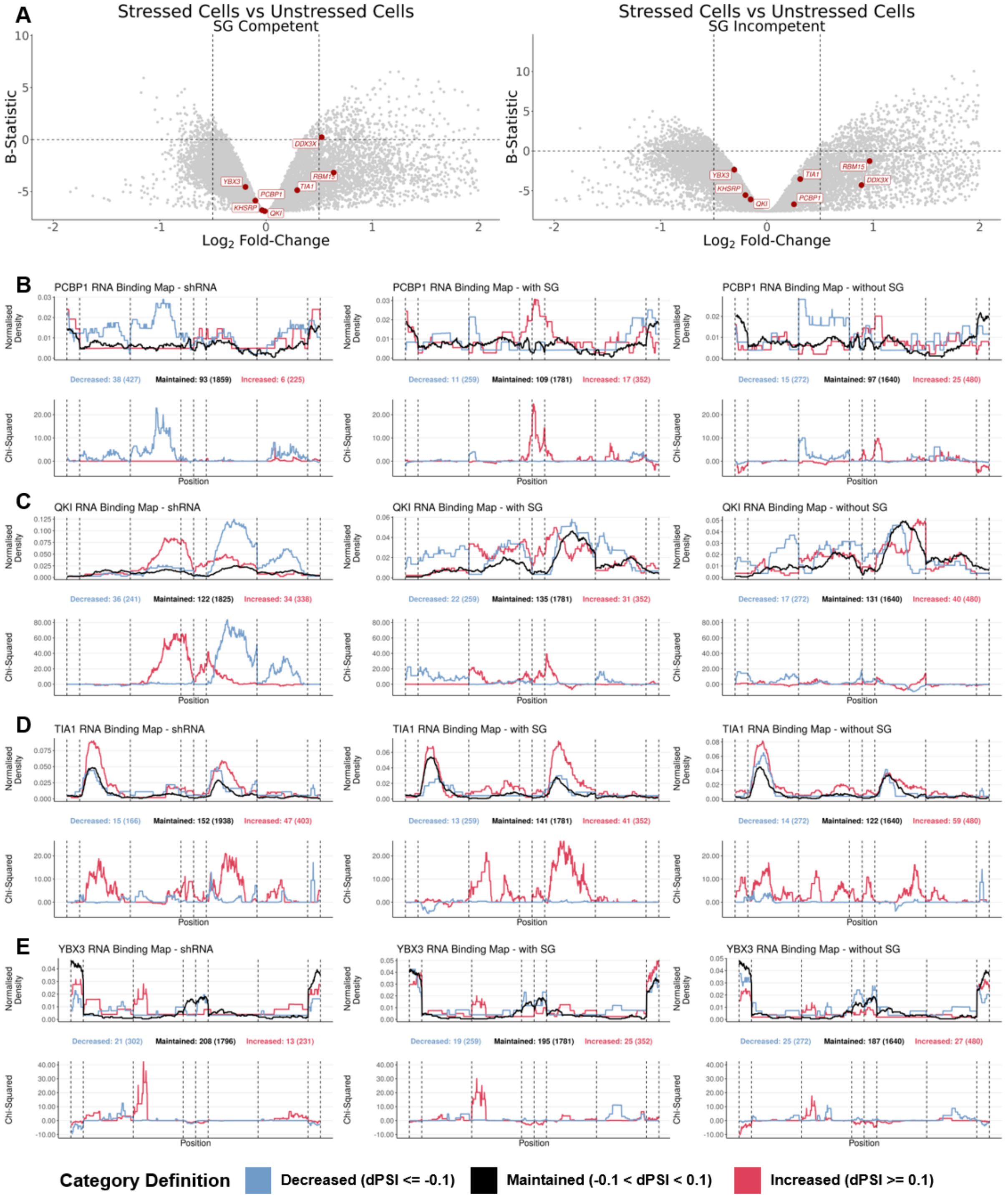


**Supplementary Figure 13. Alternative splicing alterations in SG competent and incompetent cells** (A) Volcano plot of DGEA between stressed and unstressed WCs (X-Axis – $Log_2$ Fold-Change; Y-Axis – B-Statistic) in SG competent (Left) and incompetent (Right) U2-OS and A549 cell lines subjected to dsRNA as a stressor(26); genes as circles, with RBPs with significant enrichment or depletion targets amongst differentially spliced sequences labelled and in red. Vertical dashed lines correspond to -0.5 and 0.5 $log_2$FC. Horizontal dashed line corresponds to B-statistic = 0. Only genes with -2 < log2FC < 2 are plotted. (B-E) RNA binding maps of alternative exon splicing events in shRNA-treated cells for PCBP1 (B), QKI (C), TIA1 (D), and YBX3 (E) obtained from ENCODE(23) (Left), in SG-competent cells (Middle) and in SG-incompetent cells (Right). (Top) Normalised binding of the RBP of interest to metagenomic regulatory sequences of alternative exons. The plot discriminates excluded (blue), included (red) or maintained (black) alternative exons upon RBP knockdown (Left) or upon dsRNA stress stimuli, in SG competent (Middle) and incompetent (Right) cells. The total number of events in each category can be found in brackets beneath the plot, preceded by the number of events for which there is at least one binding site for that RBP. (Bottom) Chi-Squared Statistic of excluded (blue) or included (red) normalised binding, in relation to maintained events. Vertical lines delimit the following metagenomic splicing regulatory sequences, in order: last 50 nt of upstream constitutive exon; first 200 nt of upstream intron; last 200 nt of upstream intron; first 50 nt of alternative exon; last 50 nt of alternative exon; first 200 nt of downstream intron; last 200 nt of downstream intron; first 50 nt of downstream constitutive exon.

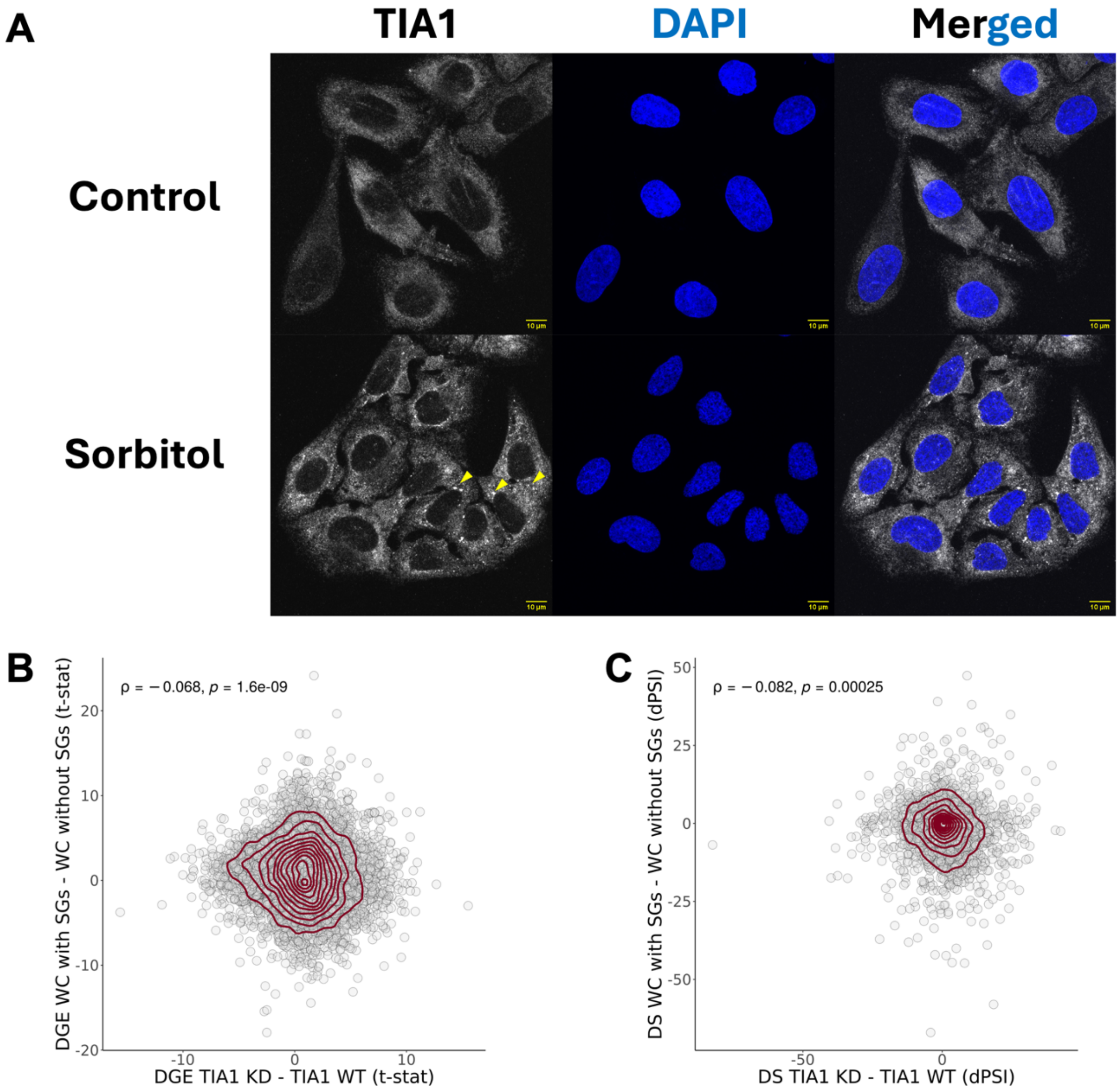


**Supplementary Figure 14. Alterations of TIA1 function upon KD and SG assembly.** (A) Immunofluorescence images of U2-OS cells untreated or treated with sorbitol (0.4 M). TIA1 is shown in white and nuclei (DAPI) in blue. Yellow arrows highlight putative SGs. The scale bar is shown on bottom right corner of each image, in yellow (10 µm). (B) Scatter plot of t-statistics of DGE between HEPG2 and K562 cells treated with shRNA against TIA1 and those with untargeted shRNA(23, 24) (X-Axis) and between U2-OS and A549 WT Cells (SG-Competent) exposed to dsRNA as a stressor against their G3BP1/G3BP2 KO counterparts (SG-Incompetent)(26) (Y-Axis); genes as circles; density contour lines in dark red, Spearman's correlation coefficient ρ and associated test's p-value on top left corner. (C) Scatter plot of differential splicing (in difference in percent spliced-in (dPSI)) between HEPG2 and K562 cells treated with shRNA against TIA1 and those with untargeted shRNA(23, 24) (X-Axis) and between U2-OS and A549 WT Cells (SG-Competent) exposed to dsRNA as a stressor and their G3BP1/G3BP2 KO counterparts (SG-Incompetent)(26) (Y-Axis); alternative splicing events as circles; density contour lines in dark red, Spearman's correlation coefficient ρ and associated test's p-value on top left corner.

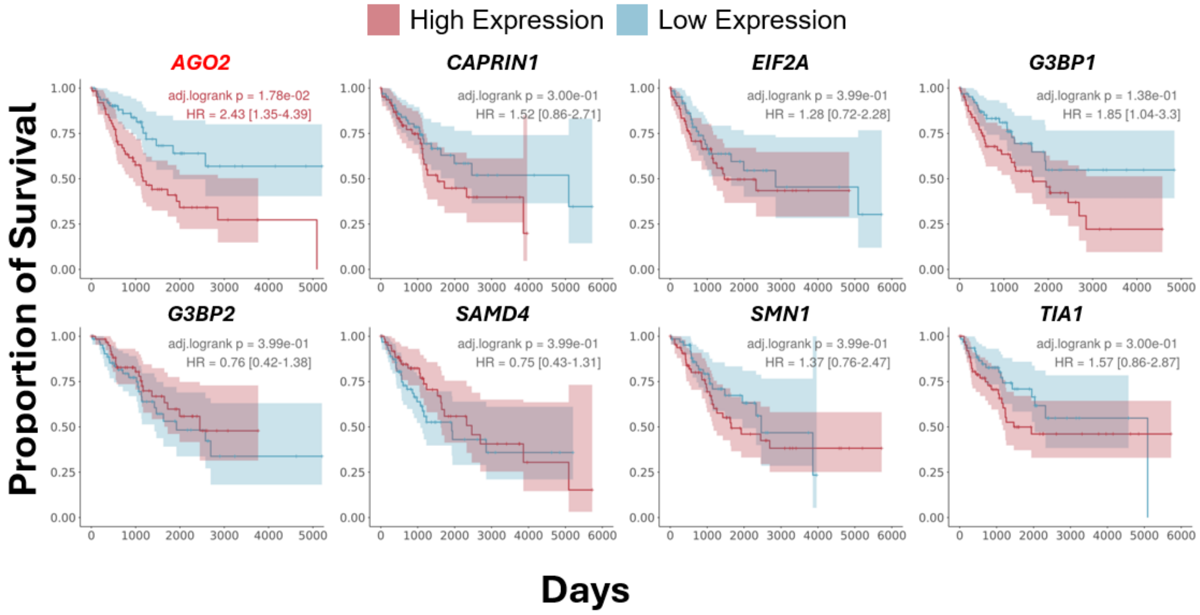


**Supplementary Figure 15. Overall survival analysis based on the expression of genes encoding SG-related proteins.** Kaplan–Meier plots showing proportion of overall survival (Y-Axis) per day (X-Axis) of patients from GDC-TCGA's SARC dataset (The Cancer Genome Atlas sarcoma)(27), stratified by expression of the indicated genes, as plot titles. Patients were grouped into high and low expression cohorts based on the bottom and top quartiles of expression values. BH-adjusted Log-rank p-values, and HR, are shown in the top right corner of each plot. Genes with an adjusted Log-rank p-value <0.05 are shown in red.

A

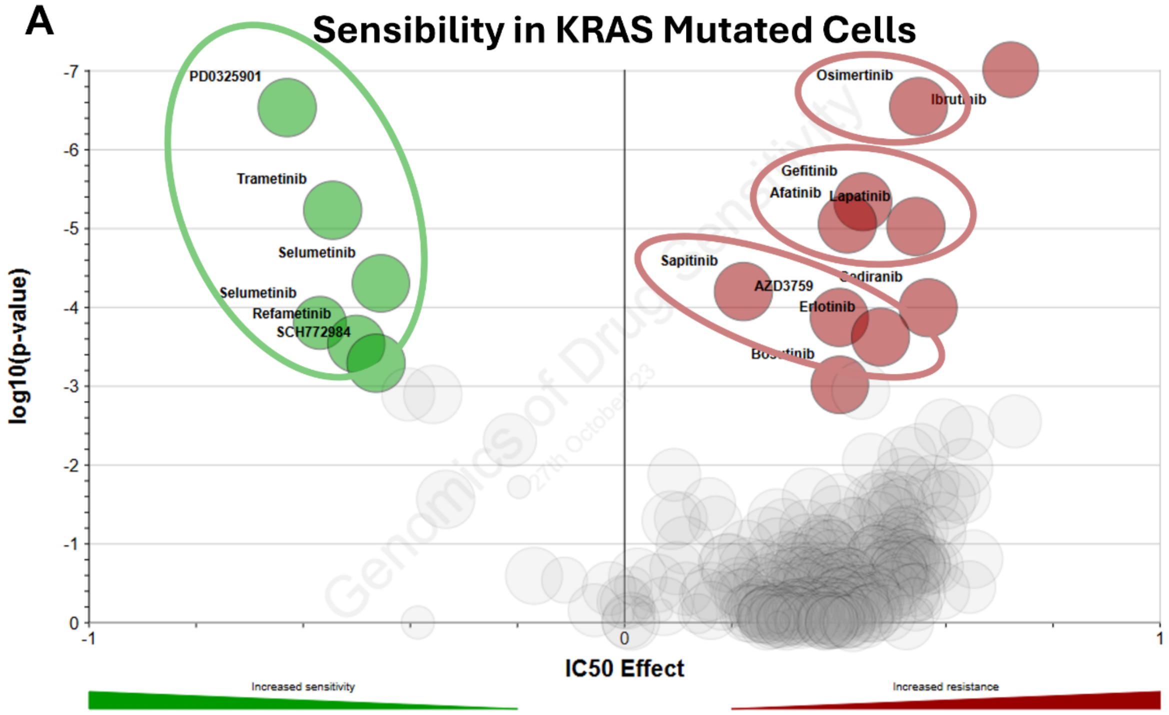


B

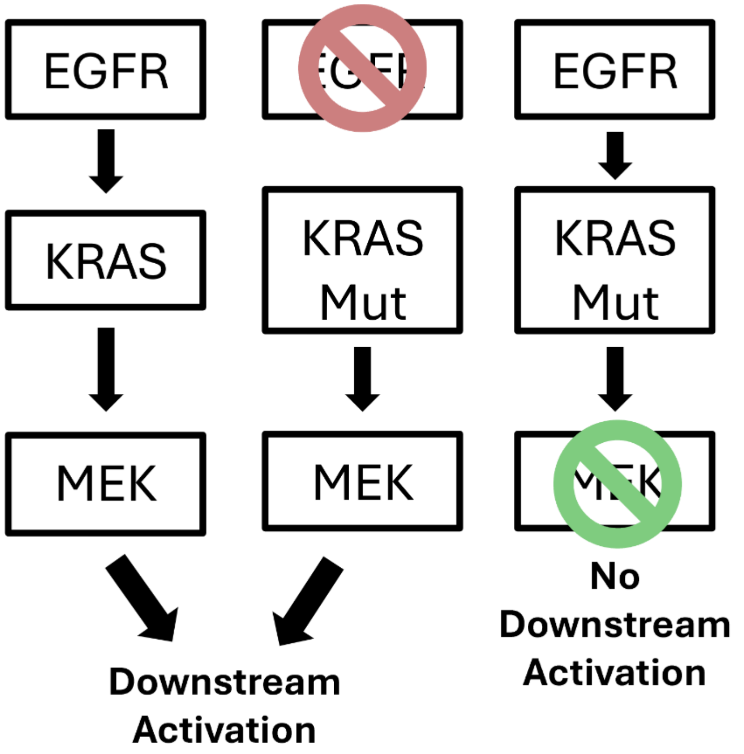


C

**Most Similar Perturbations to PTGS2 KD**

| Score | Type | Name |
|---|---|---|
| 95.16 | KD | AKT3 |
| 92.57 | KD | EGFR |
| 92.57 | KD | ERBB3 |
| 92.54 | KD | GSK3A |
| 90.77 | KD | TP53RK |
| 90.63 | KD | NUDT5 |
| 90.14 | KD | C9orf96 |
| 90.07 | KD | CDK6 |
| 89.53 | KD | PIK3CB |
| 89.45 | KD | ZNF671 |

**Supplementary Figure 16. KRAS mutation and associated drug sensitivities and resistances.** (A) Volcano plot of half maximal inhibitory concentration (X-Axis – IC50 Effect) and corresponding significance, given by MANOVA(28) (Y-Axis – $\log_{10}$(p-value)) in KRAS mutant cells; drugs/chemicals as circles, with drugs with the most differential IC50 effect labelled and coloured in green (if with increased sensitivity) or in red (if with increased resistance). Green circle around drugs which target MEK, and red circles around drugs that target EGFR. Analysis and images are obtained from GDSC(28), using the GDSC2 dataset. (B) Simplified schematic of the EGFR signalling pathway. (Left) In wild-type cells, EGFR activation triggers KRAS activation, which in turn activates MEK and downstream signalling. (Middle) In KRAS-mutant cells, inhibition of EGFR with targeted drugs (red drugs, see (A)) does not block signalling, as mutant KRAS activates MEK independently. This explains resistance of KRAS-mutant cells to anti-EGFR therapy. (Right) In KRAS-mutant cells, inhibition of MEK with targeted drugs (green drugs, see (A)) blocks downstream signalling, rendering these cells sensitive to anti-MEK therapy. (C) Table representing alterations with the highest correlation to *PTGS2* KD. Values represent CMap connectivity scores, which reflect the degree of similarity between gene expression signatures. Knockdowns of *EGFR* and *ERBB3* are highlighted in red. Analysis and visualisation were performed and adapted from the Connectivity Map (CMap) platform(29).

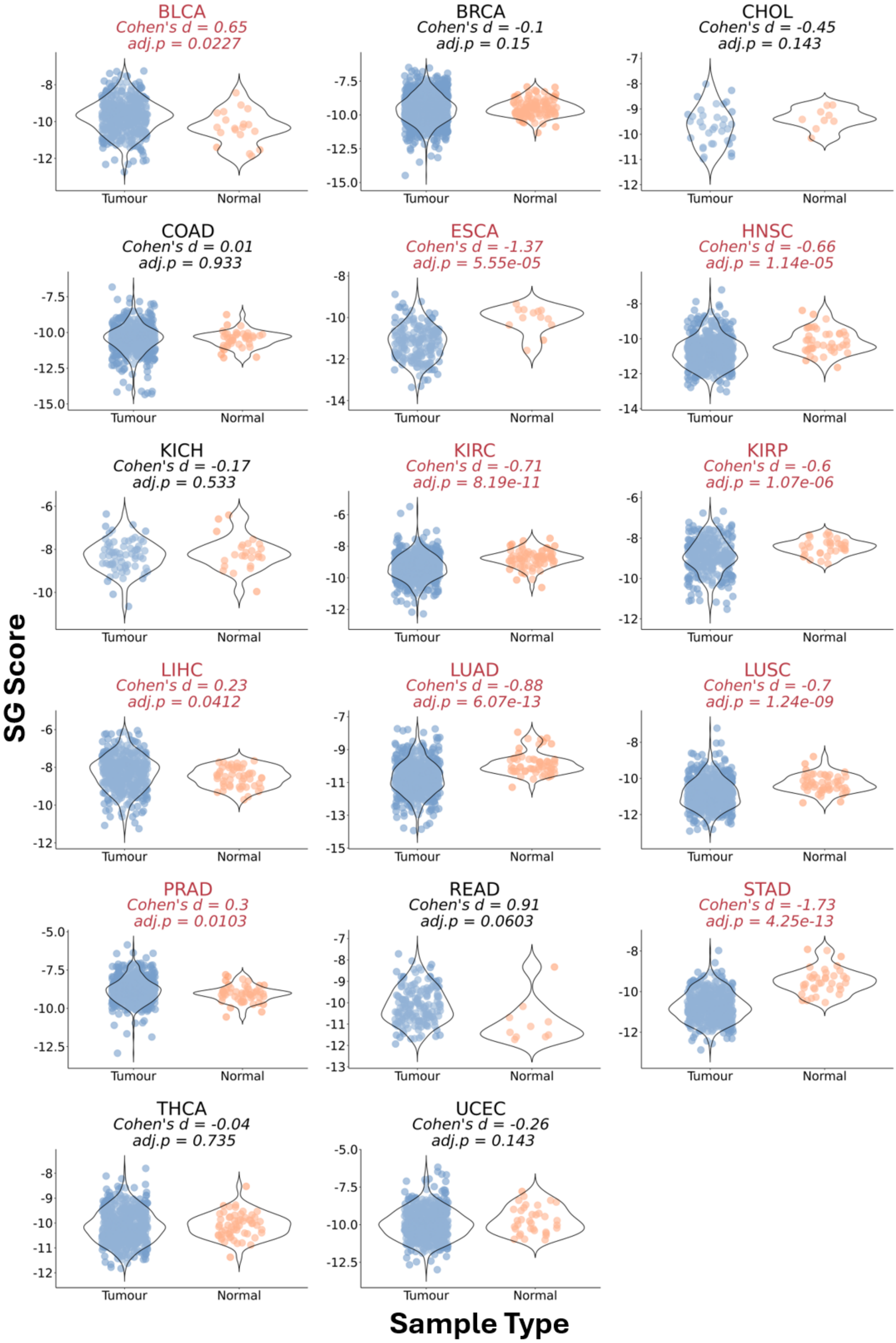


**Supplementary Figure 17. Signature score in various GDC-TCGA datasets.** Violin plots of SG signature scores, calculated through a ranking metric (see Methods), in tumour (blue) and normal tissue (orange) samples from several GDC-TCGA datasets(27, 30). Effect Size (Cohen's d) and corresponding statistical significance given by Welch's t-test (adjusted p-value) are shown above the plots. Only GDC-TCGA(27, 30) datasets with at least 5 normal and tumour samples were considered for this analysis.

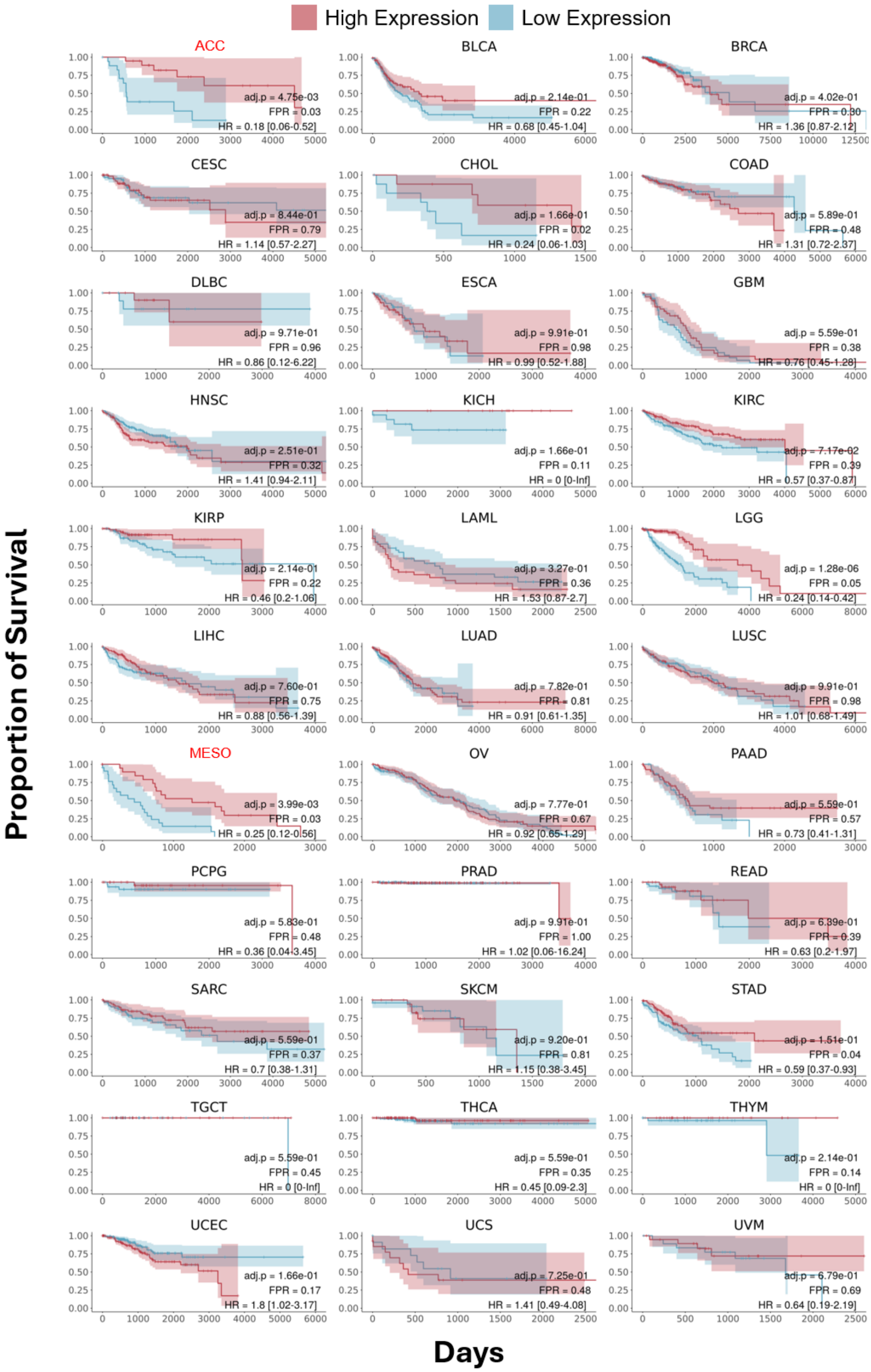
High Expression
Low Expression
Proportion of Survival
Days
ACC
adj.p = 4.75e-03
FPR = 0.03
HR = 0.18 [0.06-0.52]
BLCA
adj.p = 2.14e-01
FPR = 0.22
HR = 0.68 [0.45-1.04]
BRCA
adj.p = 4.02e-01
FPR = 0.30
HR = 1.36 [0.87-2.12]
CESC
adj.p = 8.44e-01
FPR = 0.79
HR = 1.14 [0.57-2.27]
CHOL
adj.p = 1.66e-01
FPR = 0.02
HR = 0.24 [0.06-1.03]
COAD
adj.p = 5.89e-01
FPR = 0.48
HR = 1.31 [0.72-2.37]
DLBC
adj.p = 9.71e-01
FPR = 0.96
HR = 0.86 [0.12-6.22]
ESCA
adj.p = 9.91e-01
FPR = 0.98
HR = 0.99 [0.52-1.88]
GBM
adj.p = 5.59e-01
FPR = 0.38
HR = 0.76 [0.45-1.28]
HNSC
adj.p = 2.51e-01
FPR = 0.32
HR = 1.41 [0.94-2.11]
KICH
adj.p = 1.66e-01
FPR = 0.11
HR = 0 [0-Inf]
KIRC
adj.p = 7.17e-02
FPR = 0.39
HR = 0.57 [0.37-0.87]
KIRP
adj.p = 2.14e-01
FPR = 0.22
HR = 0.46 [0.2-1.06]
LAML
adj.p = 3.27e-01
FPR = 0.36
HR = 1.53 [0.87-2.7]
LGG
adj.p = 1.28e-06
FPR = 0.05
HR = 0.24 [0.14-0.42]
LIHC
adj.p = 7.60e-01
FPR = 0.75
HR = 0.88 [0.56-1.39]
LUAD
adj.p = 7.82e-01
FPR = 0.81
HR = 0.91 [0.61-1.35]
LUSC
adj.p = 9.91e-01
FPR = 0.98
HR = 1.01 [0.68-1.49]
MESO
adj.p = 3.99e-03
FPR = 0.03
HR = 0.25 [0.12-0.56]
OV
adj.p = 7.77e-01
FPR = 0.67
HR = 0.92 [0.65-1.29]
PAAD
adj.p = 5.59e-01
FPR = 0.57
HR = 0.73 [0.41-1.31]
PCPG
adj.p = 5.83e-01
FPR = 0.48
HR = 0.36 [0.04-3.45]
PRAD
adj.p = 9.91e-01
FPR = 1.00
HR = 1.02 [0.06-16.24]
READ
adj.p = 6.39e-01
FPR = 0.39
HR = 0.63 [0.2-1.97]
SARC
adj.p = 5.59e-01
FPR = 0.37
HR = 0.7 [0.38-1.31]
SKCM
adj.p = 9.20e-01
FPR = 0.81
HR = 1.15 [0.38-3.45]
STAD
adj.p = 1.51e-01
FPR = 0.04
HR = 0.59 [0.37-0.93]
TGCT
adj.p = 5.59e-01
FPR = 0.45
HR = 0 [0-Inf]
THCA
adj.p = 5.59e-01
FPR = 0.35
HR = 0.45 [0.09-2.3]
THYM
adj.p = 2.14e-01
FPR = 0.14
HR = 0 [0-Inf]
UCEC
adj.p = 1.66e-01
FPR = 0.17
HR = 1.8 [1.02-3.17]
UCS
adj.p = 7.25e-01
FPR = 0.48
HR = 1.41 [0.49-4.08]
UVM
adj.p = 6.79e-01
FPR = 0.69
HR = 0.64 [0.19-2.19]

**Supplementary Figure 18.** Kaplan–Meier plots showing proportion of overall survival (Y-Axis) per days (X-Axis) of patients of GDC-TCGA datasets(27, 30), stratified by SG score, with dataset identification as plot title. Patients were grouped into high and low expression cohorts based on bottom and upper quartile SG Score values. BH-adjusted Log-rank p-value, HR, and FPR compared to 100 random signatures are shown on the bottom right of each plot; datasets with FPR > 0.05 are highlighted in red.

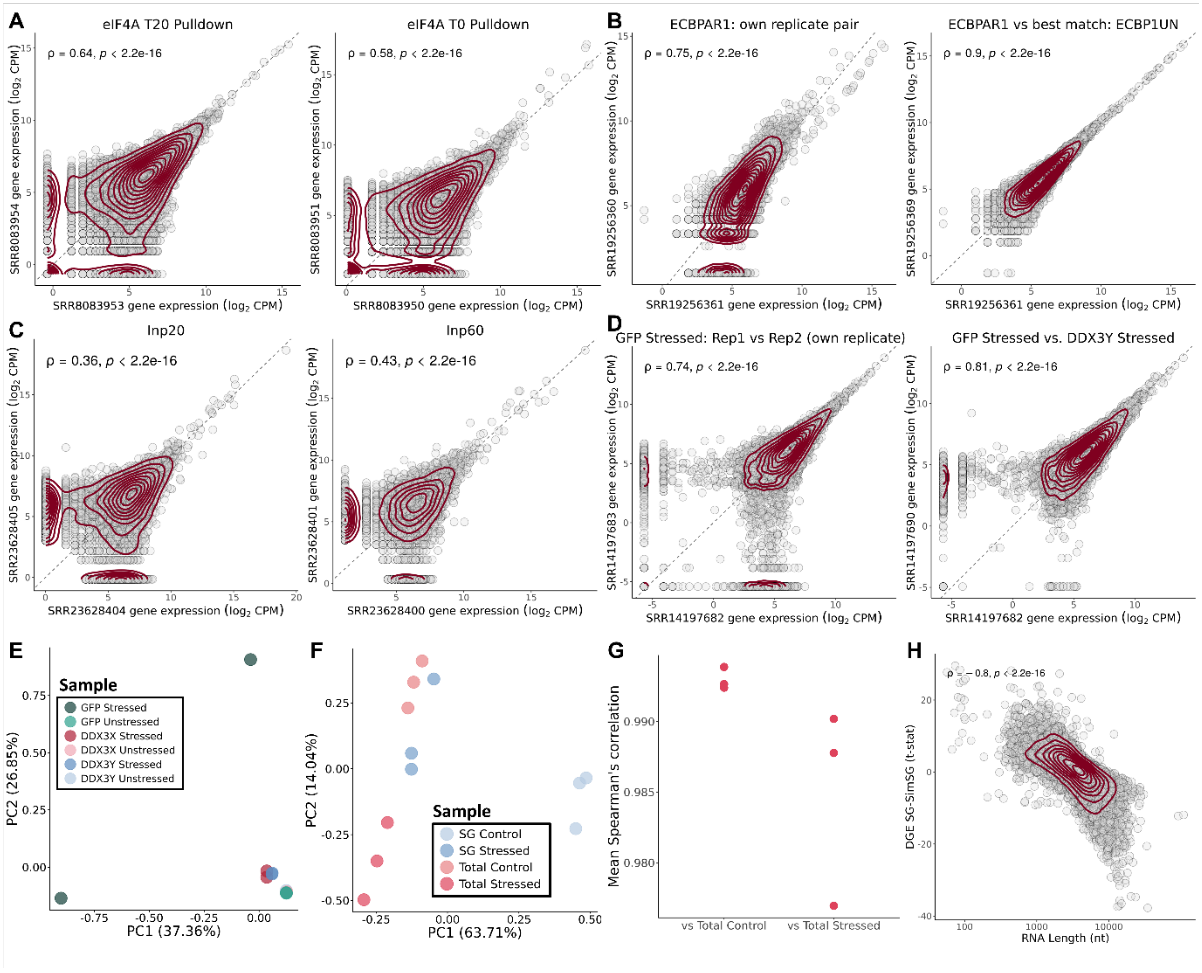


**Supplementary Figure 19. Exclusion criteria applied to publicly available SG transcriptome datasets.** (A) Scatter plots comparing expression ($log_2$CPM) in Padron 2019's dataset(4) between the two replicates of eIF4A T20 (Left) and eIF4A T0 (Right) pulldown conditions. (B) Scatter plots comparing gene expression ($log_2$CPM) in Sato 2023's dataset(6) between the two declared replicates of the ECBPAR1 (stressed G3BP1 interactome) pulldown condition (own replicate pair) (Left) and between one ECBPAR1 replicate and its best-correlating alternative sample, ECBP1UN (unstressed G3BP1 interactome), defined as the sample with the highest Spearman correlation excluding its declared replicate (Right). (C) Scatter plots comparing gene expression ($log_2$CPM) in Xiao 2024's dataset(7) between the two replicates of the Input 20 (WC samples after 20 minutes of arsenite stress) (Left) and Input 60 (WC samples after 60 minutes of arsenite stress) (Right) conditions. (D) Scatter plots of gene expression ($log_2$CPM) in Shen 2022's dataset(9) between the two declared replicates of the GFP Stressed condition (WC stressed samples) (own replicate pair) (Left) and between one GFP Stressed replicate and its best-correlating alternative sample (DDX3Y Stressed, putative SG) (Right). (E) PCA plot of gene expression of all samples from Shen 2022's dataset(9), coloured by sample type; note the separation between the two GFP Stressed replicates relative to all other replicate pairs. (F) PCA plot of gene expression of all samples from Iadevaia 2022's dataset(12), coloured by sample type. (G) Mean Spearman’s correlation in gene expression between SG Stressed samples and Total Control and Total Stressed samples, from Iadevaia 2022's dataset(12); each dot represents one SG Stressed replicate's mean correlation with all samples in the indicated group. (H) Scatter plot of t-statistics of DGE (Y-Axis) between the observed SG transcriptome and simulated SGs, and RNA transcript length (X-Axis), from

Iadevaia 2022's dataset(12). In (A, B, C, D, H), transcripts as circles, density contour lines in dark red, Spearman's correlation coefficient ρ and associated test's p-value on top left corner. In (A, B, C, D) the dashed line depicts identity.
